# MarkovHC: Markov hierarchical clustering for the topological structure of high-dimensional single-cell omics data

**DOI:** 10.1101/2020.11.04.368043

**Authors:** Zhenyi Wang, Yanjie Zhong, Zhaofeng Ye, Lang Zeng, Yang Chen, Minglei Shi, Minping Qian, Michael Q. Zhang

**Affiliations:** MOE Key Laboratory of Bioinformatics; Bioinformatics Division and Center for Synthetic & Systems Biology, BNRist; Department of Automation, Tsinghua University, Beijing 100084, China; MOE Key Laboratory of Bioinformatics; Bioinformatics Division and Center for Synthetic & Systems Biology, BNRist; School of Medicine, Tsinghua University, Beijing 100084, China; School of Mathematical Sciences, Peking University, Beijing 100871, China; Department of Biological Sciences, Center for Systems Biology, The University of Texas, Richardson, TX 75080-3021, USA; Department of Mathematics and Statistics, Washington University in St. Louis, St. Louis, MO 63130, USA; Department of Biostatistics, University of Michigan, Ann Arbor, MI 48109, USA

## Abstract

Distinguishing cell types and cell states is one of the fundamental questions in single-cell studies. Meanwhile, exploring the lineage relations among cells and finding the path and critical points in the cell fate transition are also of great importance.

Existing unsupervised clustering methods and lineage trajectory reconstruction methods often face several challenges such as clustering data of arbitrary shapes, tracking precise trajectories and identifying critical points. Certain adaptive landscape approach^1–3^, which constructs a pseudo-energy landscape of the dynamical system, may be used to explore such problems. Thus, we propose Markov hierarchical clustering algorithm (MarkovHC), which reconstructs multi-scale pseudo-energy landscape by exploiting underlying metastability structure in an exponentially perturbed Markov chain^4^. A Markov process describes the random walk of a hypothetically traveling cell in the corresponding pseudo-energy landscape over possible gene expression states. Technically, MarkovHC integrates the tasks of cell classification, trajectory reconstruction, and critical point identification in a single theoretical framework consistent with topological data analysis (TDA)^5^.

In addition to the algorithm development and simulation tests, we also applied MarkovHC to diverse types of real biological data: single-cell RNA-Seq data, cytometry data, and single-cell ATAC-Seq data. Remarkably, when applying to single-cell RNA-Seq data of human ESC derived progenitor cells^6^, MarkovHC not only could successfully identify known cell types, but also discover new cell types and stages. In addition, when using MarkovHC to analyze single-cell RNA-Seq data of human preimplantation embryos in early development^7^, the hierarchical structure of the lineage trajectories was faithfully reconstituted. Furthermore, the critical points representing important stage transitions had also been identified by MarkovHC from early gastric cancer data^8^.

In summary, these results demonstrate that MarkovHC is a powerful tool based on rigorous metastability theory to explore hierarchical structures of biological data, to identify a cell sub-population (basin) and a critical point (stage transition), and to track a lineage trajectory (differentiation path).

**Highlights:**

1. MarkovHC explores the topology hierarchy in high-dimensional data.
2. MarkovHC can find clusters (basins) and cores (attractors) of clusters in different scales.
3. The trajectory of state transition (transition paths) and critical points in the process of state transition (critical points) among clusters can be tracked.
4. MarkovHC can be applied on diverse types of single-cell omics data.

## Introduction

Cellular activities can be modeled as stochastic dynamical states in the high-dimensional space of expression levels of genes, proteins, or other molecular quantities^9^. Modeling the dynamics of these processes can greatly improve our understanding of the cell fate decision, carcinogenesis, and early diagnoses of diseases. S. Wright^10^ first proposed the adaptive landscape concept in 1932 to describe population dynamics. Waddington’s epigenetic landscape^11,12^ was proposed later and has shown increasing application in biology^13,14^.

Nowadays, high throughput single-cell technologies provide a tremendous amount of data for modeling cellular dynamics, however, the observed data suffer from a high level of noises due to both intrinsic biological stochasticity and technical errors in spite of their potential to overcome heterogeneity as shown, e.g. in exploring the lineage relations among differentiated cells^15–19^ and predicting cellular communications^20^. Computationally, clustering of cells and delineating density ridges/energy landscapes for trajectory reconstruction, are the two most important tasks, especially in topological data analysis (TDA)^5,21^. Existing methods remain facing a number of difficult challenges such as clustering non-convex and mixture-density data, detecting rare clusters, tracking precise trajectories and identifying critical points.

Density ridge, a basic idea in topological data analysis is the essence of clustering and trajectory problems but apparently ignored by a lot of existing methods. In contract, reconstructing the underlying landscape maintaining the topological structure can shed light upon these tasks and overcome most of the challenges mentioned above. Meanwhile, cells are known to be regulated by redundant genes, therefore the gene expression data observed in a high dimensional space can be embedded in lower-dimensional manifolds^22^. Different cell types express distinctive combination gene sets, which can be viewed as supports in mathematics. They shape different manifolds^23^ and form disconnected blocks if ignoring boundary points. In addition, the definition of cell types greatly depends on classification criteria and is often ambiguous. The hierarchical structure, however, uses a series of levels corresponding to cells clustering in different fineness degrees to overcome the hard threshold problem. Moreover, the hierarchical structure can integrate information concisely^24,25^ and reveal the inherent structure in many biological systems^26^. Thus, we developed a novel tool, MarkovHC, to address these issues in single-cell data analysis. MarkovHC is unsupervised (no assumption about the distribution of the data), which makes it intuitive, robust, and widely applicable. It recognizes cell clusters (basins) and cores of clusters (attractors) in different scales, tracks critical points (stage transition points) and transition paths (the trajectory of state transition) between clusters and packs all information in a hierarchical structure, which respects the topology of the data.

In this work, we compared MarkovHC with 19 popular clustering methods^19,25,27–46^ (in part C and part D of Supplementary Notes and sheet 1 of Supplementary Table 1) and 4 representative trajectory construction methods (part C of Supplementary Notes) including SPADE^15^, monocle series^16,47,48^, URD^49^ and Waddington-OT^14,26^ in general. These existing works may be divided into two classes: data-based and model-based. Since MarkovHC is unsupervised, it belongs to the first class. For a given set of samples, a Markov chain in sample state space is constructed (see Supplementary Notes) and its steady state (an invariant measure of the Markov dynamics) is obtained to approximating the probability density model with an adjustable coarse-graining scale (“temperature”) parameter. Firstly, to validate the effectiveness of MarkovHC and to show its advantages, we applied it to simulated data and several benchmark single-cell datasets. Then, we used MarkovHC to explore the hierarchical structure in mouse ES cell differentiation, the transition path and critical points in human embryo development, and the critical points in gastric cancer. These results demonstrate that MarkovHC is capable of exploring hierarchical structure in high-dimensional data, finding basins and attractors in different scales, tracking critical points, and transition paths among basins systematically. In summary, MarkovHC is a powerful and extendable method based on rigorous mathematical theory for studying the landscape of single-cell data and shows great potential in analyzing extensive types of single-cell omics data.

## Results

### The MarkovHC algorithm

In this section, we briefly introduce the proposed algorithm. The general flowchart of MarkovHC is shown in Fig. 1.

**Figure 1.**
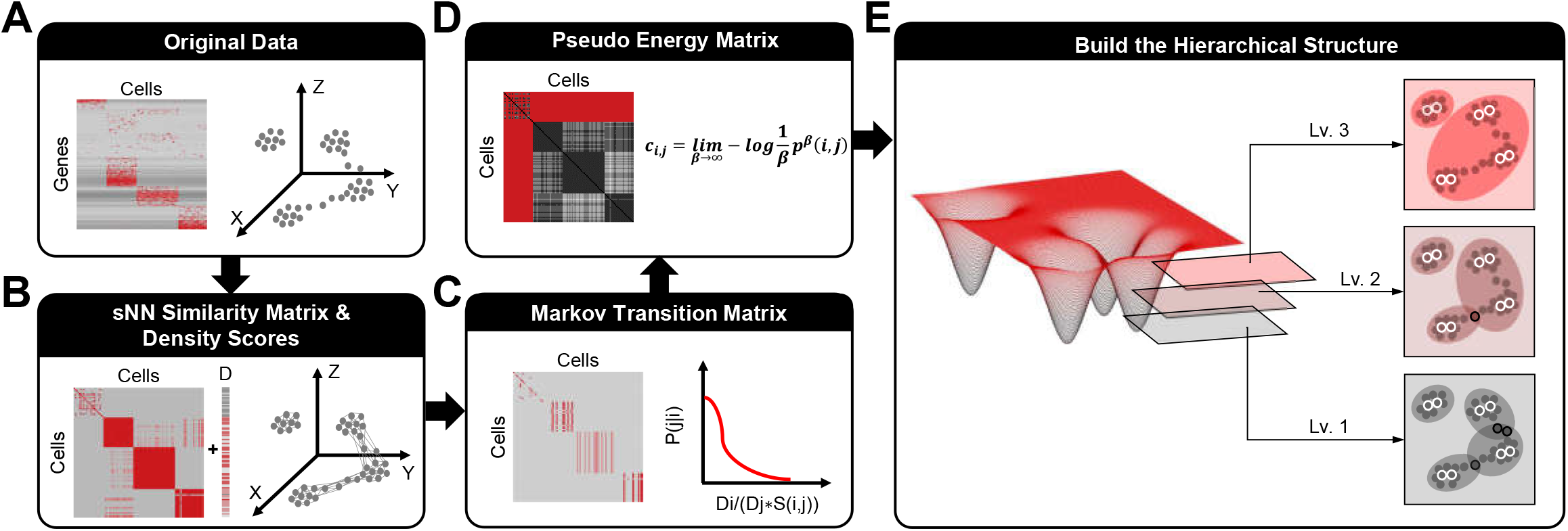
Overview of MarkovHC. (A) The original input data is the matrix of genes by cells. (B) We calculate sNN (shared Nearest Neighbors) among cells to get the cell by cell similarity matrix. Then we construct a cellular network using the similarity matrix and calculate each cell’s degree (D score) in the network. (C) The Markov transition matrix is calculated using the similarity matrix and D scores. (D) The pseudo-energy matrix is calculated based on the Markov transition matrix. (see Methods) (E) The hierarchical structure is constructed based on attractors, basins, and critical points on each level.

A gene expression (or flow cytometry) data matrix (A={agc: g=1,2,…,G, c=1,2,…,C} is taken as a generic input (Fig. 1A). In principle, treating each distinct cell as a unique point (vertex) or vector in the G-dimensional “expression” space, an initial (i.e. lowest level) Markov network can be built by defining a transition probability matrix from vertex “density” (i.e. higher local vertex weight means more cells could jump out per unit time) and edge similarity (i.e. more interaction edge strength means more “flow conductance”). The “density” may be defined as the number of cells in the same state (same expression level) or vertex degree, and the similarities among states may be calculated using the cells shared between nearest neighbor (sNN) states (Fig. 1B). Since calculating the global Euclidean distance directly in high-dimensional space tends to introduce noises and biases, sNN is used instead, which is able to explore the natural (“geodesic”) distance on the underlying “manifolds” and has been proved to get more robust results^50^ in recovering cell subpopulations^51,52^ and reconstructing trajectories^23,50,53,54^. In most graphical models, a similarity matrix alone is sufficient to define a Markov transition probability matrix. Since we prefer a pre-processing step (see Methods for details) for robustness, the degree of each node in the sNN network was used to measure the point aggregation density, (see Methods for the density score definition). Hence, the density scores of the nodes were used together with the similarity matrix to define the basic Markov transition probability. (Fig. 1C)

Given the transition probability matrix on each level, MarkovHC can find attractors and basins on each level and construct the hierarchical structure systematically. In this work, we term basins as clusters and attractors as cores of these clusters. Attractors are steady states in multi-stable systems and basins consist of attractors and their subsidiary points. These two notions have been widely used in previous works^55,56^ modeling the nonlinear dynamics of organisms and biological phenomena as dynamics systems.

Based on the metastability theory developed by one of us (Chen et.al. 1996)^4^, the hierarchical structure is built in an iterative manner. Firstly, MarkovHC finds the attractors and basins on the first level (details are in part A of Supplementary Notes). Secondly, for each point in attractors, Dijkstra shortest path algorithm^57^ is deployed to detect the shortest transition path from it to its nearest neighboring basin using the pseudo-energy matrix (Fig. 1D). The length of the shortest path represents the pseudo-energy cost in the transition from one basin (cluster) to its nearest neighboring basin (cluster). On the second level, state-space consists of basins on the first level. To form the Markov transition matrix on the second level, the transition probabilities among basins are calculated. Then, MarkovHC uses the same approach as that on the first level to find attractors and basins on the second level. Repeating this process and we can obtain attractors and basins on each level as well as the relations among basins on neighboring levels. The hierarchical structure of the levels (Fig. 1E) is built using these attractors, basins, and topological relations.

On each transition energy level, we can identify the transition path and critical points. The transition path is defined as the most probable transiting path between two basins and the critical points are defined as the closest points between two basins on the transition path. In previous works^58^, the saddle points in stochastic nonlinear dynamics were regarded as critical points. However, in practice, it’s impossible to locate perfect saddle points with discrete data. Hence, we regard boundary points on the transition path as critical points to approximate saddle points.

### MarkovHC can characterize clusters, cores, transition paths and critical points among clusters

In this section, we use simple examples to intuitively show how MarkovHC can find basins, attractors, critical points, transition paths among basins, and constructing the intrinsic hierarchical structure.

Although a large number of clustering methods have been developed, the definition of a cluster is still ambiguous. Inspired by topological data analysis (TDA) ^21^, MarkovHC algorithm is based on the topological connectivity among points, which gives us a clear and reasonable definition of a cluster. Wasserman^21^ proposed a concept called level sets for targeted density function. Here, we generalize this idea to discrete data clustering problems. For any given δ> 0, points with similarities smaller than δ are partitioned into one cluster. The larger δ is used, the larger and fewer clusters are obtained. As δ increases and reaches some critical value, some small clusters will merge into a larger one and form hierarchical structures. The above definition of a cluster is clear but lacked biological application because the interaction of two states in a biological system can often be asymmetric (no detailed balance or time-reversal symmetry). Therefore, we modified the symmetric δ connectivity to a directed graph. For any fixed δ, we can obtain a directed graph, from which we can tell basins, attractors, critical points, and transition paths. We used intuitive plots (Fig. 2A) to illustrate how MarkovHC builds up a topologically structure-based hierarchy. Note that when we mention the length of a directed line segment below, we refer to some “asymmetric distance” of biological significance, for example, the amount of transition energy consumption. With a given δ, points that can be mutually connected by directed line segments consist of attractors (white border, Fig. 2A), which are the representative points in the highest density regions of basins. Other points that can link to a given attractor are assigned to the corresponding basin. As δ increasing, according to the order of basins’ immergence, the hierarchical structure of these points can be constructed.

**Figure 2.**
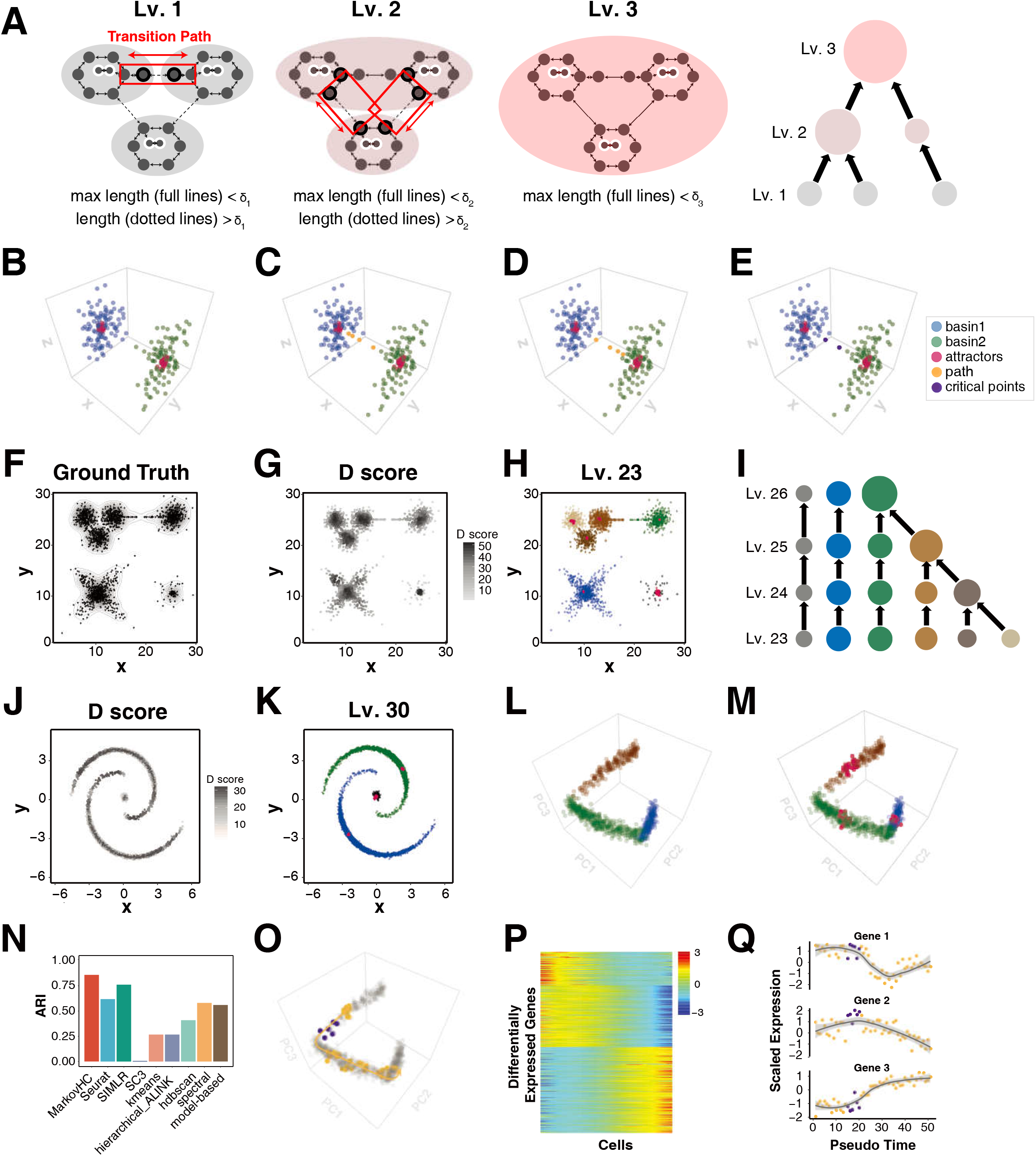
MarkovHC is able to cluster non-convex data, hetero-density data, and differentiation stages in a continuum. (A)The points that can be connected using lines (pseudo-energy) no longer than δn are partitioned into a basin on level n. In a basin, points in the highest density regions constitute attractors (points with white border). Points connecting basins’ attractors constitute the transition path (red box), and the closest points (black border) between these two basins on the transition path are the critical points. (B-E)These two group points are clustered into two basins (blue and green) by MarkovHC (B). To insight more details of these two basins, we highlighted the attractors (red), the transition paths (yellow) and critical points (purple) respectively. The points linking basin A’s attractors and basin B’s border constitute the transition path from A to B and vice versa (C, D). The transition path between A and B are the points linking their attractors, and the critical points which indicate the points jump at the boundary of these two basins are the two closest points belong to them on the transition path (E). (F) The 2-dimensional simulated data with density contours. (G) D score of each point. (H) The basins and attractors on level 23 were plotted to show the 2-dimensional layouts. And we observed that MarkovHC can simultaneously detect non-convex basins and hetero-density basins. (I) The hierarchical structure of these data. On this hierarchical structure, the sizes of basins represent the number of samples and the colors represent different basins. (J) D score of each point. (K) Blue, green and black points indicate three basins. And red points show the attractors of each basin. (L) Simulated single-cell data in three stages (generated by splatter^68^). These data included 1000 cells belonging to three stages in the 5000 genes space and these cells were projected into principle component space. (M) MarkovHC detected 3 basins on level 17 (brown, green, and blue) and the red points in each basin indicated the attractors. (N) ARI (adjust rand index) was used to evaluate the clustering accuracy of MarkovHC, Seurat, SIMLR, sc3, k-means, hierarchical clustering with average linkage, HDBSCAN, spectral clustering, and model-based, which showed that MarkovHC performed best on these data. (O) The orange points indicated the points on the transition path (orange line) from the brown basin to the blue basin. The purple points indicated the critical points. (P) The heatmap of differentially expressed genes along the transition path in (O). (Q) Three representative genes with different “gene-flow” trends in (P). The purple points indicated the critical points.

The Markov chain modeling state transition progress includes the possibility of a state returning to a previous state, by which transition paths can be modeled precisely and reasonably. The asymmetric δ connectivity gives MarkovHC the superiority to find the transition path among the basins and the critical points on the transition paths. On a specific level, with a given start basin, the transition path is the shortest path from its attractors to another basin. Within the transition path of two cell types, only a minority of cells (ridge cells) are critical in the progress^59^. If cells of one type transit to the critical points, these cells have a big chance to transit to another type. Meanwhile, detecting these critical points is of great importance in applications, for example, diagnosing the disease at an early stage^8^. As illustrated in Fig. 2A, the transition path (red box with arrows) and the critical points (black edge points) may be found.

For demonstration, we applied MarkovHC to a set of simulated points, consisting of two spheres with a bridge connecting them in a 3-dimensional Euclidean space (Fig. 2B). MarkovHC successfully identified the two basins (green, blue) and attractors (red) located at the core with high-density respectively. Meanwhile, the transition path (orange) between these two basins were tracked. Note that the critical points (purple) around the gap without topological continuity on the transition path were detected.

To accelerate computation, we used some simple clustering methods such as k-means to cluster original points into hundreds of small clusters at the first step when MarkovHC was used to analyze data sets with more than one thousand single-cells. We regarded these small clusters as new aggregated points for further processing. As long as the primary topology of the data is kept, it should be legitimate and helpful to do pre-clustering^60^. An example with the similar topological structure in Fig. 2B but with more points (2100) was analyzed with MarkovHC similarly and results were shown in Supplementary Fig. 1A-D. The results demonstrate that the base clustering will not destroy the hierarchical structure built by MarkovHC as long as the sizes of small clusters are sufficiently small to avoid clustering heterogeneous points in the same small cluster.

### MarkovHC is able to cluster non-convex data, hetero-density data, and differentiation stages in a continuum

Single-cell data in high-dimensional gene space is complicated. Therefore, clustering algorithms that assume data are convex may not always be suitable for single-cell data. For instance, k-means and complete linkage hierarchical clustering get biased results in single-cell data clustering^61^. In addition, the heterogeneity and size variety of cell populations also bring obstacles to traditional density clustering. For instance, DBSCAN^62^ is not applicable to identify rare cell types. In many cases, such as tissue development and cell differentiation, the single-cell data are in a continuum, which makes it hard to detect reasonable cluster boundaries. One solution would be using the topology of the data, which only depends on δ-connectivity^5^. Several works have proposed clustering methods based on topology^63,64^ and achieved improved performance. MarkovHC also takes advantage of topological connectivity, which embeds the topology of the original data into basins.

We tested MarkovHC on simulated data which consists of four groups of points in different shapes in a two-dimensional space (Fig. 2F). Within each group, points at the edge of the basins are colored grey (Fig. 2F) to indicate low aggregation, while central points are colored black to indicate high aggregation. As Fig. 2G shows, density scores (D scores) reflected point aggregation degree. A hierarchical structure from level 23 to 26 is sketched in Fig. 2I. Three basins could be obtained on level 26 (Supplementary Fig. 2C). On level 25, the green basin split into two basins (brown and green, Supplementary Fig. 2B) with the critical points (purple) on the bridge connecting them. On level 24, the brown basin on level 25 was further divided into two basins (brown and dark brown, Supplementary Fig. 2A), while the latter split into two smaller basins (dark and light brown, Fig. 2H).

Next, we performed MarkovHC on a simulated single-cell data with three stages in a continuum generated by splatter^65^. The data include 1000 cells in three stages in a 5000 genes space. These 1000 cells were projected onto the principal component space and colored according to their stages (Fig. 2L, Supplementary Fig. 3B). Meanwhile, these are “low-density valleys” on the density landscape, which correspond to energy barriers separating basins on the pseudo-energy landscape (Supplementary Fig. 3A). On level 17 these basins and corresponding attractors were identified (Fig. 2M). In comparison with other algorithms, adjust rand index (ARI) (Fig. 2N) and normalized mutual information (NMI) (Supplementary Fig. 3C) were calculated, which showed that MarkovHC performed better than other methods on this dataset. Specifically, we checked the transition path (orange) and the critical points (purple) from the brown basin to the blue basin (Fig. 2O). The differentially expressed genes along the transition path (Fig. 2P) and the dramatic changes of gene expression (Fig. 2Q) around the critical points demonstrate faithfully the kinetics of gene expression along the transition path.

In the above simulations, we verified three merits of MarkovHC. Firstly, the sparse basin (grey in Fig. 2H and Supplementary Fig. 2) was successfully detected, demonstrating the ability of MarkovHC to detect small basins (rare types) sensitively. Secondly, non-convex basins with arbitrary shapes (blue in Fig. 2H and Supplementary Fig. 2, helixes in Fig. 2K) could be handled by MarkovHC. Last, MarkovHC could find clusters in the continuum (Fig. 2M). These results demonstrate the potentials for MarkovHC to identify rare cell types, handle non-convex in high-dimensional single-cell data, and track development trajectory.

### MarkovHC can consistently achieve accuracy in clustering diverse types of single-cell omics data

Accurate clustering is the foundation for correct identification of cell populations and new cell types. Moving to the real data, we compared MarkovHC with popular single-cell RNA-Seq clustering methods including Seurat, SIMLR, SC3, k-means, hierarchical clustering with average linkage (ALINK), HDBSCAN, spectral clustering and model-based clustering on five benchmark datasets^34,66^ (Fig. 3A-E). The ARIs and NMIs of MarkovHC are comparable to those of these popular clustering methods (Fig. 3F). The comparison among MarkovHC, Seurat with default parameters, and hierarchical clustering with other linkages are in Supplementary Fig. 4. Remarkably, the cell atlas of C. elegans embryogenesis^66^ (Fig. 3E) is a golden benchmark to test development trajectory and lineage clustering algorithm, in which 6090 cells were annotated as 27 sub-cell types. MarkovHC constructed the hierarchical structure of this benchmark and reconstructed sub-cell types on level 4 yielding 0.64 and 0.73 on ARI and NMI respectively (Fig. 3F), which outperformed the other methods.

**Figure 3.**
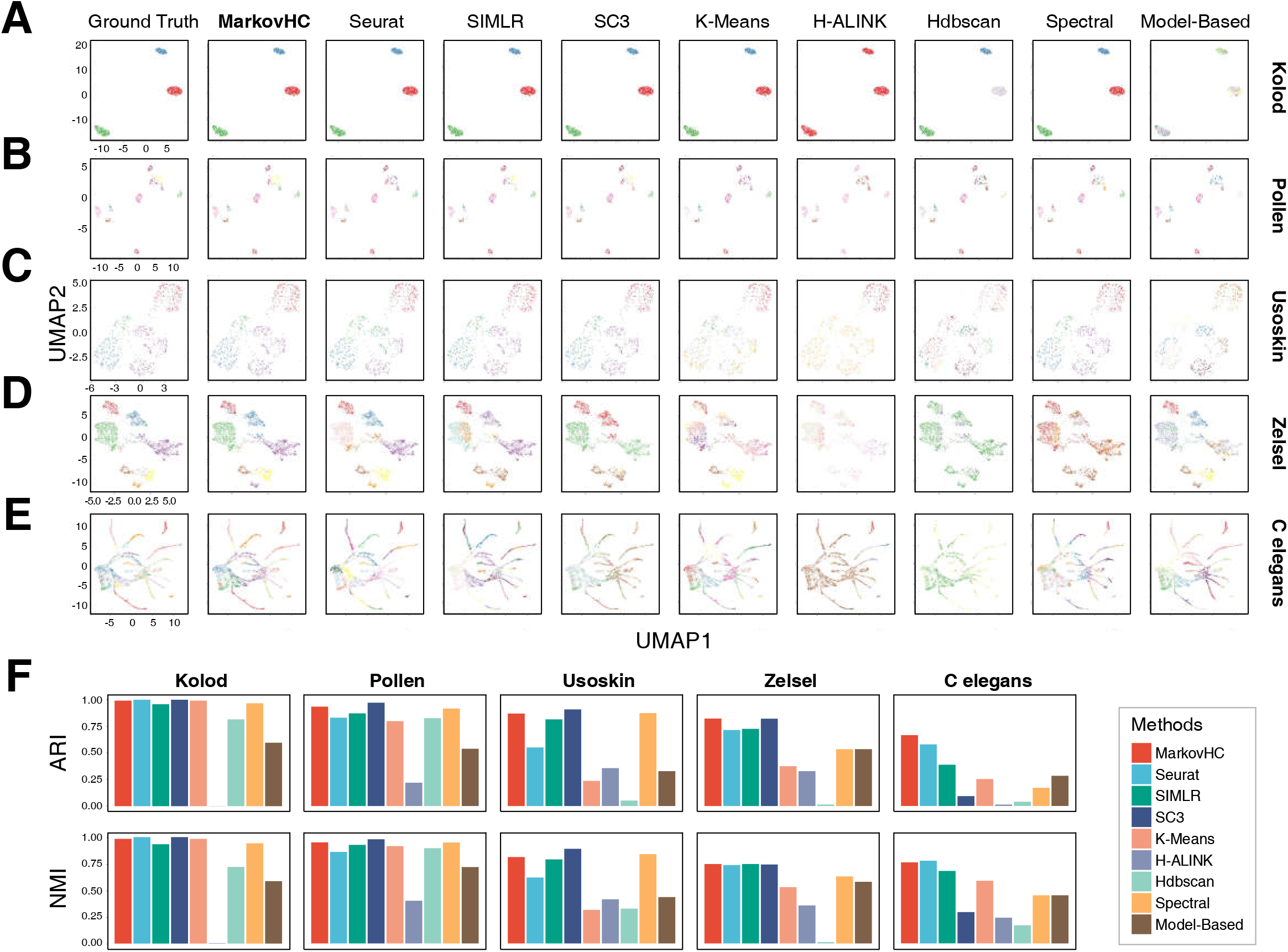
MarkovHC can consistently achieve accuracy in clustering diverse types of single-cell omics data. (A-E) Ground truths of five benchmark single-cell RNA-Seq data sets^34,66^ and clustering results of MarkovHC, Seurat, SIMLR, SC3, k-means, hierarchical clustering with average linkage, HDBSCAN, spectral clustering, and EM. (F) ARI (Rand Adjust Index) and NMI (Normalized Mutual Information) showed MarkovHC can consistently achieve top accuracy in clustering real data.

In addition, the classification and hierarchical structures of human peripheral blood mononuclear cells have been well studied in many researches^67–69^. We applied MarkovHC on the 33k PBMCs dataset to validate the applicability of MarkovHC in analyzing datasets with a large number of cells. On level 113, basins correspond to B cells, T cells, NK cells, megakaryocyte, and monocyte (Supplementary Fig. 5) could be identified. To identify subtypes of basins, we checked lower levels on the hierarchical structure. On level 111, the T cell basin was divided into CD4+ T cell basin and CD8+ T cell basin. On level 110, CD4+ T cell basin was further divided into CD4+ naïve, and CD4+ memory T cell basins. Similarly, the subtypes of B cell, CD8+ T cell, and monocyte could also be recovered on lower levels (Supplementary Fig. 5). These results demonstrate the power of the hierarchical structure, with which a specific cell type can be identified using marker genes, and the sub-cell types or cell states can be checked in detail on lower levels.

Moreover, MarkovHC is based on topology among samples without an assumption of data distribution, it can be widely used to build the hierarchical structures of other omics data besides scRNA-Seq data. As long as an appropriate similarity matrix is provided, MarkovHC algorithm can be applied. In single-cell research, another widely used technology in profiling cell populations is mass cytometry. Here, we verified the applicability of MarkovHC on mass cytometry data from Anchang’s paper^60^, which includes about 150,000 cells and 38 parameters. In this dataset, cells were obtained from a healthy donor’s bone marrow and 24 subpopulations were labeled according to the protein expression of 38 surface markers. We randomly sampled 10009 cells from the dataset in proportion and projected these cells into 2-dimensional space by UMAP (Fig. S6). The results showed that MarkovHC got a comparable performance to Seurat (Fig. S7).

Nevertheless, MarkovHC can help to perform integration analyses, for instance, the integration of scRNA-Seq and scATAC data. In Supplementary Fig. 8, there are 464 scRNA-Seq and 96 scATAC-seq samples generated for the retinoic acid-induced mESCs differentiation at day 4 in Zeng *et. al.’s* work^70^. MarkovHC identified three basins in scATAC-Seq (Supplementary Fig. 8A) data and scRNA-Seq (Supplementary Fig. 8B) data respectively. We used tSNE to visualize these data (Supplementary Fig. 8A, B), and identified differentially chromatin accessibility in scATAC-Seq (Supplementary Fig. 8C) and differentially expressed genes in scRNA-Seq (Supplementary Fig. 8D). The peaks locating highly expressed genes are more accessible than those of repressed genes. For instance, PDE10A, RSPH1, and FBXL17 were highly expressed in basin 1. And the peaks, chr17-8725942-8730132, chr17-31394370-31396459, and chr17-63652707-63655013, which correspond to these three highly expressed genes, were open in basin1. In basin 2, highly expressed genes, RERE and SP8, correspond to open peaks, chr4-149929815-149933175 and chr12-120087188-120088390 respectively. And in basin 3, expression of these genes and accessibility of corresponding peaks were low. Therefore, three basins from these two datasets could be matched according to the peak pattern and the gene pattern respectively, which were also consistent with the three subpopulations inferred by Zeng *et. al.*^70^. These results demonstrate the potential of MarkovHC in the integration of multi-omics data.

In order to verify the effectiveness of MarkovHC in discovering new biological knowledge from real data, we applied it to three other single-cell RNA-Seq datasets, i. e., human embryonic stem cell differentiation to definitive endoderm^6^, human preimplantation embryos^7^, and gastric antral biopsies in the following sections.

### MarkovHC’s hierarchical structure revealed cell lineages, cell types, and cell stages simultaneously

In this section, we show that the hierarchical structure obtained by MarkovHC could be used to explore cell lineages, cell types, and cell stages. In human cell atlas^71^, cell lineage means the developmental relationship of cells; cell type implies a notion of homeostatic persistence; and cell state refers to more inducible or transient properties. However, the boundaries among these concepts at the gene transcriptional level can be fuzzy, partly may be due to the limit of our knowledge and understanding of cellular dynamics. Fortunately, data-driven approaches, especially hierarchical algorithms can be helpful to refine these concepts.

Single-cell RNA-seq data^6^ from experiments profiling snapshots of lineage-specific progenitor cells differentiated from H1 human ES cells in specific induction circumstances was used to illustrate this point. Cells were sorted with specific markers using fluorescence-activated cell sorting (FACS) and captured for single-cell RNA-seq using the Fluidigm C1 system (Scale bar 200μm). In total, 1018 single-cells of six cell types were analyzed in this dataset. All cells were projected into 2-dimensional space using phateR^72^, in which six clusters could be found (Fig. 4A). Low-level results could be swamped by a lot of noises, while high level results may average out true biological signals. Therefore, appropriate levels for downstream analyses need to be chosen based on biological prior knowledge and low dimensional visualization. There were six basins on level 42 (Fig. 4B), around which we did further analyses.

**Figure 4.**
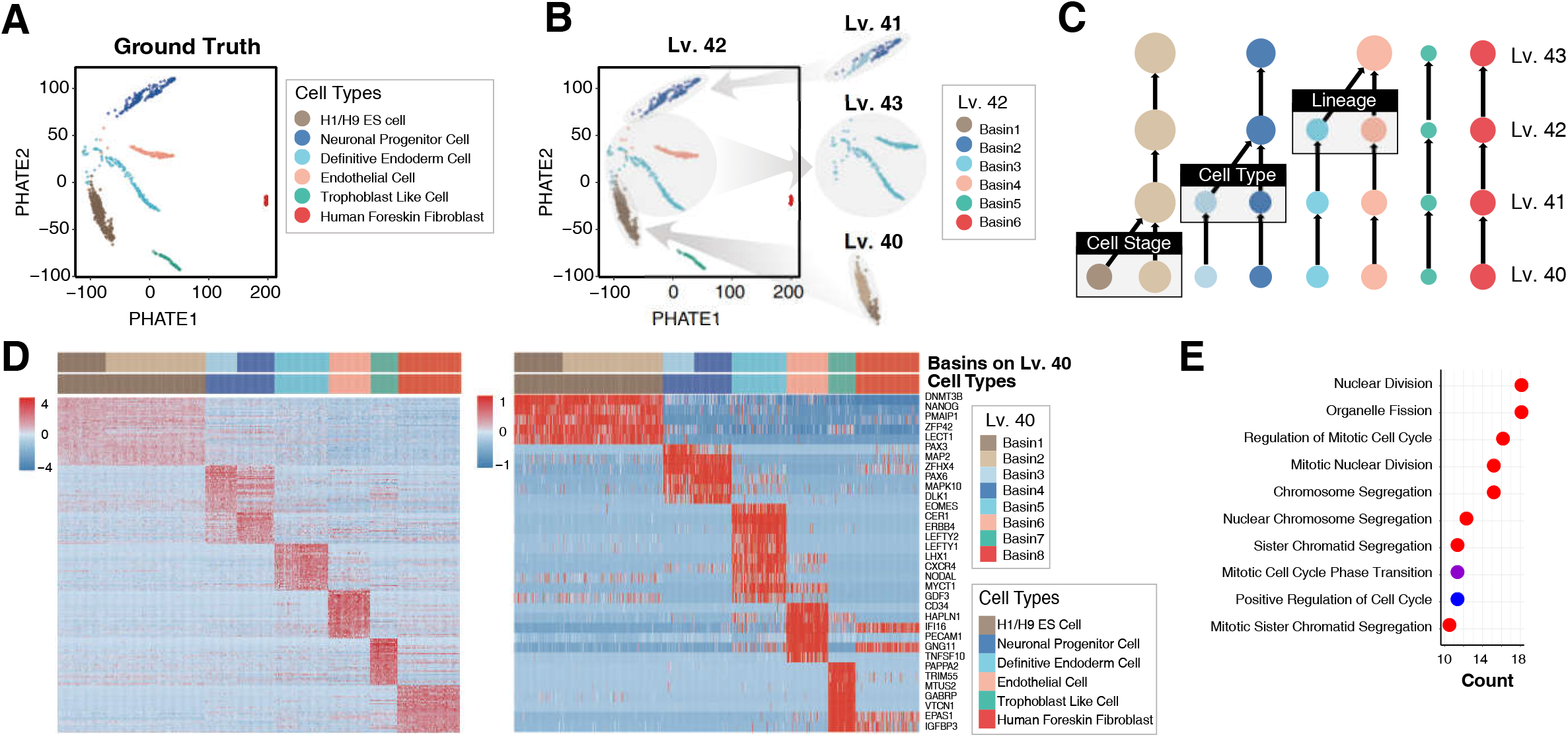
MarkovHC’s hierarchical structure revealed cell lineages, cell types, and cell stages simultaneously. (A) Single-cell RNA-Seq data6 of human ES cell-derived lineage-specific progenitors were projected into 2-dimensional space by phateR. And seven cell types are identified using marker genes. (B) On level 40, two different cell stages were found in H1 and H9 ES cells. On level 41, two cell types in the neuronal progenitor cells were found. On level 42, cells of different lineages were separated. (C) On the hierarchical structure, the difference among basins on level 42 might correspond to different cell lineages, while that among basins on level 41 and 40 might correspond to cell types and cell stages (D) The expression pattern of the 50 most significantly up-regulated genes in each basin (left) and 34 selected marker genes (right). (E) The differentially expressed genes between basin1 and basin2 on level 40 in H1 and H9 ES cells were enriched in cell division related GO terms, which suggested that these two cell stages relate to the cell cycle.

On level 42, the six basins (Fig. 4B) match the six cell types in the ground truth (Fig. 4A). On level 41, we noticed that the neuronal progenitor cells were divided into two basins. Meanwhile, the H1 and H9 ES basin was also divided into two individual basins on level 40. Relations among these basins can be visualized and studied using the hierarchical structure (Fig. 4C). Specifically, on the higher levels, the difference among basins might reflect different cell lineages, while on the lower levels it could correspond to cell types or stages. Meanwhile, the attractors are representative cells of basins, which consist of cells with significantly higher homogeneity than that of other cells in basins (Supplementary Fig. 9).

To closely inspect the difference among these basins, the 50 most significantly up-regulated genes in each basin were obtained by Wilcoxon rank sum test (Fig. 4D left). In checking the biological significance of these basins, we selected 34 lineage-specific markers from Chu’s paper^6^ to show the expression patterns of basins (Fig. 4D right). In these two figures (Fig. 4D), the pattern of basins obtained by MarkovHC through differentially expressed genes (in sheet 2 of Supplementary Table 1) were consistent with that of the selected lineage-specific markers, which demonstrated that the basins were of biological significance. To reveal the difference between H1 and H9 ES cell basins on level 40 and two small basins on level 41 in neuronal progenitor cells, we enriched gene ontology terms using the differentially expressed genes. On level 40, the differentially expressed genes (Supplementary Fig. 10) between the two basins in ES cells were enriched in terms related to cell division and cell cycle (Fig. 4E), which suggests these ES cells were in different cell cycle stages. In the brown basin, cells were in a more active division stage, while cells in the light brown basin were in the suppressive division stage. On level 41, the up-regulated genes (Supplementary Fig. 11) of cells in the blue basin were enriched in neuron differentiation (Supplementary Fig. 12A), while terms were enriched in synapse organization and forebrain development (Fig. S12B) in the light blue basin, which indicated that two subtypes of cells could be found in neuronal progenitor cells. Obviously, the definitive endoderm cell and endothelial cell are two lineages that would be merged on level 43. From the bottom to top in the hierarchical structure (Fig. 4C), cell stages, cell types and cell lineages are presented, which gives a good example of how hierarchical structure can be used to define cell identities and relationships at different resolutions along the developmental lineage trees.

### MarkovHC revealed the transition paths and critical points in human preimplantation embryo development

MarkovHC can be used to track transition paths among cell types and detect the critical points. This dataset includes scRNA-Seq data^7^ of 1529 single-cells collected from 88 human preimplantation embryos at different stages. The overall layout of these 1529 cells in the space of PHATE1-3^72^ can reflect the order of samples (Fig. 5A).

**Figure 5.**
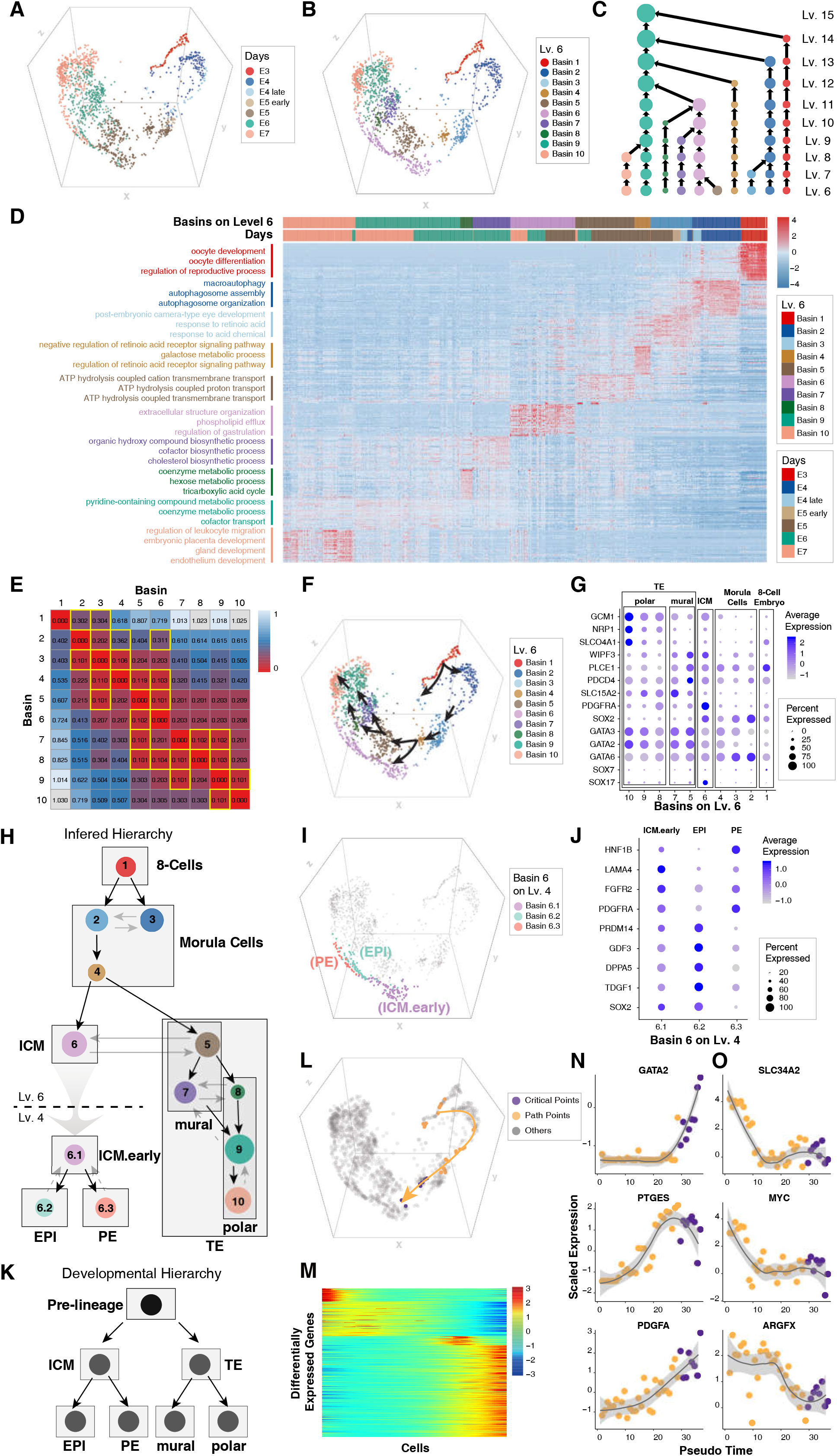
MarkovHC revealed the transition paths and critical points in human preimplantation embryo development. (A) The scRNA-Seq data^7^ of 1529 human preimplantation embryos cells from E3 stage to E7 stage were projected into 2-dimensional space by phateR. In the data, E3–E7 indicates the embryonic day. E4.late and E5.early indicate cells picked 4–6 hours later and earlier than that in E4 stage and E5 stage, respectively. (B) All cells were separated into ten basins on level 6 corresponding to ten asynchronous development stages in human preimplantation embryos cells. (C) The hierarchical structure of these cells. (D) Enriched GO terms of basin-specific gene sets suggest the main biological processes in each development stage. (E) Pseudo-energy matrix (yellow boxes indicate low energy cost) could be used to infer transition among basins step by step. (F) Three trajectories (black arrows) were inferred by the pseudo-energy matrix in (E). (G) Four main cell types which are 8-cell embryo, morula cell, ICM (inner cell mass), and TE (trophectoderm) were identified according to marker genes expression. (H) The development hierarchy was built according to the inferred trajectories in (F). The black arrows indicate development directions, the grey arrows indicate transition relations, and the grey dashed arrows indicate reversed links. (I) ICM (inner cell mass) was split into three basins corresponding to PE (primitive endoderm), EPI (epiblast), and ICM.early (the early stage of inner cell mass) on level 4. (J) Marker genes were used to identify the three basins in (I). (K) The ground truth of the development hierarchy. (L) The transition path (yellow line with the arrow) from 8 cell embryo to ICM were tracked. The yellow points indicate cells along the path and purple points indicate the critical points from morula cells to ICM. (M) Differentially expressed genes along the transition path in (L). (N) Representative genes show increasing “gene-flow” trends along the path. And gene expression varies dramatically around critical points (purple points). (O) Representative genes show decreasing “gene-flow” trends along the path. And gene expression varies dramatically around critical points (purple points).

Cells collected at the same time point can be heterogeneous due to stochasticity in cell differentiation. We first identified homogeneous clusters (Fig. 5B) of cells with MarkovHC and then tracked the transition paths and critical points among these clusters. Meanwhile, the bar plot (Supplementary Fig. 13) shows that there is only one basin in E3, and then this basin separated into two basins in E4. Basin3 is dominant in E5 early and more new basins emerge after that, which suggests that new cell populations burst at that time. As the work from Petropoulos *et. al*^7^ demonstrated, cells in these data were identified as eight cell groups, including pre-lineages, trophectoderm (TE), inner cell mass (ICM), epiblast (EPI), primitive endoderm (PE), E5mid (these cells collected at seven time points), mural and polar (Fig. 5K). For there were eight basins on level 8, we selected the levels (from level 6 to level 15) around level 8 from the hierarchical structure (Fig. 5C) for further analyses.

To reveal the biological processes in each basin, the 50 most significantly up-regulated genes of each basin were identified using Wilcoxon rank sum test (in sheet 3 of Supplementary Table 1). Overall, genes were enriched on gene ontology terms related to the process of embryonic development, while specific groups of gene ontology terms could be found in each basin (Fig. 5D, in sheet 4 of Supplementary Table 1). To infer transition paths among these basins, we calculated the pseudo-energy matrix (Fig. 5E). Because a basin transit to its closest basin (yellow boxes in Fig. 5E) on the energy landscape with the highest probability, we inferred the trajectories (Fig. 5F) and built the development hierarchy (Fig. 5H) which is basically consistent with the knowledge of human embryo development hierarchy (Fig. 5K). To identify each basin, we used the canonical marker genes from Petropoulos *et. al*^7^ and LifeMap Discovery (https://discovery.lifemapsc.com/in-vivo-development/) in the embryonic development (Fig. 5G) and identified four main cell types which were 8-cell embryo (GATA6-, SOX7-, SOX17-), morula cell (GATA6+, SOX7-), ICM (inner cell mass, SOX2+ and PDGFRA+) and TE (trophectoderm, GATA2+ and GATA3+) according to the marker genes expression. To identify sub-groups in TE, we used the differentially expressed genes of mural (WIPF3+, PLCE1+, PDCD4+, SLC15A2+) and polar cells (GCM1+, NRP1+, SLCO4A1+)^7^. To further explore sub-groups in ICM, we checked the sub-basins of ICM on level 4 and found that there were three main basins (Fig. 5I). According to their marker genes expression (Fig. 5J), we identified these three basins as PE (PDGFRA, FGFR2, LAMA4, and HNF1B), EPI (SOX2, TDGF1, DPPA5, GDF3, and PRDM14) and ICM.early. To further check gene-expression “flow” trends along the transition path from 8-cell embryo to ICM, we tracked the cells in the path (Fig. 5L, yellow points indicate cells on the transition path and purple points indicate the critical points from morula cells to ICM). 1631 differentially expressed genes along this path were obtained (Fig. 5M, in sheet 5 of Supplementary Table 1). GATA2, PTGES, PDGFA (Fig. 5N) showed increasing trends, while SLC34A2, MYC, ARGFX (Fig. 5O) displayed decreasing trends. Dramatic changes in gene expression were observed around the critical points (purple points), indicating these points as pivotal stages in the developmental process. These results demonstrate the significance of transition paths and critical points in understanding developmental dynamics and time-series biological data.

### MarkovHC detected the critical points from MSCs to gastric cancer cells

The transition process from normal gastric cells to cancer cells is complex and poorly studied. Many studies have been carried out to better understand the carcinogenesis^73–75^. For example, Zhang *et. al.* conducted a single-cell transcriptomic study on gastric antral biopsies and identified gastric cancer related cell populations. As mentioned above, transition paths and critical points can be helpful to reveal cluster relations. Here, we used MarkovHC to analyze 830 mesenchymal stem cells (MSCs) and 649 MSC-origin early gastric cancer cells (EGCs) (Fig. 6A) from one patient in Zhang *et. al.’s* study^8^.

**Figure 6.**
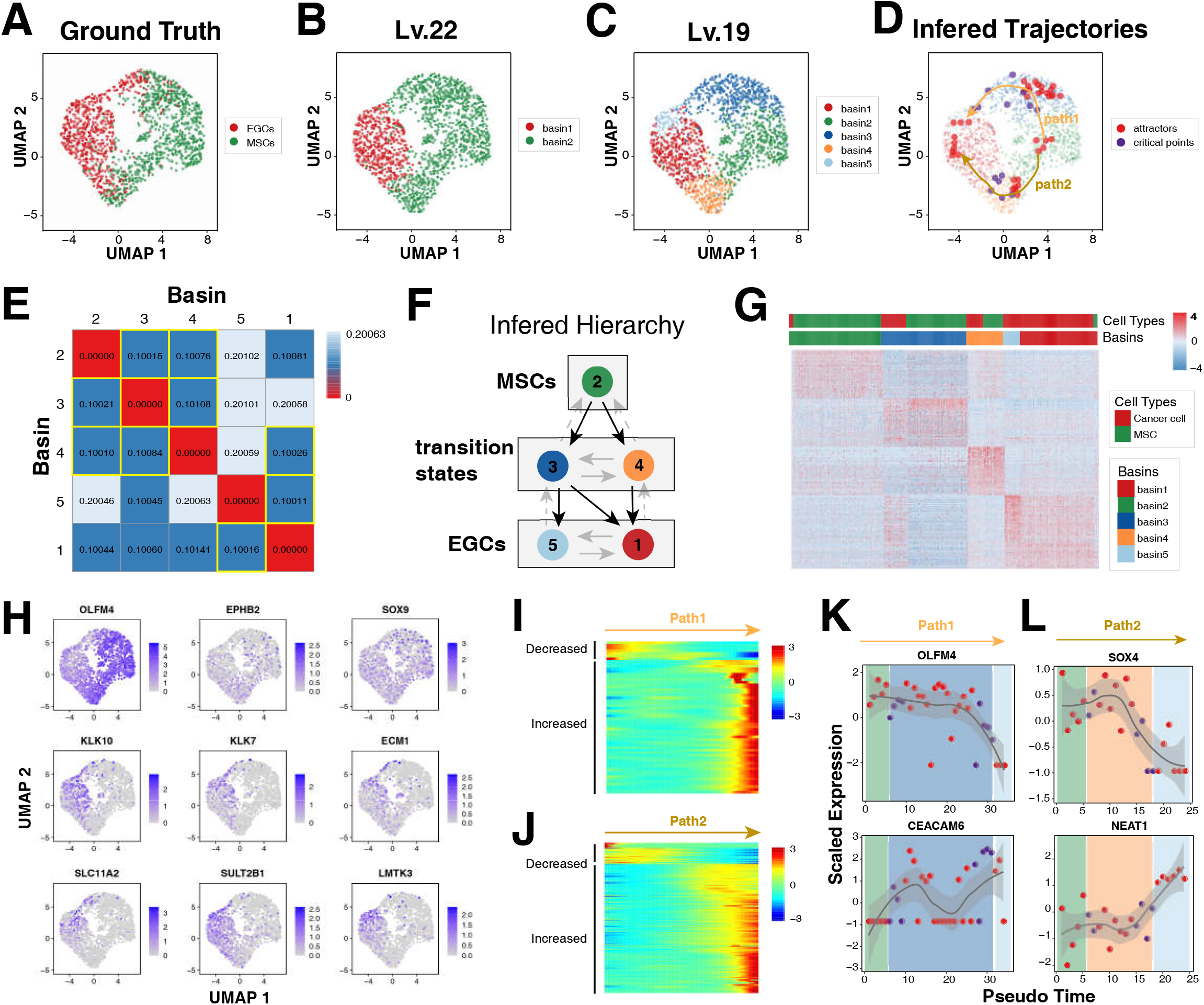
MarkovHC detected the critical points from MSCs to gastric cancer cells. (A) 830 mesenchymal stem cells (MSCs) and 649 MSC-origin early gastric cancer cells (EGCs) were projected into 2-dimensinal space using UMAP. (B) MarkovHC found two basins on level 22. (C) The basin corresponding to MSCs and EGCs split into three basins and two basins respectively on level 19. (D) Two potential separating trajectories from MSCs to EGCs were inferred. Red points indicate attractors of each basin and purple points indicate critical points on boundaries of neighboring basins. (E) Pseudo-energy matrix (yellow boxes indicate low energy cost) could be used to infer transition among basins step by step. (F) The hierarchy was inferred according to the pseudo-energy matrix in (E). The black arrows indicate differentiation directions, the grey arrows indicate transition relations, and the grey dashed arrows indicate reversed links. (G) The heatmap of basin-specific gene sets. (H) MSC markers (OLFM4, EPHB2, and SOX9) and EGC markers (KLK10, SLC11A2, SULT2B1, KLK7, ECM1 and LMTK3) were gradually decreasing and increasing respectively along the trajectories from MSCs to EGCs. (I) Differentially expressed genes along path 1 in (D). (J) Differentially expressed genes along path 2 in (D). (K-L) OLFM4 and CEACAM6 shown different “gene-flow” trends along path 1(K), and SOX4 and NEAT1 shown different “gene-flow” trends along path 2. Red points are attractors, purple points are critical points, and colors of background panels indicate basins on level 19 (C). Meanwhile, expression values of these genes dramatically changed around the critical points.

MarkovHC found two basins on level 22 (Fig. 6B), which was basically consistent with the ground truth (Fig. 6A). However, it was obvious that MSCs and EGCs got mixed up in the boundary region (Fig. 6A). Therefore, we checked the lower levels and found that the MSC basin (green) and EGC basin (red) split into three and two basins on level 19 respectively (Fig. 6C). Not surprisingly, basin 3 and 4 (Fig. 6C) were a mixture of EGCs and MSCs. The differentially expressed genes of these two basins to basin 2 (Fig. 6G) suggested that these EGCs might be in two different transition states. Meanwhile, basin 1 and 5 might be two subpopulations in EGCs (Fig. 6G). Based on these observations, two transition paths from basin 2 to basin 1 were plotted (yellow and brown arrows in Fig. 6D). From the pseudo-energy matrix (Fig. 6E), we could see it would take less energy to transit from MSCs (basin 2) to transition states (basin 3 and 4) than directly to EGCs (basin 1 and 5). Namely, MSCs could transit to transition states with a larger probability. Interestingly, we also noticed that it would cost more energy to transit from basin 3 to EGCs (basin 1 and 5) than from basin 4 to EGCs. This might suggest that basin 3 be more stable than basin 4 on the landscape and reflect the heterogeneity of these two transit states, with the latter having a greater potential to process into cancer. In addition, the two EGC basins (basin 1 and 5) could transit to each other suggesting plasticity and instability of EGCs. We presented the inferred hierarchy of these cell populations in Fig. 6F.

To further investigate the gene expression patterns in these basins, Wilcoxon rank sum test was deployed to find the top 50 differentially expressed genes (DEGs) (Fig. 6G, in sheet 6 of Supplementary Table 1), with which the GO terms were enriched in the relevant functional categories (in sheet 7 of Supplementary Table 1). In the heatmap, we observed relative homogeneity in MSCs (basin 2) and obvious heterogeneity in transition states (basin 3) and EGCs (basin 1 and 5). Specifically, highly expressed genes in basin 2 were enriched in response to topologically incorrect protein and unfolded protein related terms. This suggested that although basin 2 cells were MSC-like cells, they might have been stimulated by environment factors and could be resisting to incorrect protein folding. DEGs in basin 3 were enriched in protein targeting and protein localization related terms, while DEGs in basin 4 were enriched in intranuclear biological processes such as DNA strand elongation and DNA conformation change. The differences indicated that these two transition states responded to environmental stimulus differently. Interestingly, the up-regulated genes in basin 1 were enriched in epidermis development and primary alcohol catabolic process related terms, while those in basin 5 were responses to metal ion and detoxification related terms. These results suggested that the formation of basin 5 might be related to metal ion stimulus, while basin 1 might be associated with alcohol stimulus and epidermis damage. The latter was partially supported by the fact that this patient (p8 in Zhang *et. al.’s* paper^8^) had a chronic alcohol consumption history.

To check the transition trajectories from MSCs to EGCs (path 1 and path 2 in Fig. 6D), we examined the gene expressions of some marker genes along paths (Fig 6.H-J). MSC markers (OLFM4, EPHB2, and SOX9) and EGC markers (KLK10, SLC11A2, SULT2B1, KLK7, ECM1 and LMTK3) were gradually decreasing and increasing respectively (Fig. 6H). DEGs along path 1 and 2 were also identified (Fig. 6I-J), and used for gene function enrichment analysis (in sheet 8 of Supplementary Table 1).

Basically, two clusters of DEGs could be obtained in Fig. 6I and 6J. In path 1, the decreased genes were enriched in response to chemokine, while the increased genes were enriched in cellular response to metal ion suggesting that the potential disease progress might be driven by metal pollution^76^. Specifically, we observed OLFM4^77^, a marker for stem cells in the human intestine, deceased from MSCs to EGCs, while CEACAM6^78^, which plays important roles in invasion and metastasis in Gastric Cancer, was increased (Fig. 6K). In path 2, the decreased genes were enriched in response to mechanical stimulus, DNA damage response and signal transduction by p53 class mediator resulting in cell cycle arrest, while the increased genes were enriched in neutrophil mediated immunity suggesting this path could be the classic p53 related progression of gastric cancer^79,80^. One key gene, SOX4, deceased along path 2 (Fig. 6L), which was consistent with the results that MiR-596 down-regulated SOX4 expression and was a potential novel biomarker for gastric cancer^81^. Another marker, NEAT1^82^, is a long non-coding RNA promoting viability and migration of gastric cancer cells through up-regulation of microRNA-17, which increased along path 2 (Fig. 6L). Furthermore, expression values of these genes dramatically changed around the critical points (purple points in Fig. 6K, L), which suggested these critical points were unstable states locating at the boundaries of the basins. The two paths indicated that there might be two separating trajectories from MSCs to EGCs for this patient, corresponding to two types of potential disease progression routes. These results again demonstrate the great capability and flexibility of MarkovHC in clarifying the relationship of the cell subpopulations and the usefulness of transition paths and critical points in revealing the underlying mechanisms.

## Discussion

In this paper, we developed MarkovHC based on the metastability of exponentially perturbed Markov chain to explore the pseudo energy landscape which corresponds to the intrinsic topological structure of the data. And we also developed an R package, “MarkovHC” (https://github.com/ZhenyiWangTHU/MarkovHC). This tool is easy to use and only requires a feature×cell data matrix as the input. We showed that MarkovHC can cluster cells into populations at different resolutions, which could correspond to cell lineages, cell types, and cell states. Furthermore, we showed that the transition paths and the critical transition points among cell populations tracked by MarkovHC can reveal developmental processes, such as human embryonic development or disease procession.

Besides clustering and identifying differentiation paths, there could be other interesting types of analyses with the help of MarkovHC. First, one could calculate pseudo-time^16,47,48,83^ from the hierarchical structure. Though hierarchical ordering itself already has a causal interpretation, one could refine it using the same framework as shown in this paper. Second, one could start out from a given relevant biological level, e.g. based on prior knowledge or other clustering methods, and use MarkovHC to explore the next level up or down in the hierarchy. Thirdly, the hierarchical structure obtained by MarkovHC could be used to integrating other or multi-omics data, e.g. by using UnionCom^84^ recently developed by Cao *et. al.,* for the unsupervised topological alignment of single-cell multi-omics integration.

Finally, our method, which is based on a general mathematics theory, is flexible and robust. In the pre-processing, we used sNN to derive the initial transition probability matrix in order to overcome sparsity/dropout problems as well as computational burden. One could certainly use any other advanced manifold-learning methods^34,85^ to build the ground similarity matrix. Although such preliminary processing step has little effect^60^ on the topological structure at biologically relevant levels, it is possible to further optimize to best balance the tradeoff between efficiency and accuracy for a specific data set.

## Methods

### Pre-processing of single-cell data

We employed functions in a popular single-cell toolkit, Seurat (v.3.2.1)^30^, to pre-process scRNA-Seq count matrixes. These matrixes were normalized by NormalizeData function with factor 10000, and 3000 highly variable features were selected using FindVariableFeatures function. Then, the normalized matrixes were scaled by ScaleData function. Next, RunPCA function was used to perform linear dimensional reduction and dimensions were selected by an heuristic method generating an “Elbow plot”: principle components were ranked based on the percentage of variance explained by each one, and we always can observe an “elbow”, suggesting that the top PCs before the “elbow” could capture the majority of true signal. And these dimension reduced matrixes were the original inputs to the MarkovHC algorithm.

In particular, the pre-processing steps in DC3^70^ were used to normalize the data of 464 scRNA-Seq and 96 scATAC-seq samples (Supplementary Fig. 8) from their work. And the cytometry data (Supplementary Fig. 6) were scaled by log(x + sqrt(x^2 + 1)) and performed linear dimensional reduction by RunPCA function.

### Overview of the MarkovHC algorithm

The input data of MarkovHC algorithm is the matrix of genes by cells or proteins by cells, representing the detected mRNA or proteins expression in each cell. Without loss of generality, let s_1_, s_2_,…, s_n_ be a set of d-dimensional data points. Traditionally, the clustering goal is to divide them into different basins that accords with intuition and are interpretable:

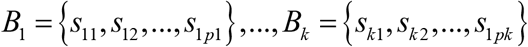 where *{B_i_,i* = 1,…,k} usually have no overlap. As we know, sometimes k, the number of clusters, is cannot be decide beforehand and may be varying depending on different goals. Therefore, the Markov algorithm is allowed to explore the intrinsic hierarchical structure in the data points. Supplementary Fig. 14 shows the target hierarchical structure of MarkovHC, where every 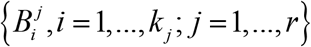 is the subset of {*s*_1_,…,*s_n_*} and 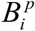 is the subset of some basins on level p+1. Note that we allow 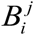 on the same level have some overlaps defined exactly by the shared points. They are of great significance in the hierarchical structure. If there are points shared by 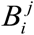 and 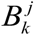, once states in 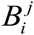 reach to these shared points after a long trip, they can easily develop into points in 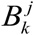 in a short time. Of course, sharing some points is not a sufficient condition for combining 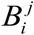 and 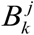 together as. If it is very hard for core points of 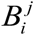 and 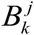 to develop into points, 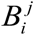 and 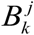 will not combine on the next level. Similar to the structure produced by traditional hierarchical clustering algorithms, when we select a level, there is a clustering result. Unlike those traditional hierarchical structures, structure produced by MarkovHC is more concise and meaningful. Every level’s clustering result is interpretable. Different levels correspond to different δ mentioned in Fig. 2. If we want to obtain a more detailed clustering result, we can select a lower level.

We can define a measure of the relation between different sample points. Some measures are undirected, generally called similarity or dissimilarity. For example, k-means and standard hierarchical clustering usually use Euclidean distance as the dissimilarity between points. Some measures are directed. For example, DBSCAN^62^ gives a directed definition of directly density-reachability. Obviously, a directed relation is more reasonable because in many cases, the status of two sample points is not equal. Let’s take a biological example. Assume A is a point representing a hematopoietic stem cell and B is a point representing a red blood cell. A is able to develop into B while the inverse process is impossible. Therefore, we consider using Markov chains to describe the whole dataset. Continue with the example above, we can set the transition probability p(A, B) > 0 while p(B, A) = 0, which means that A can reach B but B cannot return to A. If we have a Markov chain to describe the relations of different points, it is very natural to see points belonging to a same recurrent class as core points of a cluster. Points that are not recurrent states but can reach to more than one recurrent class can be seen as shared points. However, a single Markov chain is not enough to describe the intrinsic structure of data points on the ground that one Markov chain only corresponds to one clustering criterion. As we have mentioned above, for many problems, building a hierarchical structure can completely show the information contained in data. As a result, we use a family of Markov chains to describe the dataset. We can regard the input dataset as a finite state-space *S*= {1,…,*N}·{P^β^*: *β*∈[0,∞)} are a family of Markov chain transition matrices on S satisfying that there exists:

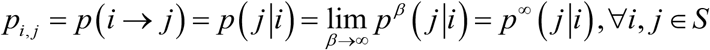

For a fixed β, if state i has a stronger directed relation towards state j, *p^β^*(*j*|*i*) would be bigger. The parameter *β* controls the movement intensity of states. When *β* becomes bigger, the whole state-space will become more inactive that the expected time taken from core states in one cluster to another cluster will exponentially rise. Finally, when *β* approaches infinity, the state movement is basically limited in core points of the same cluster. Taking advantage of the difference of movement time scale between points, we can build a hierarchical structure of the state-space. We assume the following limit exists:

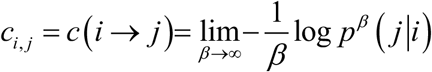

We assume that

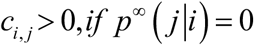

*c_i,j_* can be viewed as pseudo-energy barrier from state i to move to state j.

Under the assumptions given above, Markov chains family, which is composed of Markov chains with transition matrices {*P^β^*: *β*∈[0, ∞)} is an exponential perturbation of the Markov chain with transition matrix *P*^∞^. The core of our theory is the pseudo-energy matrix C, which concentrates all asymptotic information provided by the Markov chain family. Actually, the time scale for point ξ to directly move to point η is 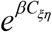. The time scale for points in one cluster to move to another cluster can also be expressed by entries of C. As a result, only with the help of C, we can construct the target hierarchical structure iteratively.

Level 1 clustering result should be the most detailed and basic one. Therefore, we consider the most inactive Markov chain, i.e. the Markov chain with transition matrix *P^β^*. Firstly, we find out all recurrent classes and see them as core points of the corresponding level 1 clusters. Actually, if ξ and η belong to a same recurrent class, there exists a path led from ξ to η such that the sum of pseudo-energy along this path is 0, which means that ξ can move to η effortlessly, and vice versa. We denote these level 1 recurrent classes as 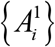, also called as “attractors”. We denote the cluster whose attractor is 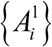 as 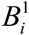, also called as a basin. Then, we need to assign non-recurrent states to appropriate clusters. Given an attractor 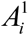 and a non-recurrent state τ, we say τ belongs to basin 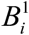 if there exists a directed path from τ to 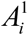 that the sum of pseudo-energy on this path is 0. Though different attractors do no overlap, it is possible that a non-recurrent state τ shared by two or more basins. We give a simple example in Supplementary Fig. 15. Assume data points are 1-dimensional and they are plot at the top of Supplementary Fig. 15. The value of vertical coordinate is the pseudo-energy, which is defined as the negative logarithm of density here. The time scale for points in 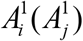 to reach τ is 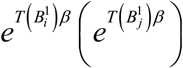 while the time scale for τ to reach 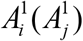 is a constant irrespective of β. (*T* 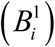 can be viewed as the least pseudo-energy cost for points in 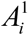 to reach τ.) Therefore, it takes a long time for the points in attractors to move to the shared points while the shared points can jump into the attractors in a very short time. Of course, the time scales for points in 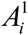 and 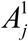 to move to τ are different. Such a difference is exactly the criterion used to define the relation between level 1 clusters (basins) and to construct higher-level Markov chains.

The construction of level 2 clustering result is based on the degree of difficulty for points in level 1 attractor 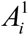 to escape from their corresponding basin 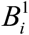, i.e. *T*(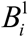). *T*(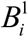) is the least pseudo-energy cost for points in 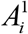 to reach other basins. Considering the characteristic of critical points, if 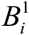 has critical points shared by other basins, then 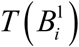 equals to the least pseudo-energy cost for points in 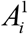 to reach 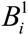’*s* shared points. We construct a new Markov chain to describe the relation between level 1 basins. The new state-space is 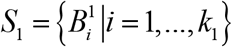, where *k_1_* is the number of level 1 basins. We set the transition probability 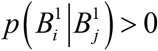 if and only if 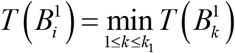 and one of the following two conditions holds:

1. 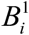 and 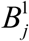 have common shared points;
2. There exists a path from 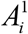 to 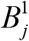 that the sum of pseudo-energy equals to 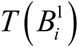. After normalization, we can obtain a transition probability matrix and create a Markov chain on state-space S1. Similar to level 1 clustering result, level 2 clustering results can be easily obtained on the basis of this new Markov chain. We illustrated it in Supplementary Fig. 16. There are four basins on level 1 (Supplementary Fig. 16A). We wrote down some time scales in the figure. For example, the time scale for points in 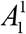 to move to shared points shared by 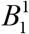 and 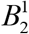 is e^β^. The rest of time scales not shown in the picture are all assumed to be bigger than e^4β^. Based on our criteria, 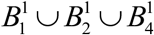 is a new level 2 basin (Supplementary Fig. 16B), which is denoted as 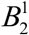. In addition, 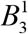 is another level 2 basin, which is denoted as 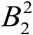. We should note that the attractor of 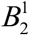 is 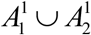, but not 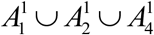, for the reason that moving from 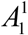 to 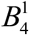 is far more difficult than moving from 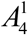 to 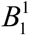.

We can construct the higher-level clustering results in a similar way until the complete target hierarchical structure is finished. Actually, we just transform the information contained in matrix C into a hierarchical structure that can better show the intrinsic relations of different data points. A more detailed derivation, which contains some definitions and theorems regarding to metastability of exponentially perturbed Markov chain is in the Supplementary Notes (rigorous mathematical proofs can be found in the original paper^4^ of Chen *et. al.).*

### The pseudo code

~~~
1 Input the Markov transition matrix P^1^ and the pseudo-energy matrix C^1^
2 P^h^=P^1^, C^h^=C^1^
3 while (TRUE){
#find attractors and partition basins
4 Find all basins belong to recurrent classes in P^h^ and we call them RB^h^ below
5 Use P^h^ to make a directed graph. In the graph, the vertexes are the basins on the current level and the edges are the transition probability between basins. Set i=1.
6   while (TRUE){
7   Select an untraversed recurrent vertex, say 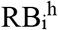, as the start vertex, find all reachable vertexes in the directed graph from 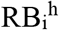 and merge the nodes in the attractors of these reachable basins to be the attractor of the basin on the level h+1, denoted as 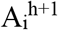.
8   Let 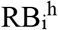 to be the target vertex, find all vertexes in the directed graph that can reach 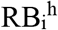 and merge the nodes in these basins to be the basin on the level h+1, denoted as 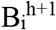. Mark all basins contained in 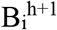 as traversed. i=i+1.
9   if (all B^h^ have been traversed) {break}
    }
10 h=h+1
#Update the pseudo-energy matrix on level h+1
   for(each basin pair on level h+1){
   The energy consumed by basin i transiting to basin j is equal to the minimum energy consumed by the nodes in basin i’s attractor transiting to the nodes in basin j. Update the values in pseudo-energy matrix C^h^}
#Update the Markov transition matrix P^h^ on level h+1
  Find the positions corresponding the values smaller than α% quantile of all values in C^h^. α% is a customized parameter. P^h^ (positions) = 1, P^h^ (other positions) = 0, and convert P^h^ to be a Markov probability matrix by normalizing the sum of each row of it to 1.
11 if(there is only one basin or all basins cannot transfer to other basins){
   return the result
12   }
13 }
~~~

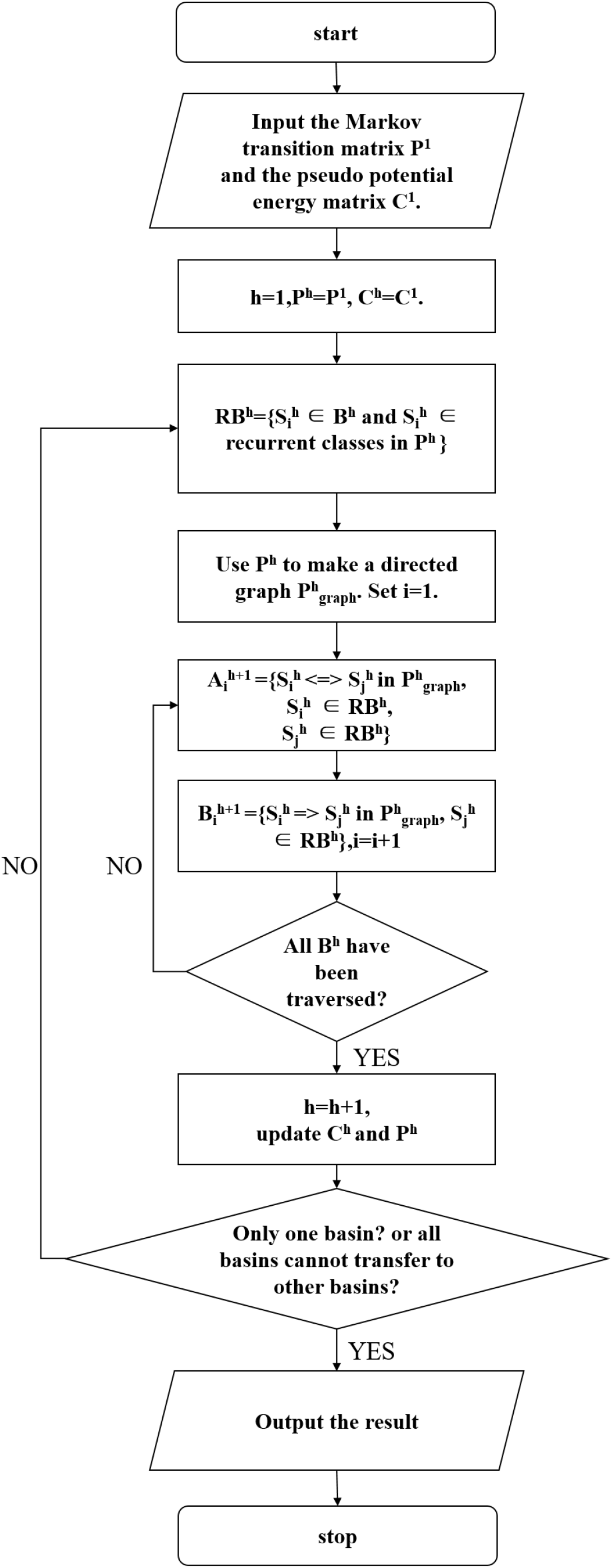
The flow chart.

To build the level h+1 in the hierarchical structure, we need to use the Markov transition matrix P^h^. At first, we must find all basins belong to recurrent states in the Markov transition matrix P^h^. As the mathematical proof in part B of Supplementary Notes, that if a state i is a recurrent state in the Markov transition matrix, then there must be an eigenvalue 1 with a corresponding eigenvector whose i th entry is non-zero. If a state i is a transient state, then for all eigenvalues 1, the i th entry of the corresponding eigenvector must be 0. In this way, we can quickly identify all recurrent states.

The nodes in the attractors of these recurrent state basins can reach each other with positive transition probabilities are merged to be the attractor of the basin on level h+1. For example, suppose there are 10 basins on level h, and 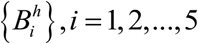 are recurrent states. 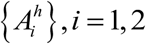 have a positive transition probability path to each other and 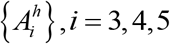 have a positive transition probability path to each other. Then, the nodes in 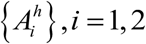 are merged to be 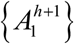 and the nodes in 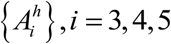 are merged to be 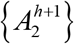. The nodes in other basins, which can reach recurrent state basins with positive transition probability but recurrent state basins cannot reach them, are partitioned to the recurrent state basins. For example, 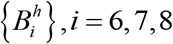 have positive transition probability path to 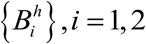 but 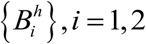 cannot reach them, then all nodes in 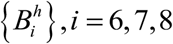 are merged to 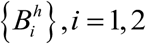 as 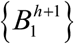.

For implementation convenience, the Markov transition matrix P^h^ is converted to a directed graph. In the graph, the vertexes are the basins on level h and the edges are the transition probability between basins. Let each recurrent state to be the start vertex to find all reachable vertexes from it. The attractors in these basins are merged to be the attractor of one basin on level h+1. For example, 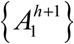.

Next, these basins are used as target vertexes to find all other basins have a positive transition probability path to them. All nodes in these basins are merged to 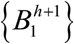. After all the cells have been traversed, we can partition all nodes into basins on level h+1.

To build the level h+2, we need to calculate the Markov transition matrix P^h+1^ representing the transition probability among each basin on level h+1 first. And the matrix P^h+1^ is derived from C^h+1^. On level h+1, the pseudo-energy from 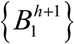 to 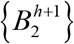 is the minimum pseudo-energy from the nodes in 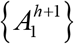 to the nodes in 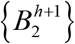. In details, we find the minimum value in the matrix C^h+1^ at first and set 1 to the corresponding positions in the Markov transition matrix P^h+1^. Then, we convert the updated P^h+1^ to a Markov probability matrix by normalizing the sum of each row of it to 1. After we obtain the matrix P^h+1^. The details of building the level h+2 are same as that building level h+1 based on P^h^.

There is an approximation in the Markov transition matrix P^h^ (h>1) calculation. Strictly according to theory, only the positions, which corresponding positions in the C^h-1^ matrix where the value is strictly equal to the minimum, is set to 1 in P^h^. To accelerate basins merging, the entries whose corresponding position in the C^h-1^ matrix having a value smaller than the α% quantile of all values in C^h-1^, are set to be 1 in P^h^. The α% is a customizable parameter.

There’s another optional approximation in this algorithm, which can provide considerable acceleration. On each level, we can merge some small basins to its closest qualified basin when some big basins already form. Naturally, we can finish the construction of a hierarchical structure with fewer iterations. This approximation is built on the belief that if a big basin has already existed on some level, it indicates that we’ve reached a macroscopically meaningful level and hence basins of small sizes can be considered as noise basins. This approximation shows great algorithm efficiency gain and leads to similar results compared with the accurate version when there’s no huge density discrepancy in data. However, when there’s a huge density discrepancy in data, which means that some clusters have obviously larger density over other clusters, we don’t recommend this approximation. This approximation requires three customized parameters to specify the size of noise basins, qualified basins, and big basins respectively. Our empirical recommendation is as follows. We regard basins consisting of less than 10 nodes as noise basins, basins consisting of more than 10 nodes as qualified basins and basins consisting of more than 20 percentage nodes of all as big basins.

Iterating the process above until there is only one basin left or all pseudo-energy consumed by the transition among basins is equal to infinity, which indicates all basins cannot transit to others definitely, the hierarchical structure of the input data is constructed completely.

### Technical details of the MarkovHC algorithm

#### Step1: Calculate sNN similarity matrix and construct a sNN network

The sNN (shared Nearest Neighbors) between two points is the number of K nearest neighbor intersection, therefore sNN is a robust similarity measurement. Single-cell data is high dimensional and has serious dropout events, thus we used sNN to measure similarities among cells and computed a cell-by-cell sNN similarity matrix si,j between node i and node j, which is also equal to the strength of connectivity between the two nodes:

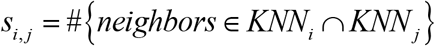

where KNNi is the K nearest neighbors of node i. Then we use the matrix to construct a network. The nodes of the network are cells and the edges of the network are the similarity among cells.

#### Step2: Calculate density score of each node

Accurate density estimation is a challenging task in high dimensional data, especially in single-cell RNA sequencing data. Therefore, we use the degree of each node in sNN network to measure the node aggregation as our density score:

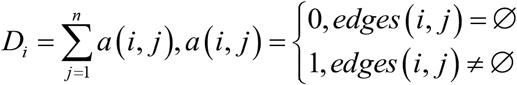

#### Step3: Calculate the Markov transition matrix and the pseudo-energy matrix

The transition probability from node i to node j is defined:

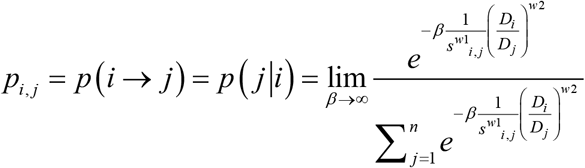

where w1 and w2 are constant parameters determining the tradeoff between the importance of similarity and density score, and both of them are set to 2 by default. For the convenience of calculation, we can derive the concise formula of p(j|i):

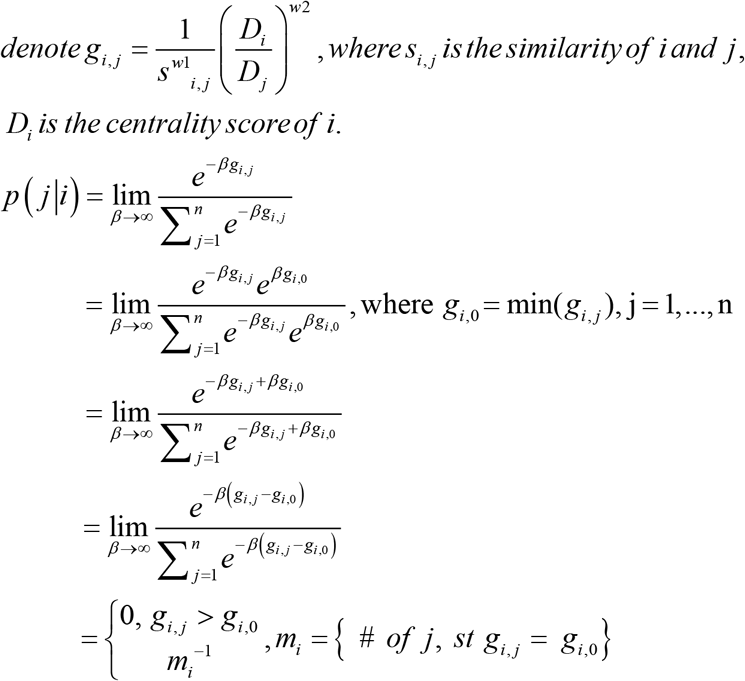

We denote the Markov transition matrix as P1 below.

The consumed pseudo-energy by the direct transition from node i to node j is:

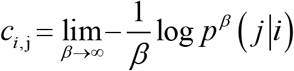

We also can derive the concise formula for calculation convenience:

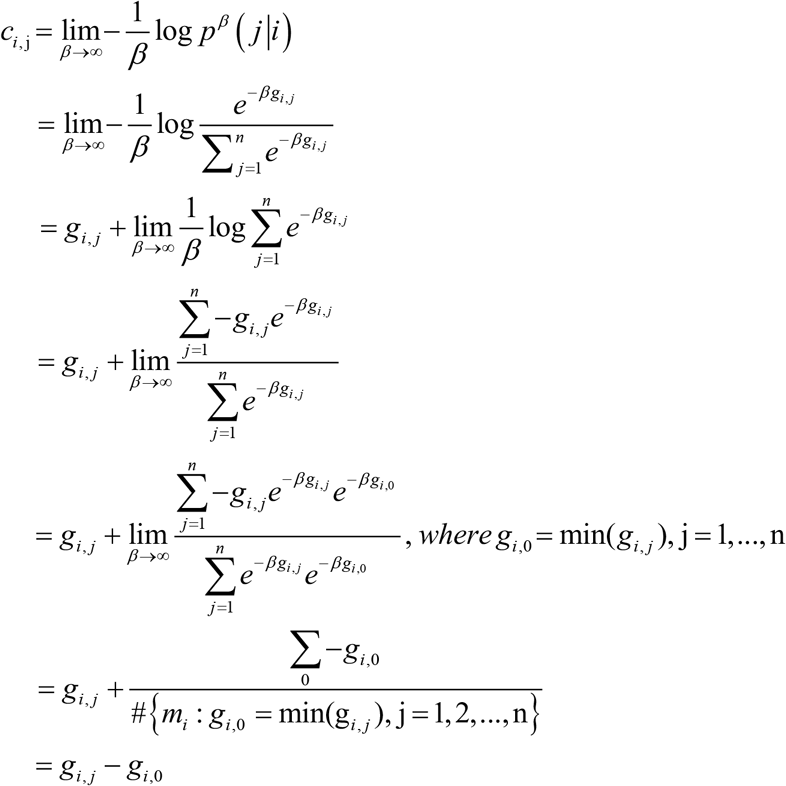

This pseudo-energy matrix represents the pseudo-energy cost when making a direct transition between nodes. We can calculate geodesics (min-energy path) on the surface of this “energy landscape” by using Dijkstra algorithm^57^. If node i cannot transit to node j, the corresponding entry in the matrix is set to be infinite. We denote this reduced pseudo-energy matrix as C1 below.

#### Step4: Pre-clustering

To accelerate the speed of steps below, we can perform pre-clustering at the first stage to cluster all nodes into hundreds of small clusters. These small clusters are homogenous. The number of rough clusters is a customizable parameter in our tools and there are five optional pre-clustering methods, which are k-means, single linkage hierarchical clustering, complete linkage hierarchical clustering, average linkage clustering, and Louvain clustering. The mean density score of the nodes in the small cluster is used as the density score of the small cluster. The number of shared nearest neighbors among nodes is defined as the similarity of them, and the maximum similarity among nodes in two small clusters is used to define the similarity between cluster m and cluster n:

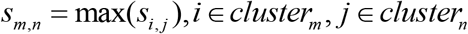

Users could skip this pre-clustering step, but we recommend this for two reasons. First, such preliminary clustering does no harm and won’t largely change the upperlevel hierarchical structure, as long as the number of clusters is large enough after this step. Second, pre-clustering can significantly reduce computational time. By default, we cluster all nodes into 200 small clusters using k-means.

If pre-clustering in this step was performed, we regard small clusters as nodes. For convenience, we still call these ‘small clusters’ as ‘nodes’ below.

#### Step5: Find the attractors and partition the basins of each level to build the hierarchical structure

Firstly, we explain concepts of attractor, basin, the Markov transition matrix P, and the pseudo-energy matrix C. Attractors and basins are sets of nodes. And basins also can be regarded as clusters and attractors can be regarded as cores of clusters. Matrix P represents the transition probability between basins, and the pseudo-energy matrix represents the energy cost by making the transition between the basins.

On level 1, every node is an attractor and also is a basin according to a mathematical proof (Definition1.1 and Definition1.2 of part A in Supplementary Notes). Namely, there are n attractors and n basins for n nodes data. And there is only one node in each basin. Therefore, level 1 has been built directly according to this conclusion. The attractors are denoted as 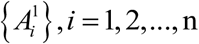, the basins are denoted as 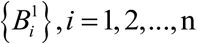, the n×n Markov transition matrix is denoted as P^1^ that represents the transition probability between 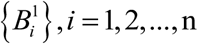, and the n ×n pseudo-energy matrix is denoted as C^1^ representing the pseudo-energy consumed by transition among 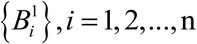.

On a higher level, such as level h, the attractors are denoted as 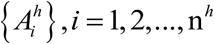, the basins are denoted as 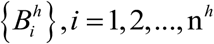, the n^h^×n^h^ Markov transition matrix is denoted as P^h^ that represents the transition probability between 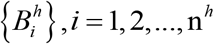, and the n^h^×n^h^ pseudo-energy matrix is denoted as C^h^ representing the energy cost by transition among 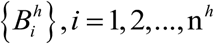.

### Comparison with other methods

It should be noted that there is a resolution parameter in Seurat which is critical to the results, with increased values leading to more clusters. For unknown data, users can only adjust this parameter based on experience and visualization of data. In the process of our evaluation, we fine-tuned this parameter in Seurat to cluster points into the same number of clusters as that in the ground truth and used the results as the best results of Seurat. For comparison, we also used the default parameters of Seurat to get the default results. As for methods that need to set the number of clusters, we set it equal to that in the ground truth. And the results of MarkovHC were taken from the level with the equal number of basins as that in the ground truth.

### Level selection strategy

Selecting levels appropriately from the cluster hierarchy is important for identifying meaningful cell basins.

For users who have enough prior knowledge about data, we recommend a top-down strategy to do it. Firstly, users can identify cell basins on high levels according to marker genes expression. Secondly, users should track the ambiguous basins from high levels to lower levels and identify the sub-basins by marker genes. Lastly, meaningful basins and the relationships among these basins can be extracted from the hierarchical structure by this strategy.

For users who do not have enough prior knowledge about their data, we suggest them to use the plot of “pseudo-energy vs level plot” to assist level selection. As the level increases, the threshold of pseudo-energy cost for basin transition increases accordingly. We divide the “pseudo-energy vs level plot” into two regimes, a noise regime and a signal regime. Low-level results could be swamped by a lot of noises, so the pseudo-energy change slowly in the noise regime. But the pseudo-energy dramatically changes in the signal regime for basins on the high-level may correspond to big cell clusters with biological significance. So levels around the boundary of the noise regime and the signal regime may have biological meaning. We showed the “pseudo-energy vs level plots” of the simulation data in Fig. 2F (Supplementary Fig. 21A) and four real data sets (Supplementary Fig. 21B. data in Fig. 4; C. data in Fig. 5; D. data in Fig. 6; E. data in Supplementary Fig. 20) used in this paper.

### Find the transition paths and critical points

With the help of Markov chain’s directivity, we can determine the transition path, which is the most probable path supported by large deviations theory^86^. And the transition path from basin A to basin B constituted by points on the shortest geodesic path from A’s attractors to B on the pseudo energy landscape. We employed Dijkstra algorithm to find the shortest path in implement. After the transition path was obtained, the two closest points between basin A and B on the path were determined as the critical points from A to B.

### Differentially expressed genes and gene ontology term enrichment analysis

Wilcoxon rank sum test in Seurat (v.3.2.1) was employed to detect differentially expressed genes among cell groups. Differentially expressed genes along the transition path were identified by differentialGeneTest function in monocle (v.2.14.0)^16^. And GO terms of gene sets were enriched by enrichGO function in clusterProfiler (v.3.14.3)^87^.

## Data availability

The links for downloading datasets used in this paper are in the part G of the Supplementary Notes.

## Code availability

R package “MarkovHC” and codes for analysis in this paper are freely available at GitHub (https://github.com/ZhenyiWangTHU/MarkovHC).

## Acknowledgements

This work was supported by Major Program of National Natural Science Foundation of China (81890991), National Key Research and Development Program of China (2018YFA0801402), to M.Q.Z., Major Program of National Natural Science Foundation of China (81890994) to Y.C.. We thank Dr. Peng Zhang, Yuyao Song and Dr. Kui Hua of Tsinghua University for helpful discussion and suggestion.

## Author contributions

Conceptualization: M.P.Q., M.Q.Z., Z.Y.W. and Y.J.Z.;

Methodology: Z.Y.W. and Y.J.Z.;

Software: Z.Y.W.;

Formal Analysis: Z.Y.W.;

Investigation: Z.YW., Z.F.Y., L.Z. and M.L.S.;

Resources: M.Q.Z.;

Data Curation: Z.Y.W. and L.Z.;

Writing - Original Draft: Z.Y.W.;

Writing - Review & Editing: M.Q.Z., Z.F.Y., Z.Y.W., Y.J.Z., L.Z., Y.C., M.L.S. and M.P.Q.;

Visualization: Z.F.Y. and Z.Y.W.;

Supervision: M.Q.Z. and M.P.Q.;

Project Admini strati on: M.Q.Z.;

Funding Acquisition: M.Q.Z.

## Supplementary Notes

### Part. A Definitions of metastability of exponentially perturbed Markov chain.

Consider a finite state space *S* ={1,2,…, *N*}. {*P^β^:β* ∈ [0, ∞)} are a family of Markov chain transition matrices on S satisfying that there exists

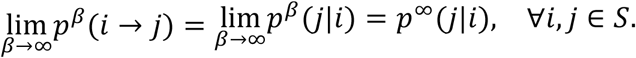

In addition, we assume the following limit also exists,

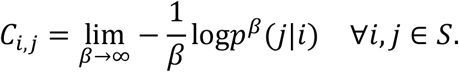

We assume that

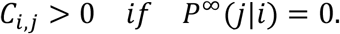

Under the assumptions given above, Markov chains family, which is composed of Markov chains with transition matrices {*P^β^:β* ∈ [0, ∞)}is an exponential perturbation of the Markov chain with transition matrix *P*^∞^.

Let *K* be a subset of *S*, we define *G*(*K*) as the set of maps *g*: *K* → *S* such that *g* maps no non-empty subset of *K* into itself. Let *p* ∈ *K* and *q* ∈ *S\K*. We say that *g* ∈ *G*(*K*) leads from *p* to *q* if there are distinct *x*_0_,…, *x_k_* ∈ *K* satisfying that

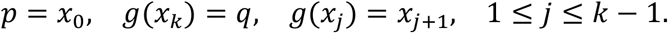

We define

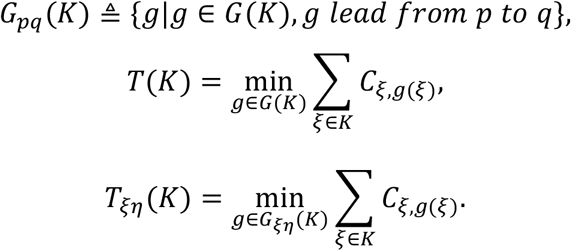

The metastable structure of exponentially perturbed Markov chain is defined iteratively in Chen’s paper^1^. We present some fundamental definitions here.

#### Definition1.1

1. Define *ξ* ⇒ *η* if *C_ξη_* = 0;
2. Define *ξ* → *η* if there is a sequence {*ξ_i_, i* = 0,1,…, *k*} such that

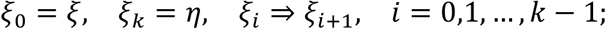
3. Define *ξ* ↔ *η* if *ξ* → *η* and *η* →. *ξ*.

#### Definition1.2

A level 1 attractor 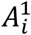 is a subset of *S* such that

1. ∀*ξ*, 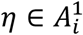, *ξ* ↔ *η*;
2. If 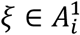, *ξ* → *η*, then 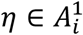.

We can see that, 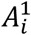 is actually a recurrent class of Markov chain with transition matrix *P*^∞^.

#### Definition1.3

The corresponding attractive basin 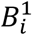 of a level 1 attractor 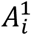 is defined as

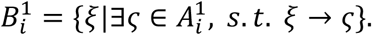

#### Definition1.4

Define 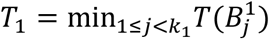, where *k*_1_ is the number of level 1 attractors.

#### Definition1.5

1. Define 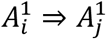 if there exist 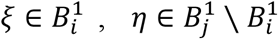 such that 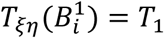;
2. Define 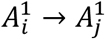 if there is a sequence 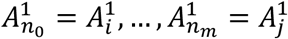 such that

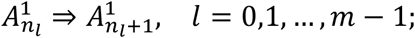
3. Define 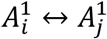 if 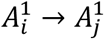 and 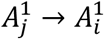.

#### Definition1.6

A level 2 attractor 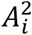 is the union of some level 1 attractors 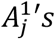, such that

1. 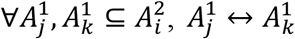;
2. If 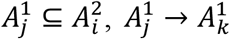, then 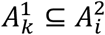.

#### Definition1.7

The corresponding level 2 attractive basin 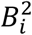 of a level 2 attractor 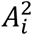 is defined as

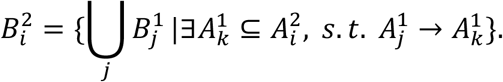

#### Definition1.8

Define

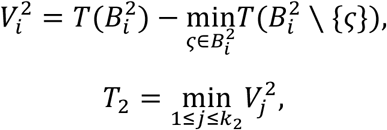

where *k*_2_ is the number of level 2 attractors.

Suppose that we have defined level *k* attractors 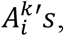, level *k* attractive basins 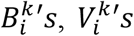 and *T_k_*. Then we define attractors and attractive basins of higher levels as follows.

#### Definition1.9

1. Define 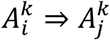 if there exist 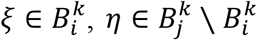 such that

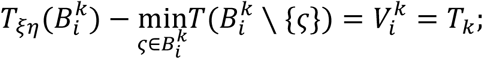
2. Define 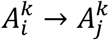 if there is a sequence 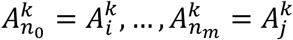 such that

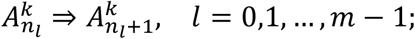
3. Define 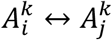 if 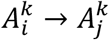 and 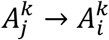.

#### Definition1.10

A level (*k* + 1) attractor 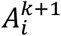 is the union of some level k attractors 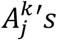, such that

1. 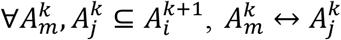;
2. If 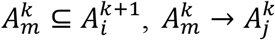, then 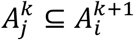.

#### Definition1.11

The level (*k* + 1) attractive basin 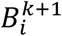 of a level (k+1) attractor 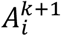 is defined as

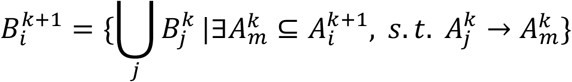

#### Definition1.12

Define

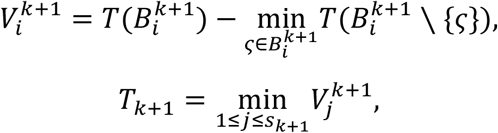

where *s*_*k*+1_ is the number of level (k+1) attractor.

Conventionally, we define 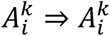 for any *i* and *k*.

Based on definitions presented above, we can construct a hierarchical structure that has been shown in the model part of paper. On a given level k, 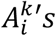 are sets of core points. 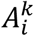 can show the main features of its corresponding cluster, i.e. 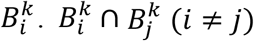 are sets of critical points. We can consider states in 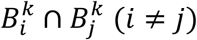 as saddle points. Normally, states in 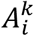 can hardly develop into states in 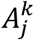. However, once states in 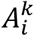 reach states in 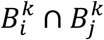, the probability of reaching states in 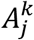 would significantly increase.

As we can see, the essence of the construction of level 1 attractors is to find recurrent classes of Markov chain with transition matrix *P*^∞^. Actually, the essence of higher levels construction is also to find recurrent classes. To achieve this goal, we need to construct a proper higher level Markov chain. On a given level *k*, our target state space is 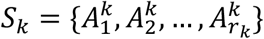, where *r_k_* is the number of level *k* attractors and *P_k_* = (*p_k_*(*i|j*)) is the transition matrix on *S_k_*.

We define an *r_k_* × *r_k_* auxiliary matrix *AU_k_* = (*au_k_*(*i, j*)) as follows,

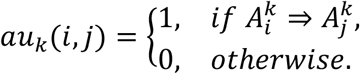

Considering that our goal is to find recurrent classes, we can simply set transition matrix *P_k_* = (*p_k_*(*i|j*)) as follows,

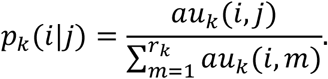

Apparently, the Markov chain constructed above is only an assistant tool. It doesn’t alter the metastable structure of the origin exponentially perturbed Markov chain.

Here, we give further analysis of the metastability of exponentially perturbed Markov chain. Given the basic definition of the metastability of exponentially perturbed Markov chain, we have a basic sketch of our algorithm. The most critical question is how to judge whether 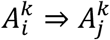 or not. In order to design an applicable algorithm, we present some additional definitions and lemmas here to simplify the procedure of judging whether 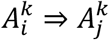 or not.

#### Definition1.13

K is a subset of state space S, for *g* ∈ *G*(*K*), define

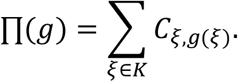

#### Lemma 1.1

For any *i* ∈ {1,2,…, *r*_1_}, where *r*_1_ is the number of level 1 attractors, we have

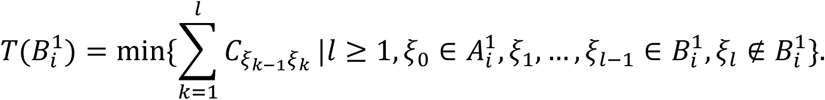

#### Lemma 1.2

Assume *i* ≠ *j*, then

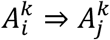

if and only if 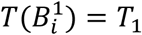 and at least one of the two following conditions holds:

1. 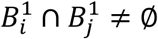;
2. There exists 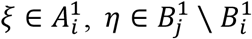 and 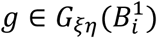 such that 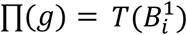.

proof.

#### Necessity

If *i* ≠ *j* and 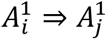, according to definition 1.5, there exists 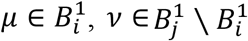 such that 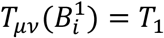. Then we have

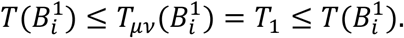

Thus, we can obtain that 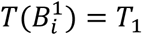. Assume that neither condition 1 nor condition 2 holds. Let h be the map that satisfies 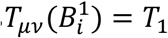, i.e. 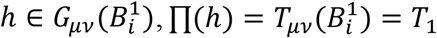. Because condition 2 doesn’t hold, 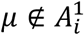. For any 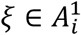, there is a set of positive integers PI such that for any k in PI, 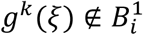. Denote the smallest one in PI as 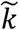 and define 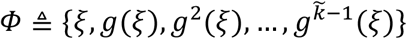. According to lemma 1.1, we have 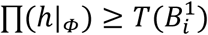. Besides, using the former conclusion, we have 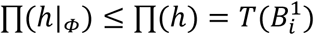. Therefore, we obtain that

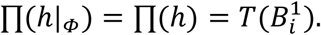

As condition 2 doesn’t hold, 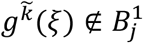. As condition 1 doesn’t hold, 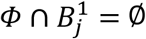. Thus, there is 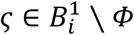 such that *h*(*ζ*) = *v*. But we know that

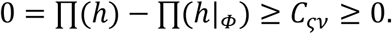

Therefore, we have

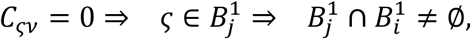

which contradicts the assumption made at the beginning.

#### Sufficiency

Assume that 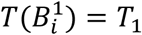. To prove 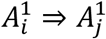, we only need to prove that there exists 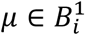 and 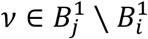 such that 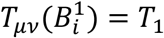. If condition 1 holds, there exists 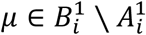 and 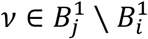 such that *C_μη_* = 0. In addition, according to lemma 1.1, there exists 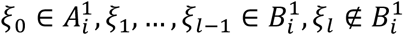 such that 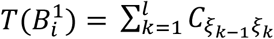.

If 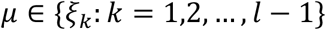, we set *ξ_t_* = *μ*. We define 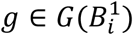 as follows.

Firstly, define

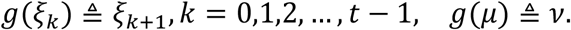

Then, if 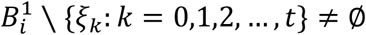, randomly pick a 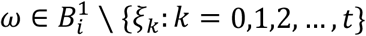. There exists 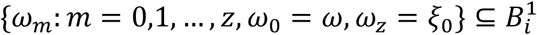 such that 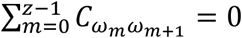. Define 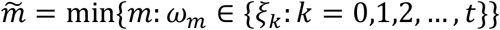. Define

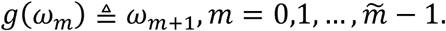

Repeat this process until the definition of g is complete. It is obvious that 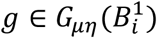 and 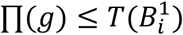. Therefore, we have

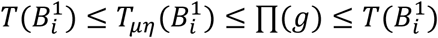

which results in 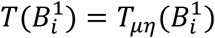.

If 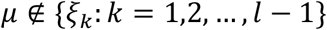, we firstly define

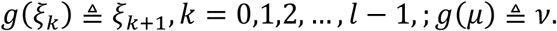

Then we can complete the definition of g in a similar way shown above. Similarly, we can obtain 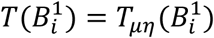. If condition 2 holds, i.e. there exists 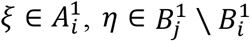 and 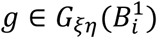 such that 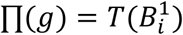, we have

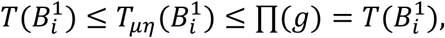

which also results in 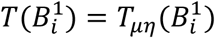.

##### Lemma1.3

Assume 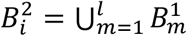, we have

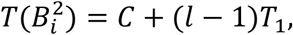

where 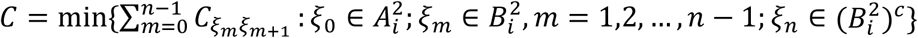.

Proof.

In order to show that 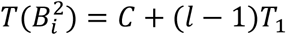, we have to find a map 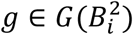 satisfying 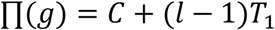 at first. Assume that we have found a path

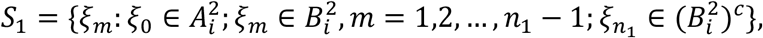

such that 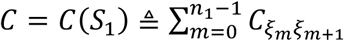 (In order to simplify our proof, we use notations like “path” and “*C*(*S*_1_)”.). Without loss of generality, we assume 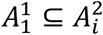 and 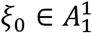. Define *g*(*ξ_m_*) = *ξ*_*m*+1_, *m* = 1, 2,…, *n*_1_ – 1.

According to definition 1.4, 1.5 and lemma 1.1, there exists a path 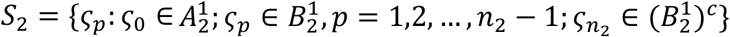 leading from 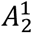 to 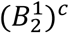.

If 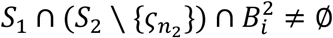, assume the first intersection of path *S*_1_ and path *S*_2_ is *ξ*_*t*_1__ = *ζ*_*t*_2__. Define a new path 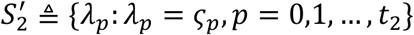.

If 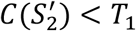, we create a path concatenating the front part of *S*_2_ and the later part of 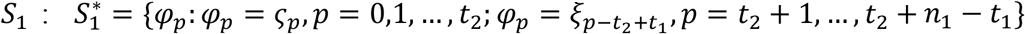. Then we have 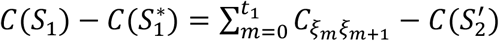. Noticing that 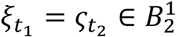, according to the definition of *T*_1_, we have 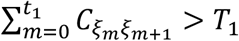, no matter whether 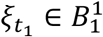 or not. Therefore, we have 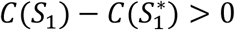, which contradicts with the assumption that *C* = *C*(*S*_1_).

If 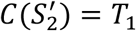, define *g*(*λ_p_*) = *λ*_*p+1*_, *p* = 0,1,…, *t*_2_ – 1.

If 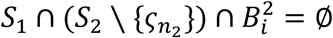, define *g*(*ζ_p_*) = *ζ*_*p*+1_, *p* = 0,1,…, *n*_2_ – 1.

For 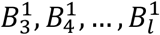, we continue to define g in the same way as above. At last, for any 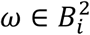 that has not been assigned value under map g, similar to the proof of lemma 1.2, we can find a 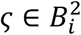 such that *C_ωζ_* = 0. Define *g*(*ω*) = *ζ*. Then, we finish the definition of g such that 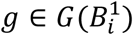 and 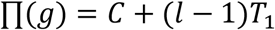.

Nest, we will show that for any 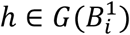, we have 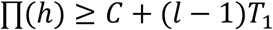. For any 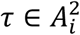, under the map h, there exists a path

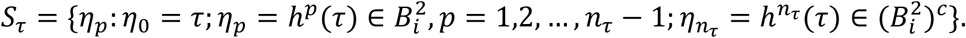

Define 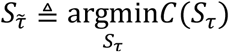. Without loss of generality, we assume 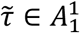 and 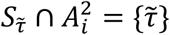.

We will prove 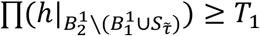 firstly. Because 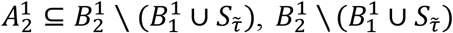 is not empty. We simply denote 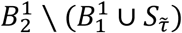 as *M*_2_. For any 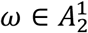, under the influence of map h, there exists a path

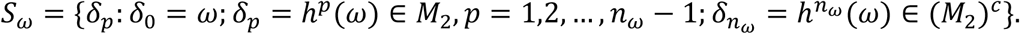

We will use reduction to absurdity to prove *C*(*S_ω_*) ≥ *T*_1_. Suppose *C*(*S_ω_*) < *T*_1_, according to the definition of *T*_1_, we have 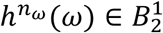, which means that 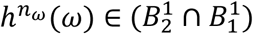 or 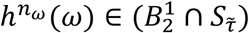:

If 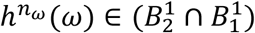, we can easily elongate path *S_ω_* into path

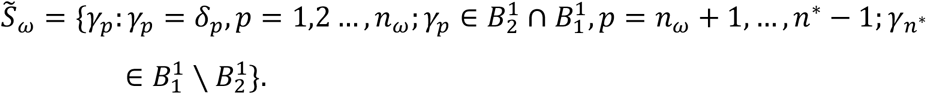

Then we have 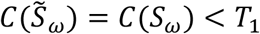, which contradicts with the definition of *T*_1_.

If 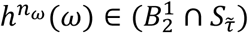, splitting at *h^n_ω_^*(*ω*), we concatenate path *S_ω_* with the later part of path 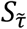 to get a new path 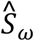. Because *C*(*S_ω_*) < *T*_1_, we have 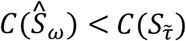, which contradicts with our assumption.

To conclude, *C*(*S_ω_*) ≥ *T*_1_. Further, we have 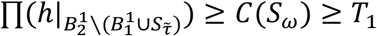. Similarly, define 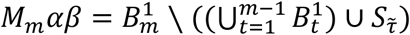, *m* = 2, 3,…, *l*. Using the same method as above, we can prove that 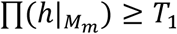, *m* = 2,3,…, *l*. Therefore, we have

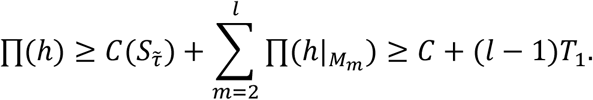

Combining the results of two main parts shown above, we have

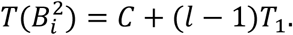

##### Lemma1.4

Assume 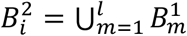, we have

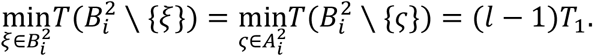

Corollary 1.1

Assume 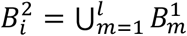, we have

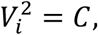

where

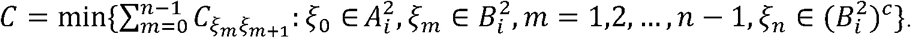

The proof of lemma 1.1 can be found in Chen’s paper. Since the proof of lemma 1.4 is similar to lemma 1.3’s, we skip it. Lemma 1.3 and 1.4 can directly lead to corollary 1.1. As the principles are similar to lemmas and corollary shown above, we only present some lemmas without proofs as follows.

##### Lemma1.5

*k* ≥ 2. Assume *i* ≠ *j*, then 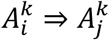 if and only if 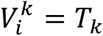 and at least one of the two following conditions holds.

1. 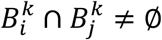
2. There exists 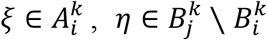 and 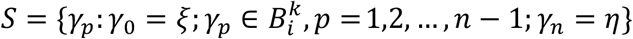 such that 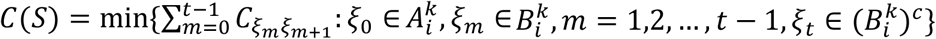.

##### Lemma1.6

*k* ≥ 2. Assume 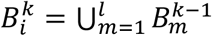, we have

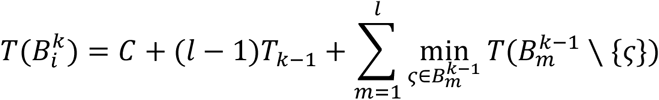

where

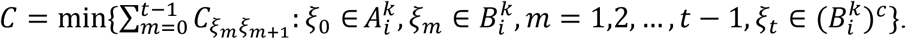

##### Lemma1.7

*k* ≥ 2. Assume 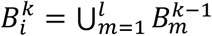, we have

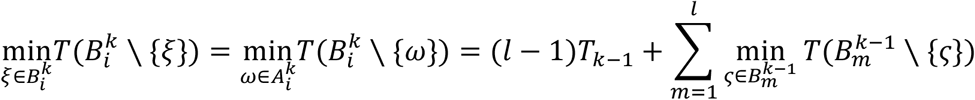

Corollary 1.2

*k* ≥ 2. Assume 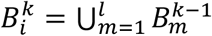, we have

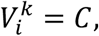

where

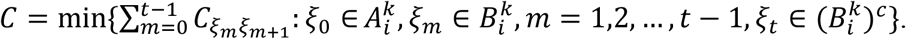

#### Part. B Methods to find recurrent class

In this part, we show how to decide whether a state is recurrent or not and further find out all attractors. Assume we are considering a discrete Markov chain *M*_0_ on finite state space *S*_0_ with transition matrix *P*_0_. Without loss of generality, assume *S*_0_ = {1,2,…, *n*}. Before describing our methods, we present some theorems and propositions to lay theoretical foundations.

##### Theorem2.1

State i is a transient state. *π* is an invariant distribution. Then *π*(*i*) = 0, where *π*(*i*) refers to the ith component of *π*.

##### Theorem2.2

Assume *M*_0_ is irreducible and recurrent. *π* is an invariant distribution. If *M*_0_ is aperiodic, then 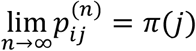, where 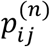 refers to the (*i,j*) entry of (*P*_0_)^*n*^. If *M*_0_ is periodic, 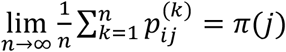.

##### Proposition 2.1

Assume *M*_0_ is irreducible and recurrent. Then there is a unique *π* satisfying *πP*_0_ = *π* and 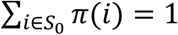. Furthermore, for any *i* ∈ *S*_0_, *π*(*i*) > 0.

Proof.

Denote *Q* to be *P*_0_ – *I_n_*, where *I_n_* denotes the *n × n* identity matrix. For any 1 ≤ *i* ≤ *n*, 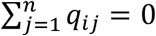, where *q_ij_* refers to the (*i,j*) entry of matrix *Q*. Therefore, *rank*(*Q*) < *n*. Then, there exists a nonzero vector *η* satisfying *ηQ* = 0, i.e. *ηP*_0_ = η. Define 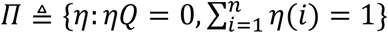. For any 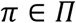, for any positive integer *k*, 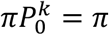. According to theorem 2.2, there exists an invariant distribution *μ* satisfying that 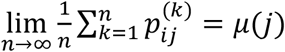. Denote [*μ, μ*,…, *μ*]_*n×n*_ as *Γ*. We have

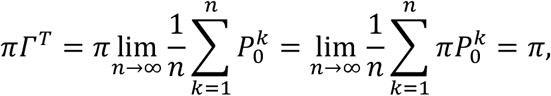

which means *π* = *μ*. Thus, the size of *⊓* is 1. If there exists *i* such that *μ*(*i*) = 0, then select a *j* ≠ *i* satisfying that *μ*(*j*) > 0 and a *k* satisfying that 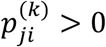. Therefore, 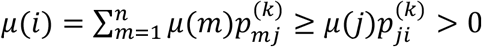, which contradicts the assumption.

##### Proposition 2.2

If *i* is a recurrent state, there exists an invariant distribution *π* of *M*_0_ s.t. *π*(*i*) > 0.

Proof.

Denote *A* to be {*j*: *j* → *i*}. (We say *j* → *k* if there exists {*r*_0_ = *j,r*_1_,…, *r_t_, r*_*t*+1_ = *k*} satisfying that *P*_*r*_*s*_*r*_*s*+1__ > 0, *s* = 0, 1,…, *t*.) Then *A* is closed and all states in *A* are recurrent state of *M*_0_. Denote *M*_1_ to be Markov chain on state space *A* with transition matrix *P*_0_|*_A_*. Then *M*_1_ is finite, irreducible and recurrent. According to proposition 2.1, there exists an invariant distribution *μ* of *M*_1_ subject to that all components of *μ* are positive. Then we define an n-dimensional vector *π* as follows:

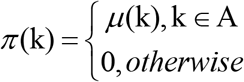

*π* satisfies that *π* is an invariant distribution of *M*_0_ and *π*(*i*) > 0.

##### Proposition 2.3

*i* is a transient state. *π* is a non-zero n-dimensional vector subject to *πP*_0_ = *π*. Then *π*(*i*) = 0.

Proof.

According to theorem2.2, we know that the limit of (*P*_0_)^*k*^ exists and we denote *P* to be 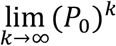. As *i* is a transient state, for any *j* ∈ *S*_0_, 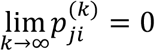. Therefore, all components in the ith column of *P* are 0. Because *πP*_0_ = *π*, we have

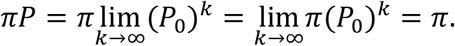

Thus, 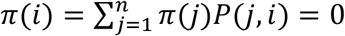.

##### Proposition 2.4

Assume *M*_0_ has k attractors: *A*_1_, *A*_2_,…, *A_k_*. Then the dimension of solution space of equation (*P*_0_ – *I_n_*)^*T*^ *x* = 0 is exactly k.

Proof.

Denote *B* to be 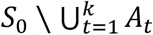. (We assume 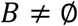 as the 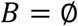 case is just a simpler case.) A non-zero n-dimensional vector x satisfying (*P*_0_ – *I_n_*)*^T^x* = 0 means that *xP*_0_ = *x*. According to proposition 2.3, for any *j ∈ B, x*(*j*) = 0. Denote *P_t_* to be *P*_0_|_*A_t_*_, *t* = 1,2,…, *k*. Denote *M_t_* to be a Markov chain on state space *A_t_* with transition matrix *P_t_, t* = 1,2,…, *k*. Then *M_t_* is a closed, irreducible, recurrent Markov chain. Thus, for *t* = 1,2,…, *k*, according to proposition 2.1, there exists a unique vector *μ_t_* subject to *μ_t_P_k_* = *μ_t_* and 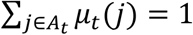. Combining that *xP*_0_ = *x* and *x*(*B*) = 0, we have

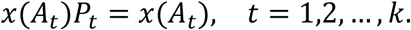

Based on the uniqueness of solution mentioned above, we can easily conclude that the dimension of solution space of equation (*P*_0_ – *I_n_*)*^T^x* = 0 is exactly k.

##### Proposition2.5

Assume the dimension of solution space of equation (*P*_0_ — *I_n_*)*^T^x* = 0 is k. Assume the k orthogonal solution vectors are *π*_1_, *π*_2_,…, *π_k_*. Then for any *i* ∈ *S*_0_, *i* is a recurrent state of *M*_0_ if and only if there exists 1 ≤ *t* ≤ *k*, satisfying that *π_t_*(*i*) ≠ 0; *i* is a transient state of *M*_0_ if and only if for all 1 ≤ *t* ≤ *k, π_t_*(*i*) = 0.

Proof.

If *i* is a transient state, according to proposition 2.3, we know that for all 1 ≤ *t* ≤ *k, π_t_*(*i*) = 0. If *i* is a recurrent state, according to proposition 2.2, there exists an invariant distribution *μ* of *M*_0_, s.t. *μ*(*i*) > 0. Therefore, as *π*_1_, *π*_2_,…, *π_k_* comprise an orthogonal basis, there exists 1 ≤ *t* ≤ *k*, satisfying that *π_t_*(*i*) ≠ 0.

Based on proposition 2.5, we can distinguish whether a state is recurrent or not by computing all orthogonal eigenvectors corresponding to eigenvalues 1 of matrix (*P*_0_)^*T*^. Then we can find out all attractors and basins in the way we introduce in part C. Actually, when we choose to use MHA to analyze a large number of data points, directly computing eigenvectors is not the wisest choice. We should notice that the state with biggest density must be a recurrent state. We can utilize this information to simplify our algorithm.

##### Proposition2.6

Assume *i* is a recurrent state. Denote *T_i_* to be 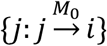. (We say 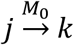 if there exists {*r*_0_ = *j, r*_1_,…, *r_t_, r*_*t*+1_ = *k*} satisfying that 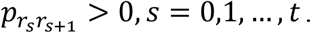) Denote *P*_0_(*T_i_; T_i_*) as 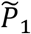. Denote *P*_0_(*S*_0_ \*T_i_; S*_0_ \*T_i_*) as *P*_2_. Normalize 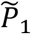 by row into *P*_1_. Denote *M*_1_ to be Markov chain on state space *T_i_* with transition matrix *P*_1_. Denote *M*_2_ to be Markov chain on state space *S*_0_ \ *T_i_* with transition matrix *P*_2_. For any *j* ∈ *T_i_*, j is a recurrent state of *M*_1_ if and only if j is a recurrent state of *M*_0_. For any *j* ∈ *S*_0_\*T_i_*, j is a recurrent state of *M*_2_ if and only if j is a recurrent state of *M*_0_.

Proof.

We define 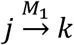 and 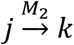 similar to the definition of 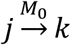 presented above.

Part 1. Assume *j* ∈ *T_i_* and j is a recurrent state of *M*_0_. If *j* = *i*, it is obvious that j is a recurrent state of *M*_1_. If *j* ≠ *i*, because j is a recurrent state of *M*_0_ and 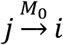, we have 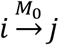. Then we have 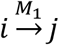, which means that j is a recurrent state of *M*_1_.

Part 2. Assume *j* ∈ *T_i_* and j is a recurrent state of *M*_1_. If *j* = *i*, it is trivial that j is a recurrent state of *M*_0_. If *j* ≠ *i*, because 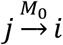, we know 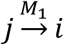. Considering that j is a recurrent state of *M*_1_, we have 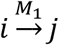, which indicates that 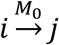. Thus we obtain that j is a recurrent state of *M*_0_.

Part 3. Assume *j* ∈ *S*_0_\ *T_i_* and j is a recurrent state of *M*_0_. For any *k* ∈ *S*_0_\*T_i_* subject to *k* ≠ *j* and 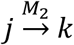, we have 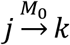. As j is a recurrent state of *M*_0_, 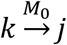 holds. Then we have 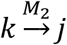. Thus we can conclude that j is a recurrent state of *M*_2_.

Part 4. Assume *j* ∈ *S*_0_ \ *T_i_* and j is a recurrent state of *M*_2_. According to proposition 2.2, there exists a invariant distribution *μ* whose size is equal to |*S*_0_ \ *T_i_*| satisfying that *μ*(*j*) > 0. Then we define an n-dimensional vector *π* as follows:

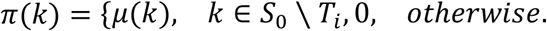

We easily know that *πP*_0_ = *π* and *π*(*j*) > 0. According to theorem 2.1, we can conclude that j is a recurrent state of *M*_0_.

Assume the state with biggest density is *i*. And we use the same notations in proposition 2.6. We divide the origin Markov chain *M*_0_ into two sub-chains *M*_1_ and *M*_2_. It is easy to know that *M*_1_ only contain a single recurrent class, which can be determined by computing the single eigenvector corresponding to eigenvalue 1 of matrix *P*_1_. According to proposition 2.6, this recurrent class is an attractor of *M*_0_. Next, we apply the same method on *M*_2_ because the state with biggest density among states in *S*_0_ \ *T_i_* is also a recurrent state. We do it iteratively until we get all attractors.

### Part. C Comparison with popular clustering methods and trajectory construction methods.

We compared MarkovHC with 19 popular clustering methods (table S1) and 4 representative trajectory construction methods in theory.

These clustering methods including partition-based methods such as k-means^2^, density-based methods such as DBSCAN^3^, OPTICS^4^ and HDBSCAN^5^, graph-based methods such as Louvain clustering (in Seurat^6^), Markov Cluster Algorithm (MCL)^7^ and scanpy^8^, linkage-based methods such as SLINK^9^ (also called single-linkage clustering), ALINK^10^ (also called average-linkage clustering) and CLINK^11^ (also called complete-linkage clustering), spectral method^12^, model-based method^13^, multikernel learning-based methods such as SC3^14^, SIMLR^15^, and hierarchical clustering methods such as classic hierarchical clustering, CIDR^16^, pcaReduce^17^, mpath^18^, SINCERA^19^, TooManyCells^20^, and BackSPIN^21^. These methods have their advantages and disadvantages. For example, k-means methods are classic but often suffer from poor classifications when the data is non-convex and low robustness induced by initial seeds selection. DBSCAN, on the other hand, can handle convex and non-convex clusters but relies on two sensitive parameters to define a dense region, i. e., the radius of the neighborhood and the minimum number of points. HDBSCAN can be seen as a conceptual and algorithmic improvement over OPTICS, which is a density-based clustering method and provides a complete clustering hierarchy composed of all possible density-based clusters. This method extracts the hierarchy from MST (minimum spanning tree)^22^, which introduces biases from its non-unique topology. Louvain clustering methods are widely used, but cannot generate hierarchical structures of single-cell data directly. Even though different clustering results can be used to build a clustering tree by tuning the resolution parameter, the process heavily relies on experiences and might be tricky to skip cell populations of significantly biological meaning due to overly rough partition. SLINK, ALINK, and CLINK can build hierarchical structure, however, the choice of linkages in measuring the similarity among samples is tricky and different orders of processed samples will lead to totally different results. SC3 and SIMLR perform well when applied to some scRNA-Seq datasets, but cannot provide hierarchical structures automatically either. Among these clustering algorithms, Markov Cluster Algorithm (MCL)^7^ invented by Van Dongen et al. is the most similar algorithm to ours. We will compare MarkovHC and MCL detailedly in part D. All problems mentioned above, including non-convex clustering, hetero-density clustering, recognition of small clusters referring to rare cell types, and construction of robust hierarchical structures reflecting data topology, can be solved by MarkovHC.

Meanwhile, a large number of algorithms have been proposed to track cell transition paths. In Schiebinger’s paper^23^, they proposed Waddington-OT and comprehensively reviewed 62 methods, which were categorized into three classes. In Ji’s paper^24^, they compared their TSCAN with 11 other methods. We are not going to make a very detailed comparison in this paper, but only choose four representative methods for discussion, which are Spanning-tree progression analysis of Density-normalized events (SPADE)^25^, monocle series^26–28^, URD^29^, and Waddington-OT^23^. SPADE aims to extract a cellular hierarchy from high-dimensional cytometry data. But it has two inherent shortcomings. Firstly, the density-dependent down-sampling may ruin the original structure in data and obscuring the density information of samples. Secondly, MST (minimum spanning tree) is not unique and very sensitive to parameter selection in the example case^30^(in Fig. S17). Monocle projects original data into a two-dimensional space and orders cells in pseudo time via MST. Hence, the problems of MST also exist in the original monocle. As we reviewed above, MST is widely used in designing trajectory tracking algorithms which may lead to bias. To break these limitations, URD uncovers developmental trajectories from the viewpoint of extending diffusion maps. Briefly, URD operates discrete random walks on a cell graph to approximate the continuous process of diffusion, which can reflect cell differentiation. Although URD can get an approximate transition path by the Monte Carlo method, it cannot get a precise transition path and cannot find critical transition points in the transition path. Waddington-OT is the latest algorithm-based on optimal-transport theory up to date. It can track the ancestors and descendants for every single-cell, however is unable to find critical points on trajectories either. In contrast, MarkovHC calculates a precise transition path consistent with theory and can detect critical transition points simultaneously.

### Part. D Comparison with Markov Cluster Algorithm (MCL).

Markov Cluster Algorithm (MCL)^7^, a graph clustering algorithm based on flow simulation, was invented by Van Dongen et al. in 2000. The MCL is one of the most used clustering algorithms in bioinformatics, but we found there were two points of MCL could be improved. The first one is that the clustering results are sensitive to the parameters used in the expansion and inflation steps. Therefore, it is not clear which clustering result is meaningful. The second one is that MCL cannot get a hierarchical structure. Fig. S18 is from Van Dongen’s thesis^7^. We can see that although the parameter R can control the scale of clustering, the results with different scales cannot be combined to a hierarchical structure. For example, the subgraph in the red circle is split when R is set to 1.4 (in the left top panel).

We ran MCL on a 2-dimensional simulation data (in Fig. S19), which was also used in our manuscript to test MarkovHC (in Fig.2 F). The KNN graph (Fig. S19A) was inputted into MCL. Here, e is the expansion parameter and i is the inflation parameter. And we tested MCL using 18 combinations of parameters. 45 clusters were get using the default parameters (Fig. S19B), in which e and i were equal to 2. Then we set e and i to 1, 180 clusters were get (Fig. S19C). After several adjustments, four clusters were detected (Fig. S19D). Although we tried more combinations of parameters, we couldn’t get better results. And MCL even collapsed under most parameter combinations.

In theory, MCL is based on operating Markov matrix product iteratively, but MarkovHC is based on the global pseudo-energy matrix derived from the Markov matrix. And MCL may miss meaningful partitions but get too rough or too meticulous results for the difficulty in parameter selection. However, MarkovHC will not miss them benefiting from the hierarchical structure. So, we believe MarkovHC has better practical significance.

In summary, although both MCL and MarkovHC cluster points by calculating the geodesic distance and the basic ideas of them are quite similar, MarkovHC has four advantages over MCL. Firstly, the clustering results of MCL are sensitive to the expansion parameter and the inflation parameter, which are difficult to select. But there are no such sensitive parameters in MarkovHC. Secondly, MCL cannot get a hierarchical structure. But MarkovHC can get a hierarchical structure easily. Thirdly, the transition probability of MarkovHC includes density information which is ignored in the transition probability of MCL. Fourthly, the workflow of MarkovHC not only can be used in a secrete-time Markov chain but also can be used in a continuous-time Markov chain.

### Part. E MarkovHC can reveal non-equilibrium biological processes.

In Fig. S20, we used MarkovHC to analysis 280 mouse ESCs in Buettner *et. al.’s* paper^31^. These cells were stained with Hoechst 33342 and Flow cytometry sorted for G1, S, and G2M stages of the cell cycle (Fig. S20A; 96 cells in G1, 96 cells in G2M, and 88 cells in S). We observed three basins match these three stages on level 17 (Fig. S20B). And minimum entries of each row in the pseudo-energy matrix suggest the direction of the cell cycle (yellow boxes in Fig. S20C left).

For the definition of pseudo-energy is

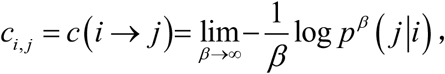

the transition probability matrix can be calculated by

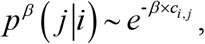

We can get the pseudo-energy matrix of these basins (Fig. S20C left). Without loss of generality, we let *β=5000* in this dataset. Let diagonal entries equal to 0 and perform row normalization to get the transition probability matrix (Fig. S20C right). From the transition probabilities among these three basins, we can get a non-equilibrium ‘flow circle’ which is 1 (G1) →3 (S) →2 (G2M) →1 (G1).

These results demonstrate that MarkovHC can reveal non-equilibrium biological processes.

### Part. F Transition probability estimation.

To quantitatively evaluate how good P_ij_ can fit the ‘real transition probability’, we compared transition probabilities calculated by quadratic programming and MarkovHC.

The authors of Waddington-OT paper^23^ reprogramed secondary (2°) MEFs from E13.5 embryos and collected cells to do single-cell RNA sequencing. We checked these time series scRNA-Seq data and found that sub-clusters in Day2, Day2.5 (the night in Day2), and Day3 can be clustered by Seurat^6^ (Fig. S22) and matched according to the differentially expressed genes (Fig. S23). So we used data in these 3 time point and deployed quadratic programming to calculate the transition probability matrix. The quadratic programming (QP) form of this issue is shown below.

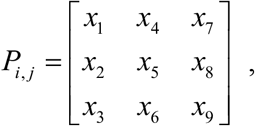

is the transition probability matrix, and we have the cell proportions at 3 time points which can be denoted as 3 column vectors, d1, d2 and d3. So the QP form is

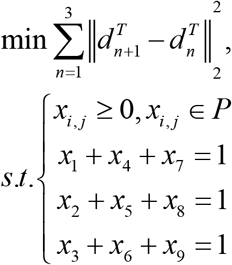

Although the entries in P_ij_ among clusters (Fig. S22) calculated by QP (the left matrix in Fig. S24) and the P_ij_ among basins (Fig. S25) calculated by MarkovHC (the right matrix in Fig. S24) are different, the transition relationships among these basins indicated by P_ij_ are consistent.

### Part. G Data sets.

1. The codes generating simulation data in Fig. 2 are in the ‘MarkovHC’ repository (https://github.com/ZhenyiWangTHU/MarkovHC).
2. The data and labels in Fig. 3A-D were downloaded from ‘SIMLR’ repository (https://github.com/BatzoglouLabSU/SIMLR).
3. The data and lineage labels of C. elegans embryogenesis in Fig. 3E were downloaded from GSE126954.
4. The 33k PBMCs dataset were downloaded from 10X genomics support (https://support.10xgenomics.com/single-cell-gene-expression/datasets/1.1.0/pbmc33k).
5. The mass cytometry of PBMC were downloaded from the supplementary materials of Anchang’s paper^30^ (https://www.nature.com/articles/nprot.2016.066).
6. The scRNA-Seq and scATAC-seq data of the retinoic acid-induced mESCs differentiation at day 4 were downloaded from GSE115968 and GSE107651.
7. The scRNA-Seq data were downloaded from GSE75748. In these data^32^, the labeled cell types include neuronal progenitor cells (NPCs, ectoderm derivatives, n=173), DE cells (endoderm derivatives, n=138), endothelial cells (ECs, mesoderm derivatives, n=105), trophoblast-like cells (TBs, extraembryonic derivatives, n = 69), undifferentiated H1 (n = 212) and H9 (n=162) human ES cells, and human foreskin fibroblasts (HFFs, n= 159).
8. The scRNA-Seq data of 1529 single-cells were downloaded from E-MTAB-3929 (https://www.ebi.ac.uk/arrayexpress/experiments/E-MTAB-3929/).
9. The scRNA-Seq data of 830 mesenchymal stem cells (MSCs) and 649 MSC-origin early gastric cancer cells (EGCs) were downloaded from GSE134520.
10. The scRNA-Seq data of mouse ESCs in Buettner *et. al.’s* paper^31^ was downloaded from E-MTAB-2805 (https://www.ebi.ac.uk/arrayexpress/experiments/E-MTAB-2805/).

## Supplementary Figures

**Supplementary figure 1.**
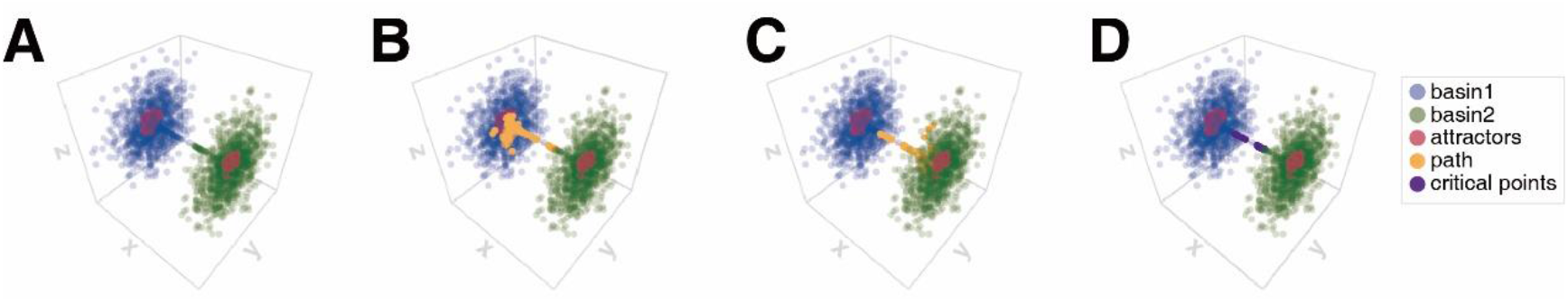
Basins, attractors, transition path, and critical points obtained by incorporating a pre-clustering strategy to save computational time.

**Supplementary figure 2.**
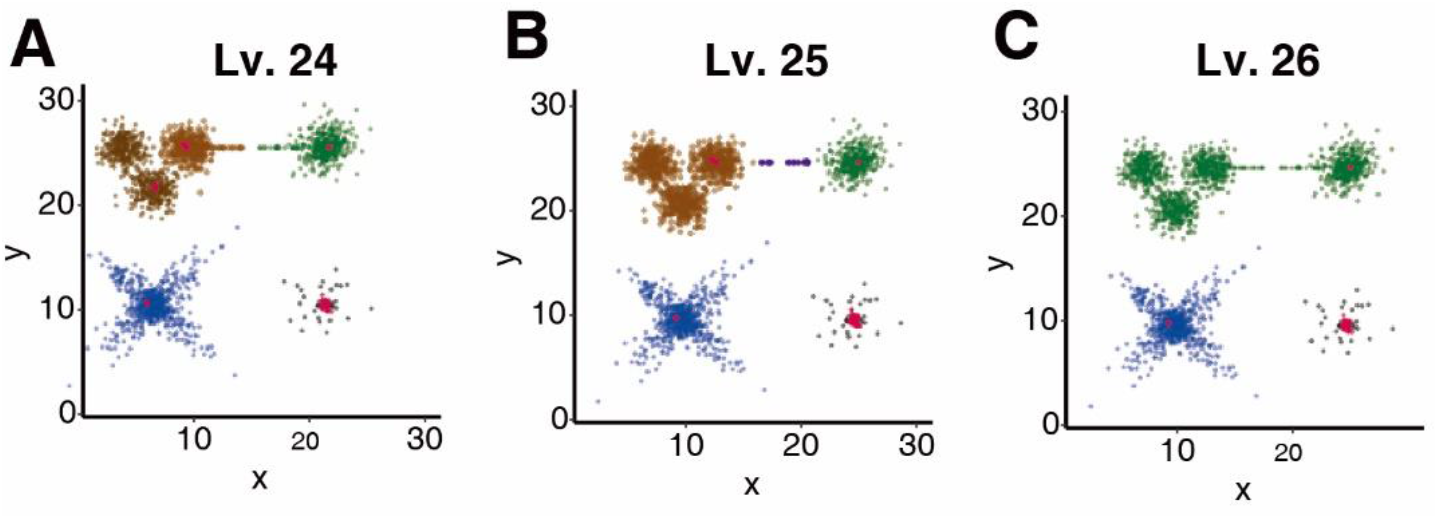
The basins, attractors and critical points on level 24, 25 and 26. A. On level 24, the brown basin on level 25 was further divided into two basins. B. On level 25, the green basin on level 26 split into two basins (brown and green) with critical points (purple) on the bridge connecting them. C. Three basins could be found on level 26.

**Supplementary figure 3.**
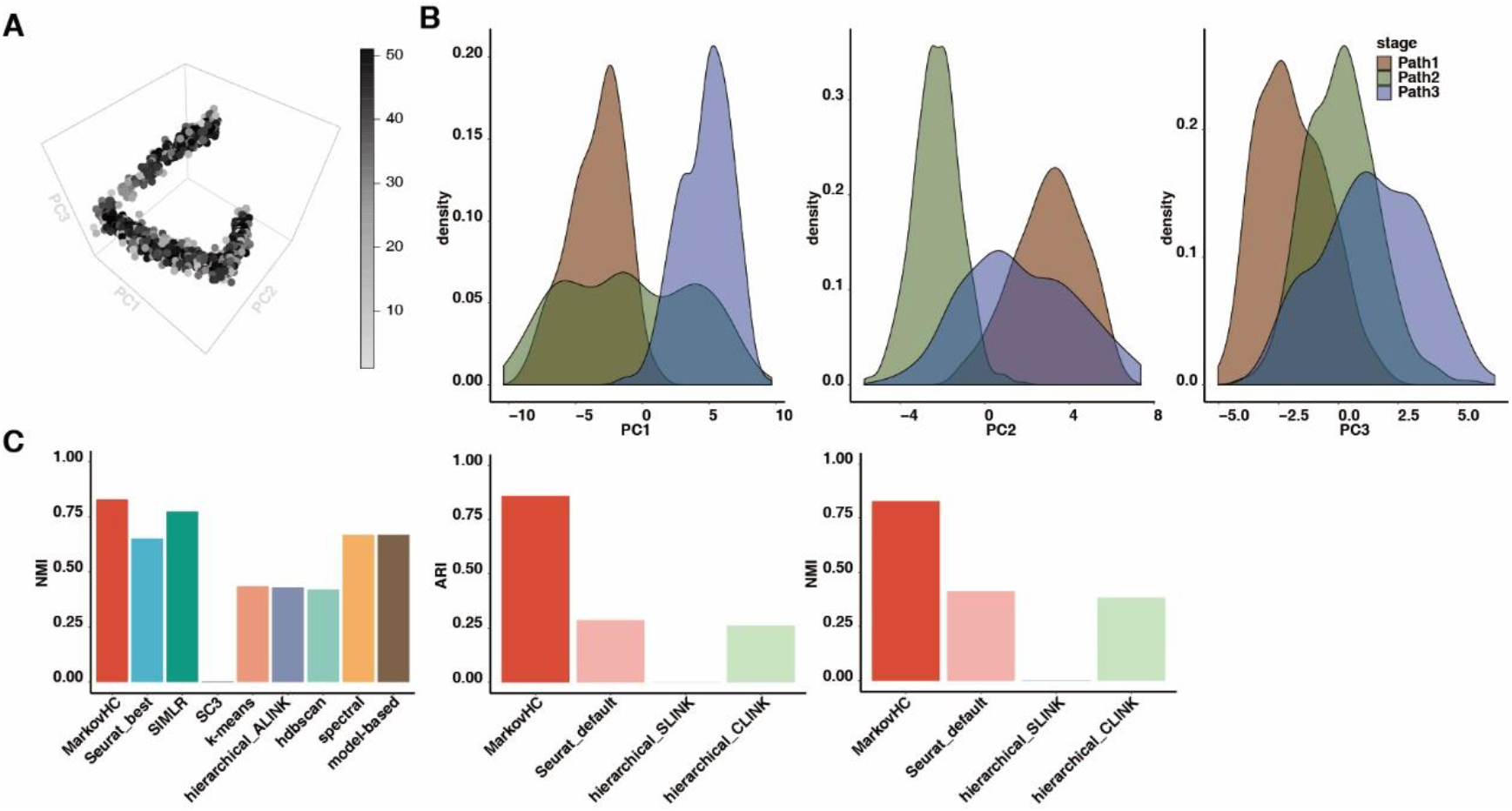
More results of the simulated single cell data with three stages in continuum. A. D scores of these points. B. The distribution of each stage on the first, second, and third principle component. C. Comparisons (NMI and ARI) with Seurat (with parameters fine-tuned), Seurat with default parameters, SIMLR, sc3, k-means, hierarchical clustering with different linkages, HDBSCAN, spectral clustering, and model-based clustering.

**Supplementary figure 4.**
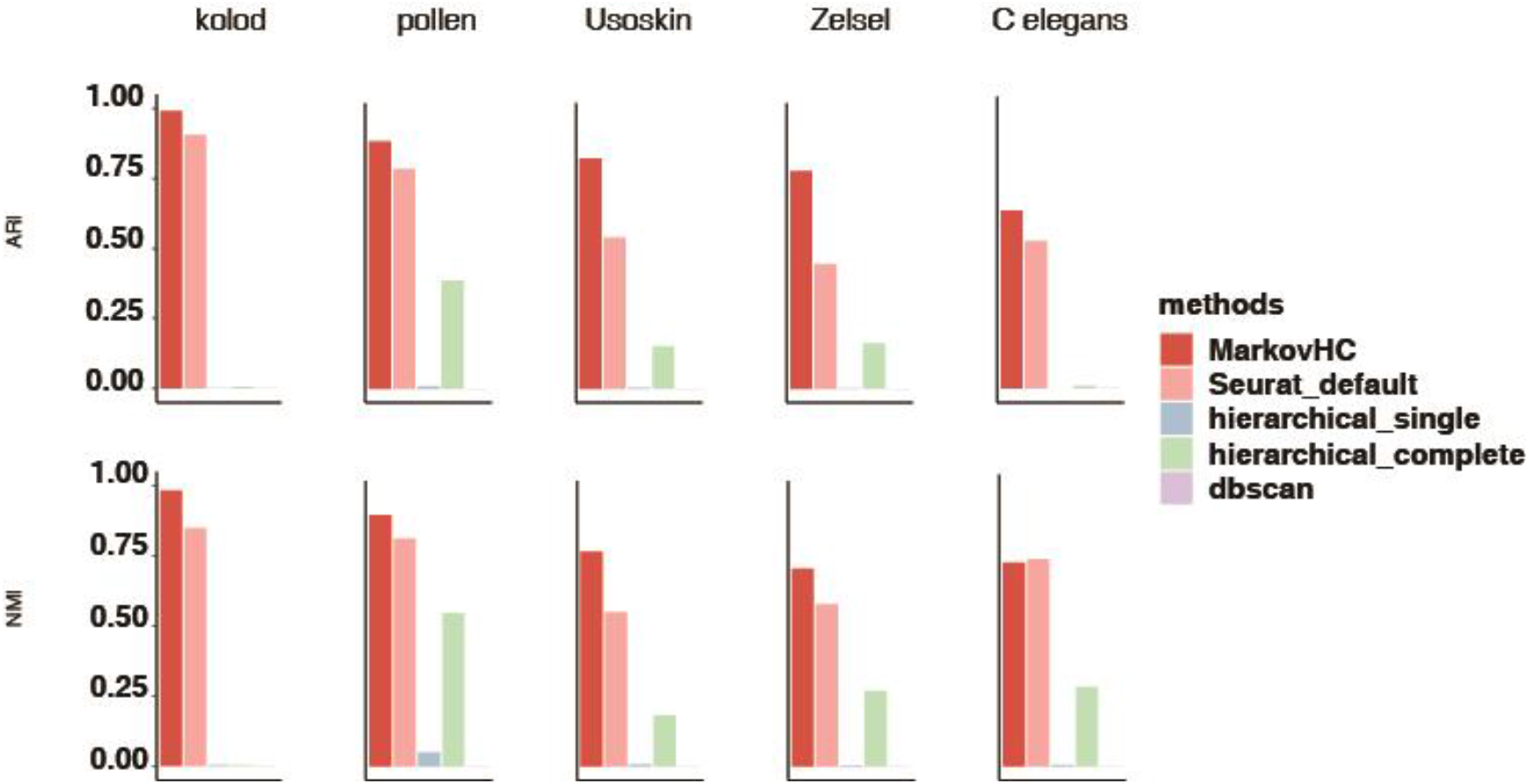
The comparison among MarkovHC, Seurat with default parameters and hierarchical clustering with other linkages. ARI and NMI show MarkovHC’s clustering accuracy is the best one on these data.

**Supplementary figure 5.**
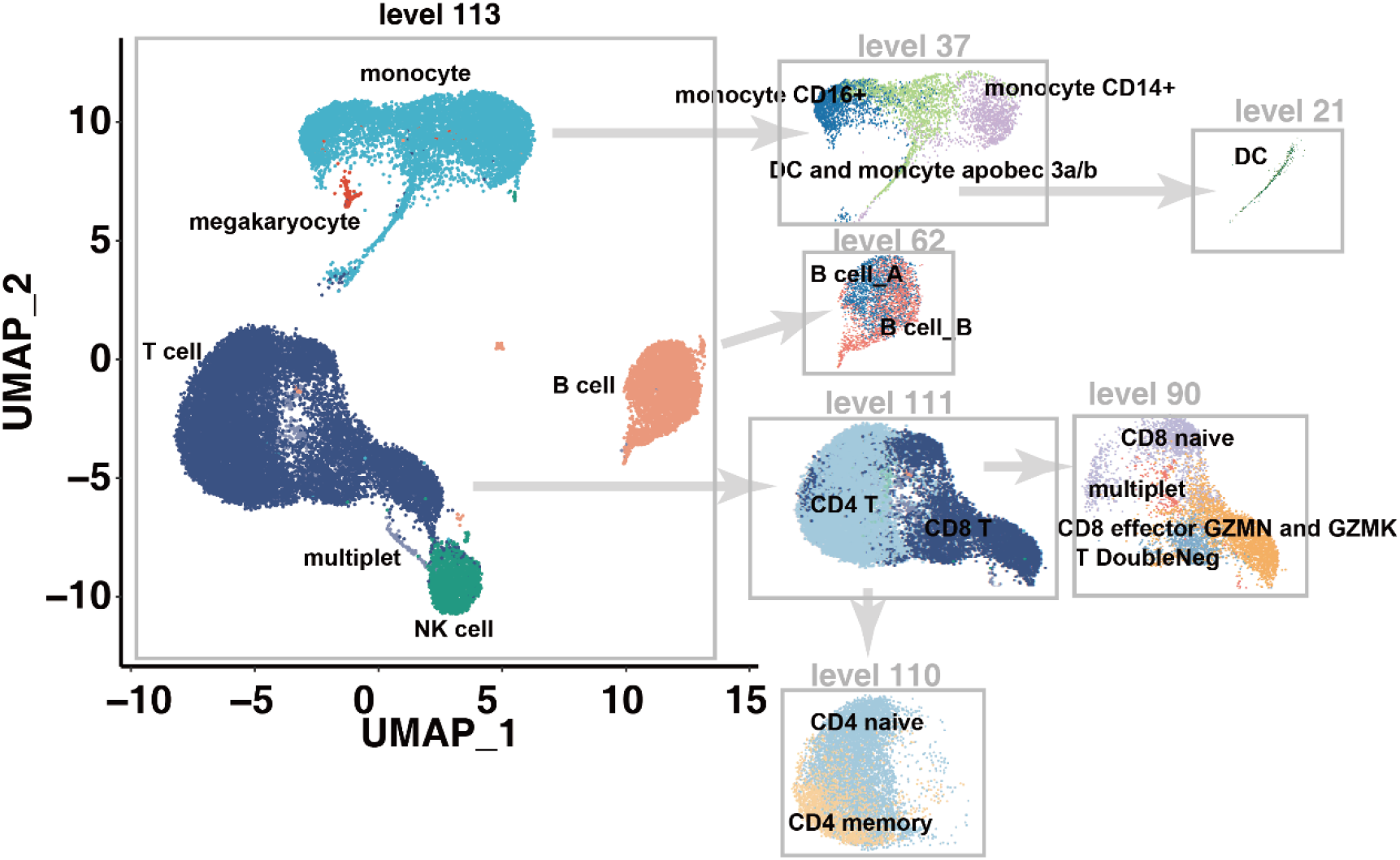
Basins in the 33k PBMCs dataset. On level 113, basins correspond to B cells, T cells, NK cells, megakaryocyte and monocyte could be identified. On level 111, the T cell basin was divided into CD4+ T cell basin and CD8+ T cell basin. On level 110, CD4+ T cell basin was further divided into CD4+ naïve and CD4+ memory T cell basins. Similarly, the subtypes of B cell, CD8+ T cell and monocyte could also be recovered on lower levels.

**Supplementary figure 6.**
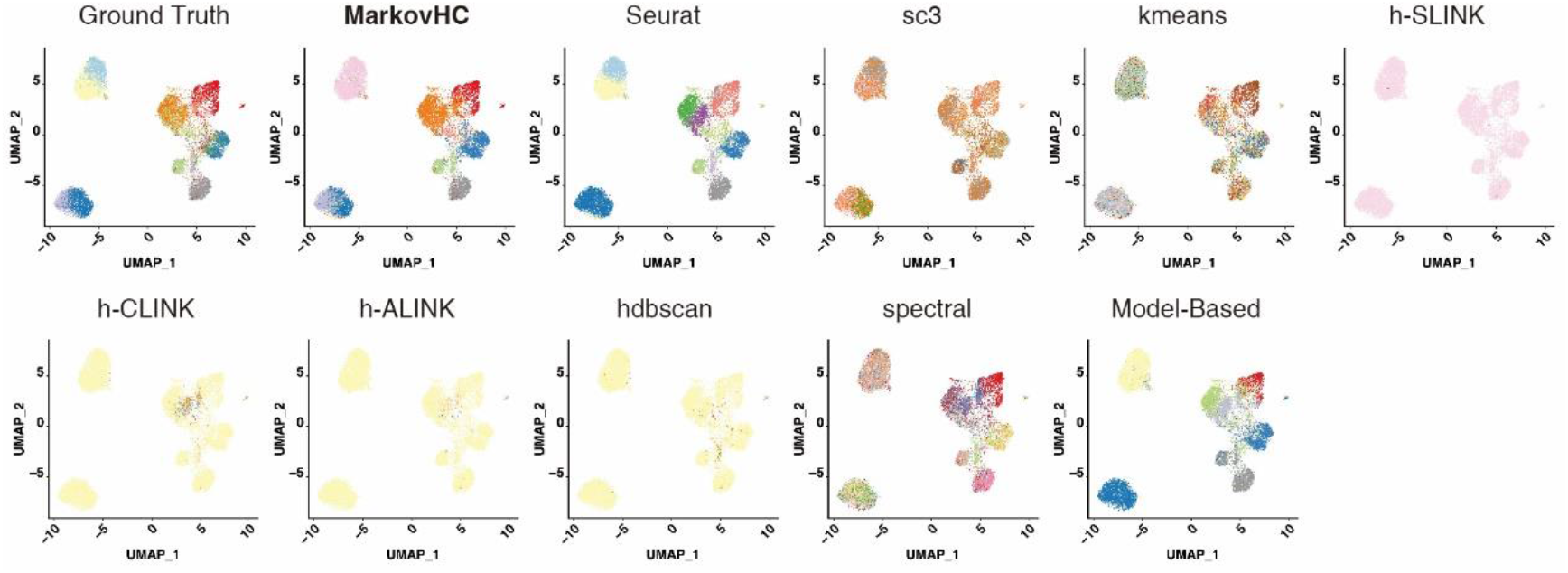
Clustering results of MarkovHC and other popular clustering methods. 10009 cells were projected these cells into 2-dimensional space by UMAP and colored according to the clustering results.

**Supplementary figure 7.**
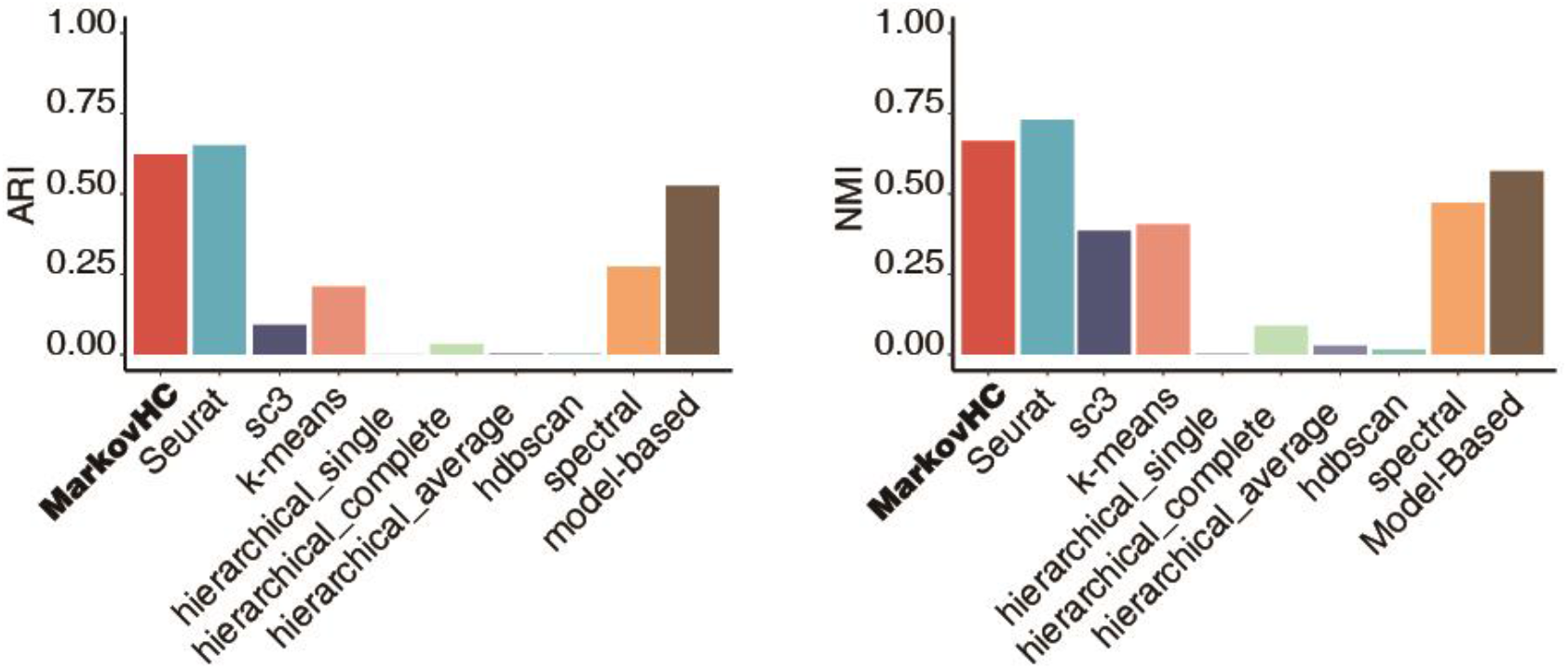
Compare MarkovHC’s clustering accuracy with those of other popular clustering methods on cytometry data. The results showed that MarkovHC got comparable performance to Seurat on this dataset. We noticed that sc3, k-means, simple hierarchical clustering methods and hdbscan couldn’t get good results when applied on this dataset. Spectral clustering and model-based clustering were good enough either. Meanwhile, we also run SIMLR on this dataset but the program was stuck several days for some reason. So we didn’t get its performance.

**Supplementary figure 8.**
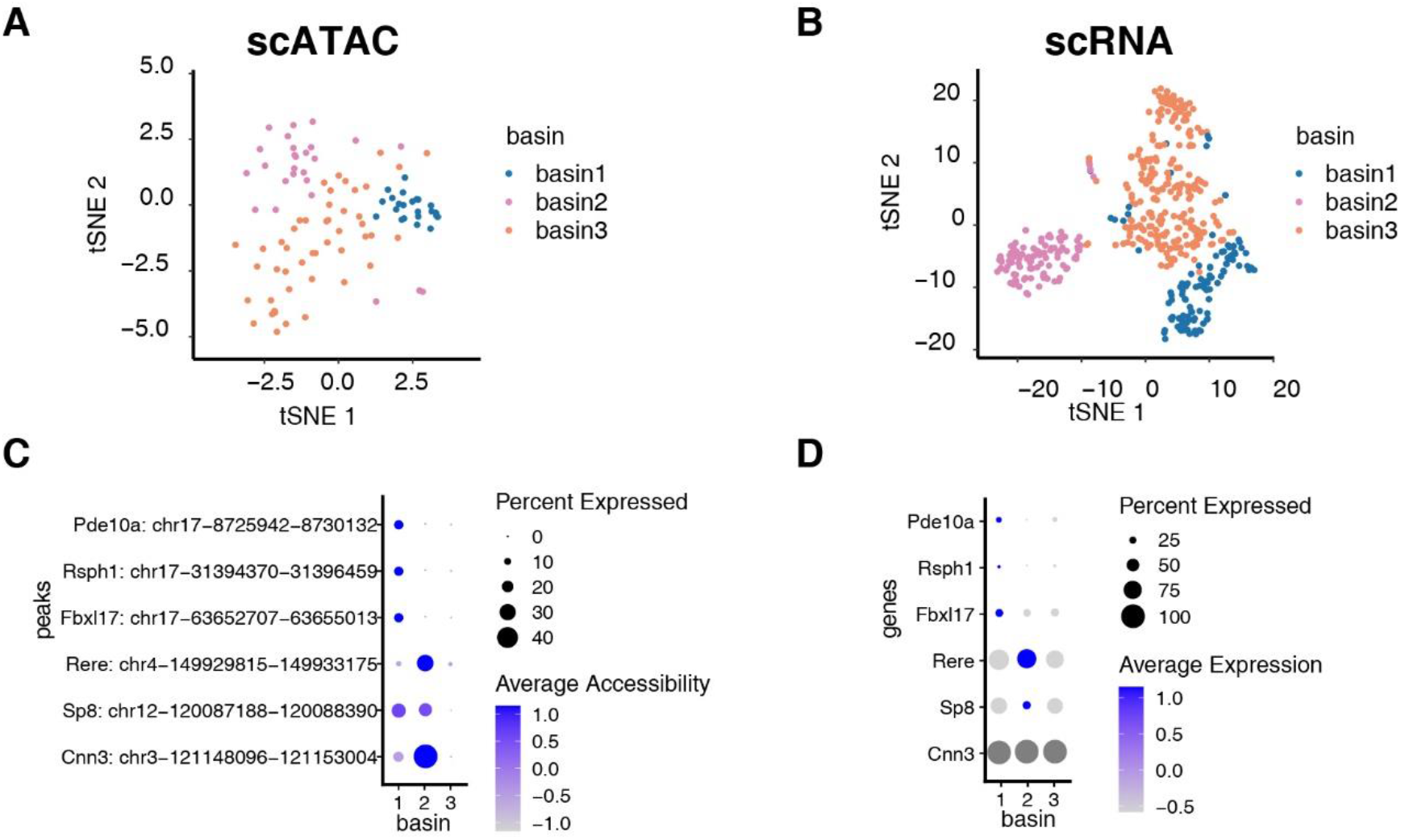
MarkovHC identified three basins in scATAC-Seq data and scRNA-Seq data.

**Supplementary figure 9.**
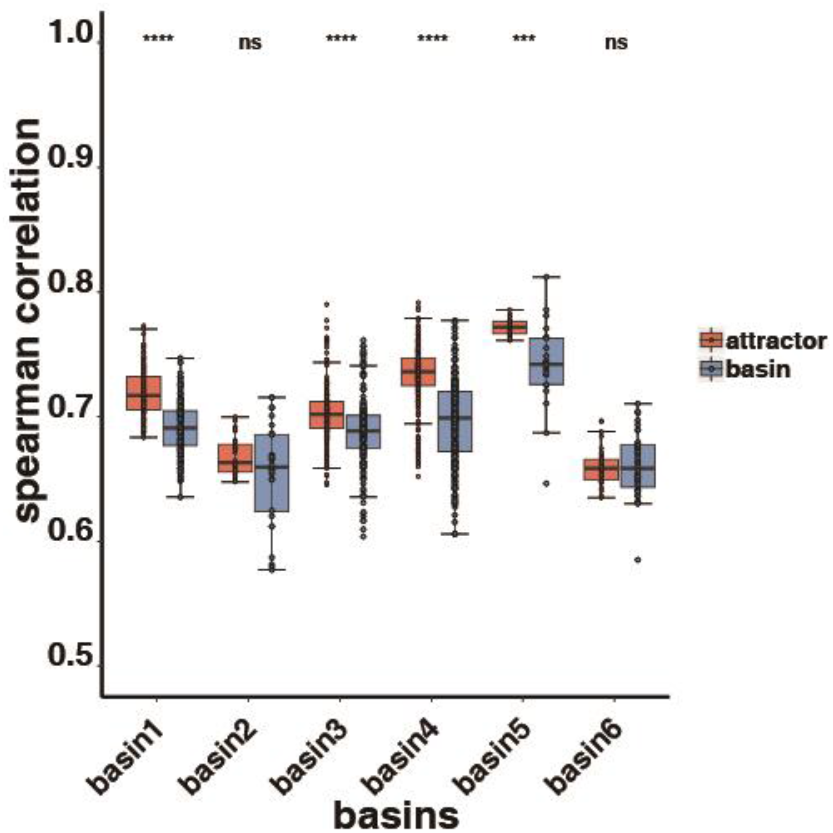
The spearman correlations among attractors and others in the basins. The spearman correlations of each cell pair in attractors are significantly higher than those of the same number of random selecting cell pairs in other basin cells which demonstrates that our hypothesis of the feature of attractors is correct in real biology data.

**Supplementary figure 10.**
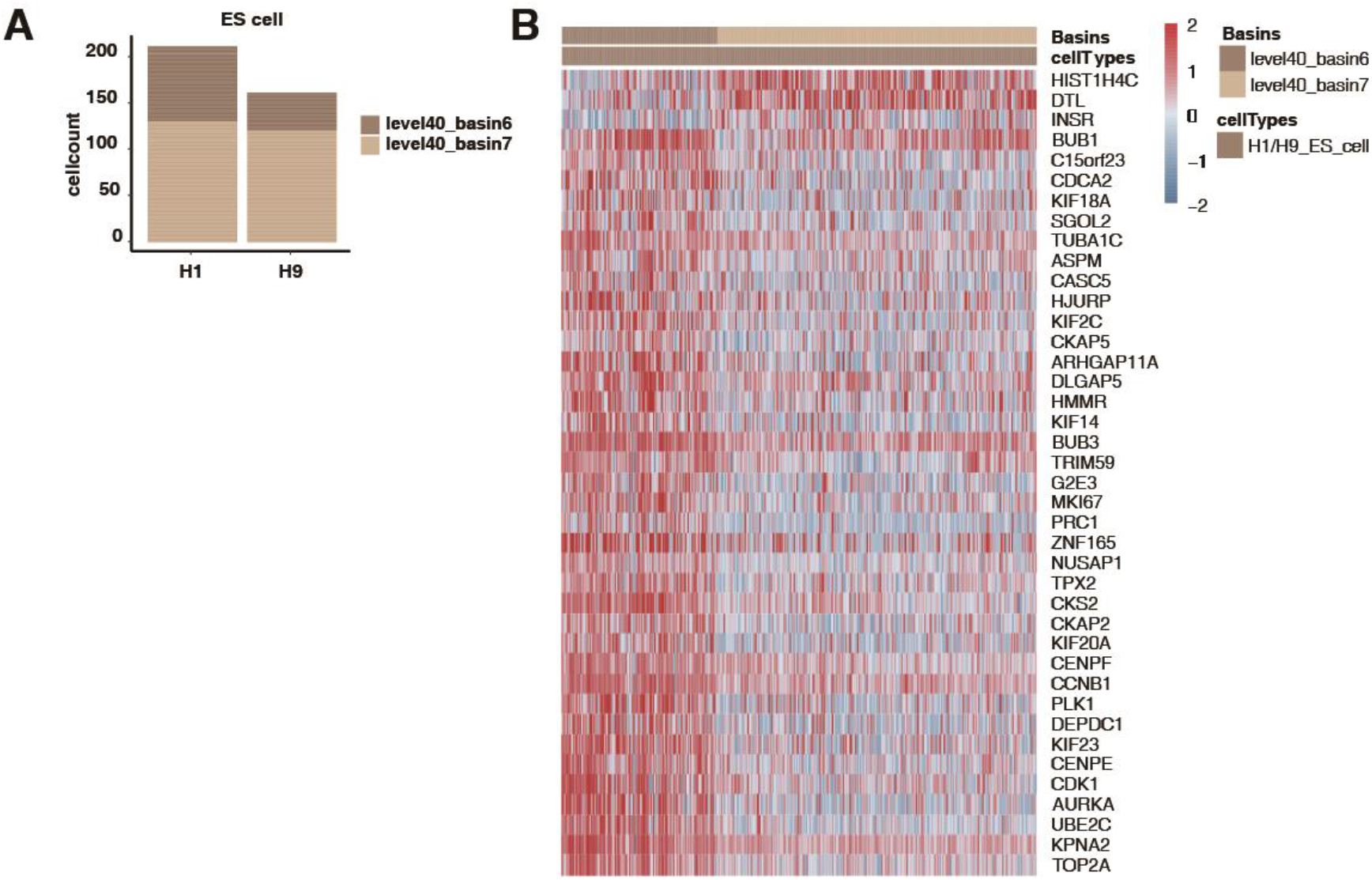
Two basins in the H1/H9 ES cells. A. The bar plot shows the proportion of these two basins in H1 and H9. B. The heatmap of differentially expressed genes between these two basins.

**Supplementary figure 11.**
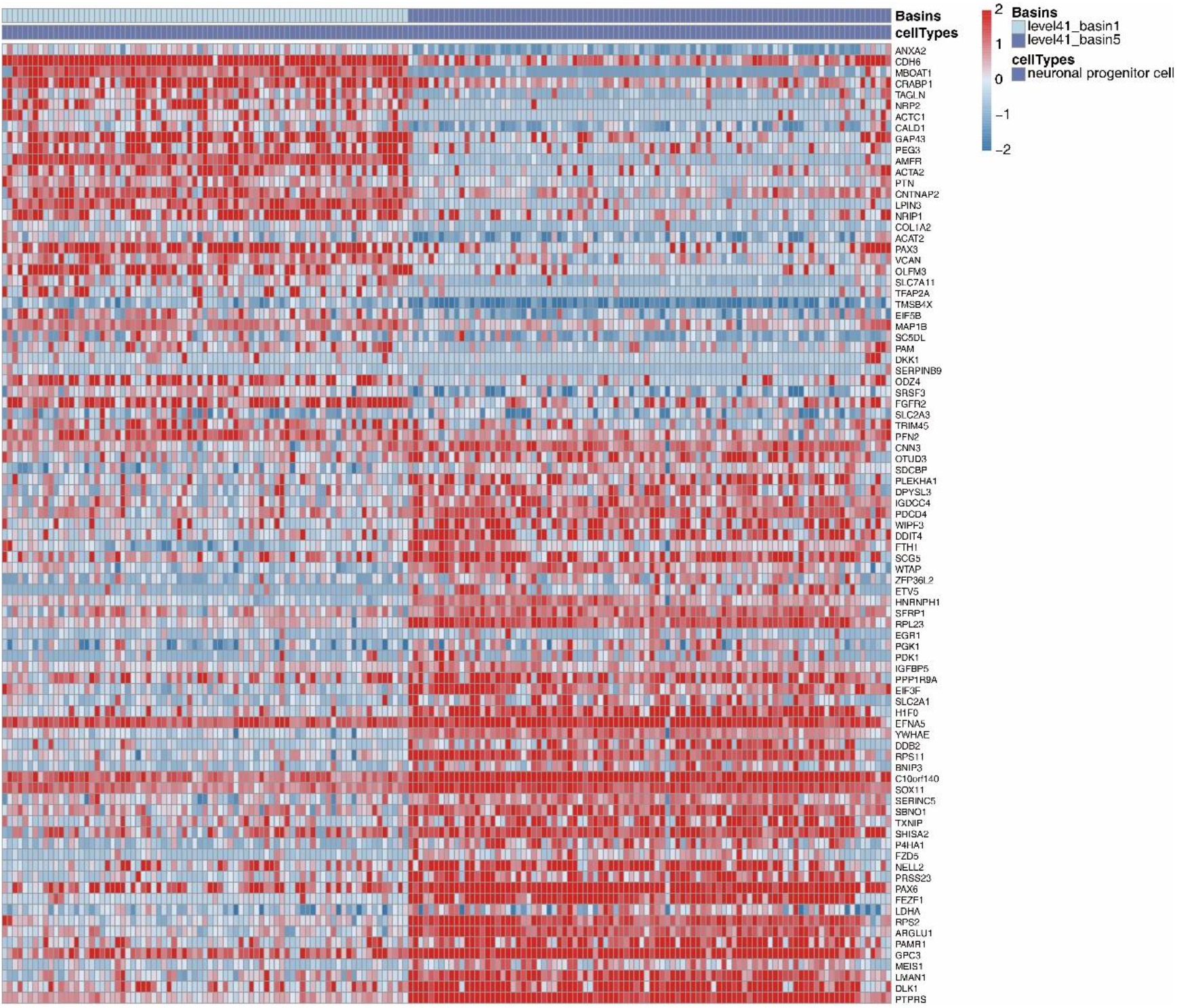
The heatmap of differentially expressed genes between two basins in neuronal progenitor cells.

**Supplementary figure 12.**
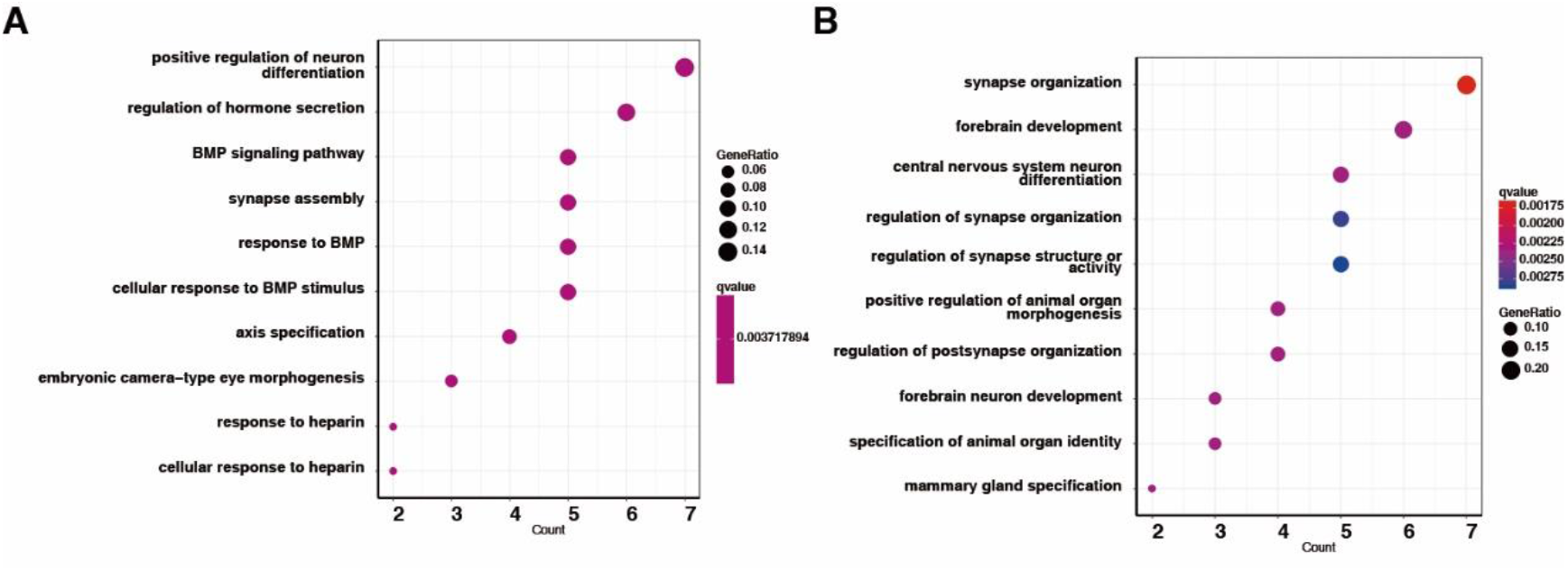
GO terms of up-regulated genes in two subtypes in neuronal progenitor cells.

**Supplementary figure 13.**
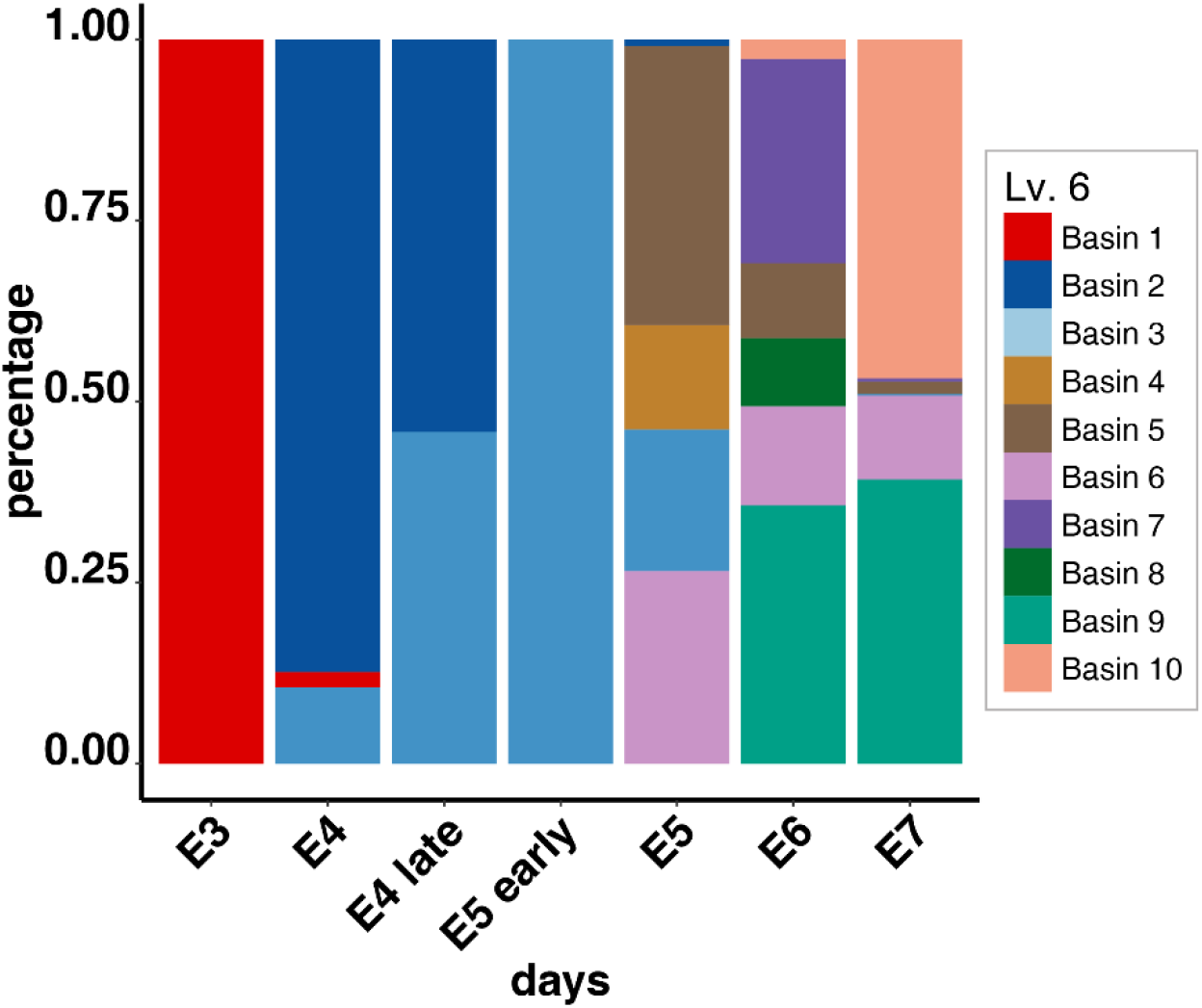
The proportion of basins in each time point.

**Supplementary figure 14.**
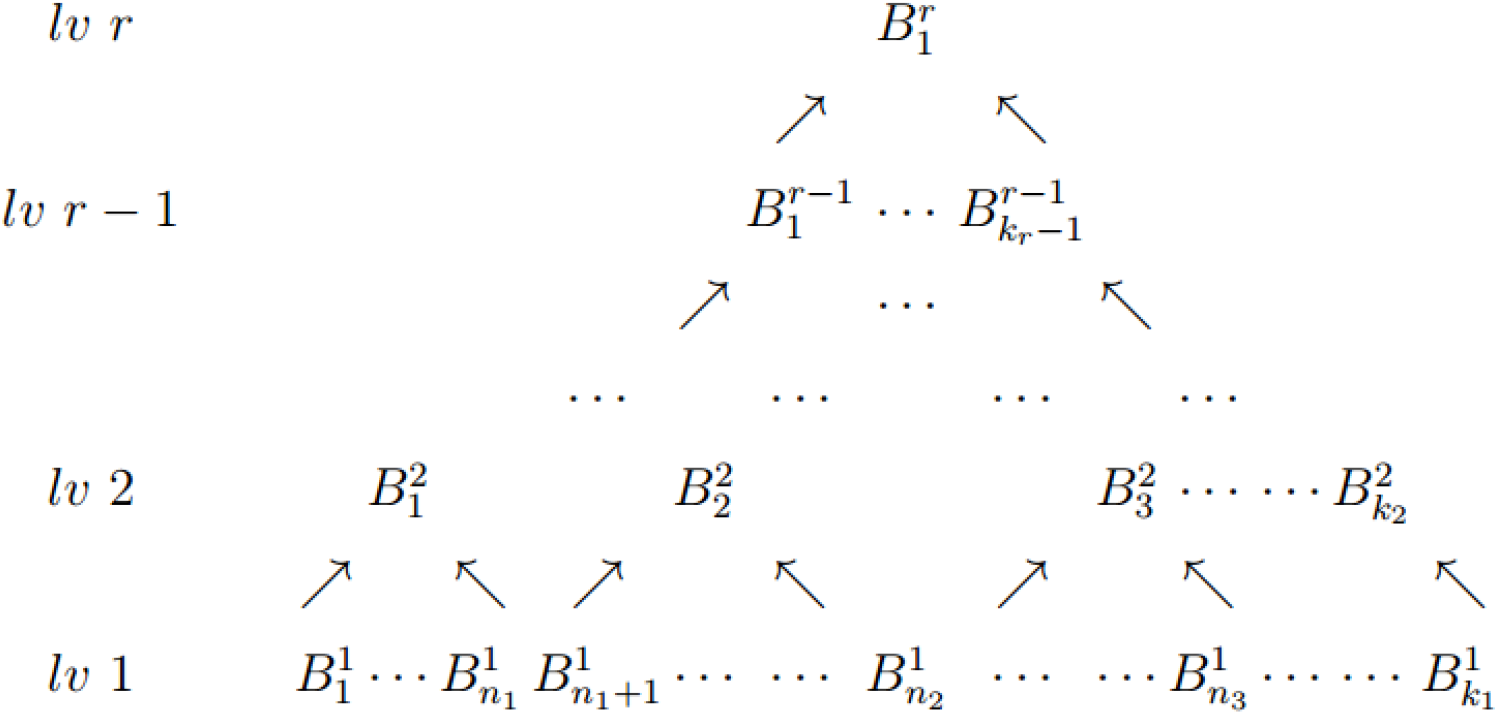
The target hierarchical structure of MarkovHC. 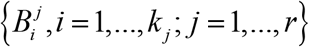 is the subset of {*s*_1_,…,*s_n_*} and 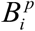 is the subset of some basins on level p+1.

**Supplementary figure 15.**
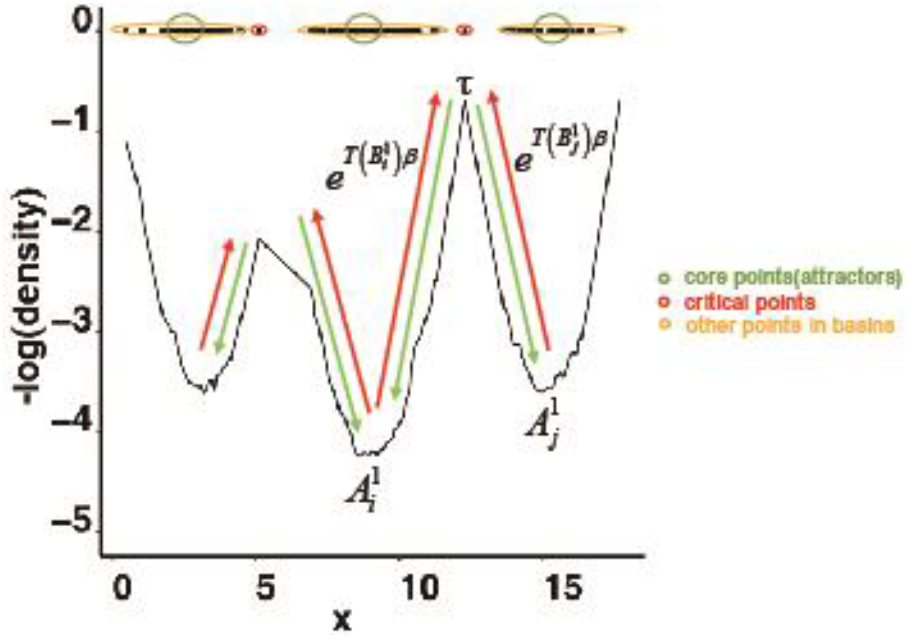
1-dimensional example of landscape. The points are 1-dimensional and they are plotted at the top of the panel. The value of vertical coordinate is the pseudo potential energy, which is defined as the negative logarithm of density here. The time scale for points in 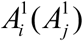 to reach τ is 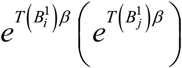 while the time scale for τ to reach 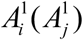 is a constant irrelative with β. (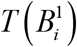 can be viewed as the least consumption of pseudo potential power for points in 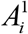 to reach τ.) Therefore, it takes a long time for points in attractors to move to shared points while shared points can jump into attractors in a very short time. Of course, the time scales for points in 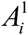 and 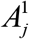 to move to τ are different. Such a difference is exactly the criterion used to define the relationship between level 1 clusters (basins) and to construct higher-level Markov chains.

**Supplementary figure 16.**
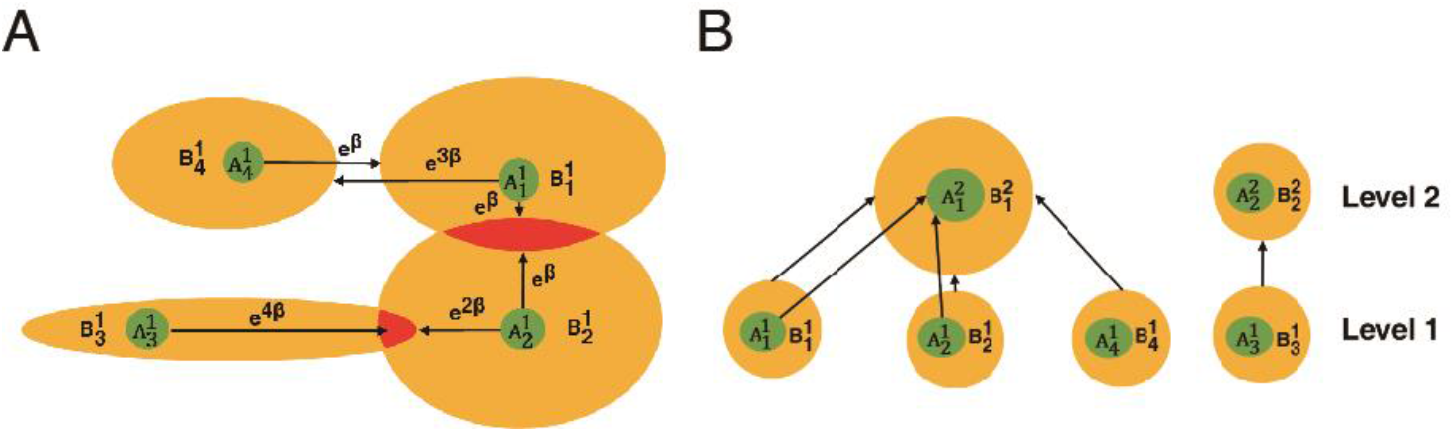
The illustration of the criteria for basin merging. A. Four basins on level 1. B. Basing on our criteria, 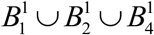 is a new level 2 basin.

**Supplementary figure 17.**
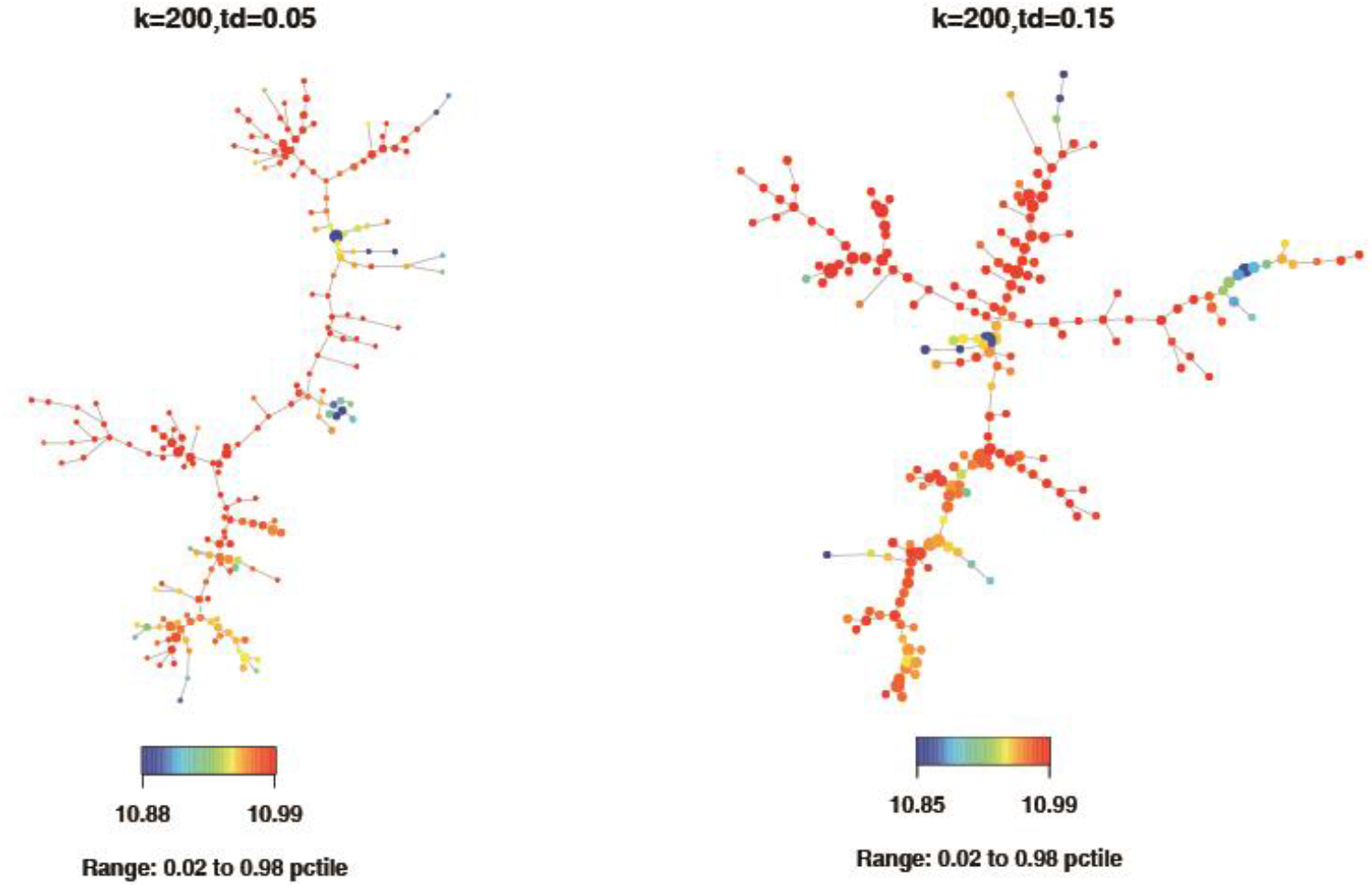
Minimum spanning trees constructed by SPADE is not robust. We applied spade on the cytometry data of PBMC twice, and only changed one of insignificant parameters, downsampling_target_pctile, in the algorithm. The structures of MST (minimum spanning tree) were very different. For the stochastic process in the algorithm, even we ran it twice with same parameters, the MSTs were different either.

**Supplementary figure 18.**
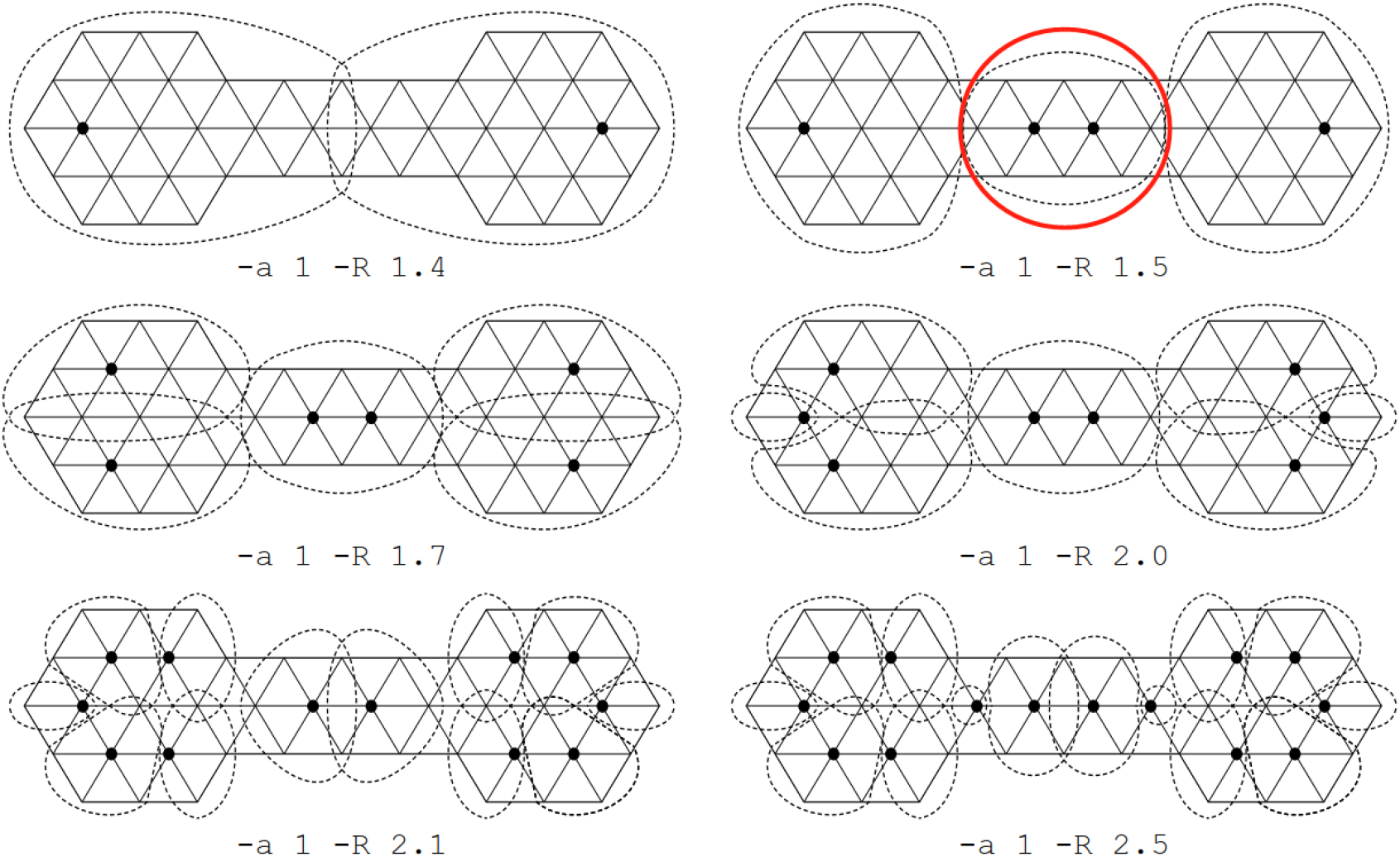
Clusterings resulting from the MCL algorithm for the graph in Van Dongen’s Ph.D. thesis.

**Supplementary figure 19.**
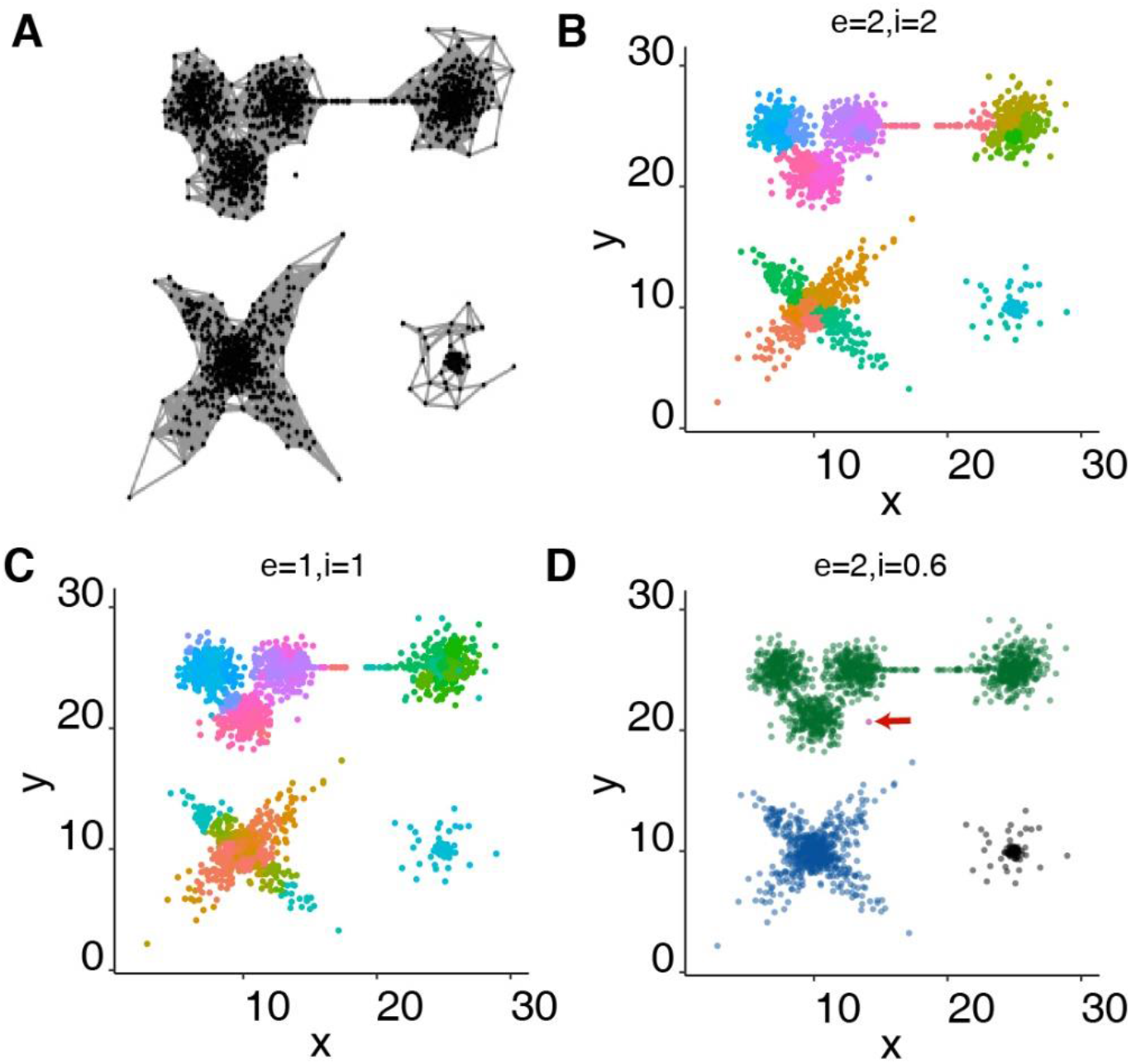
Clustering results of MCL on simulation data in Fig.2 F.

**Supplementary figure 20.**
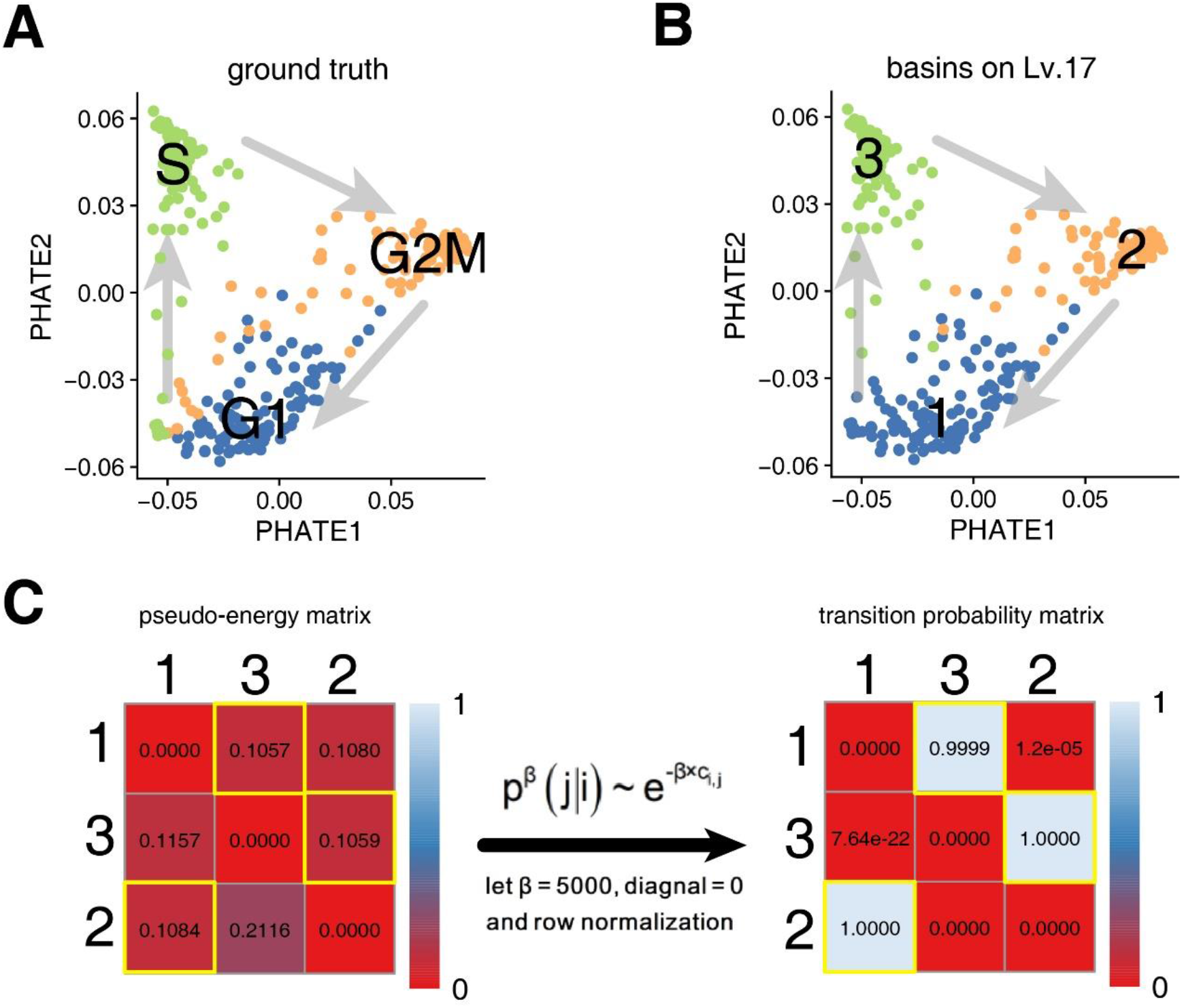
MarkovHC can reveal non-equilibrium biological processes.

**Supplementary figure 21.**
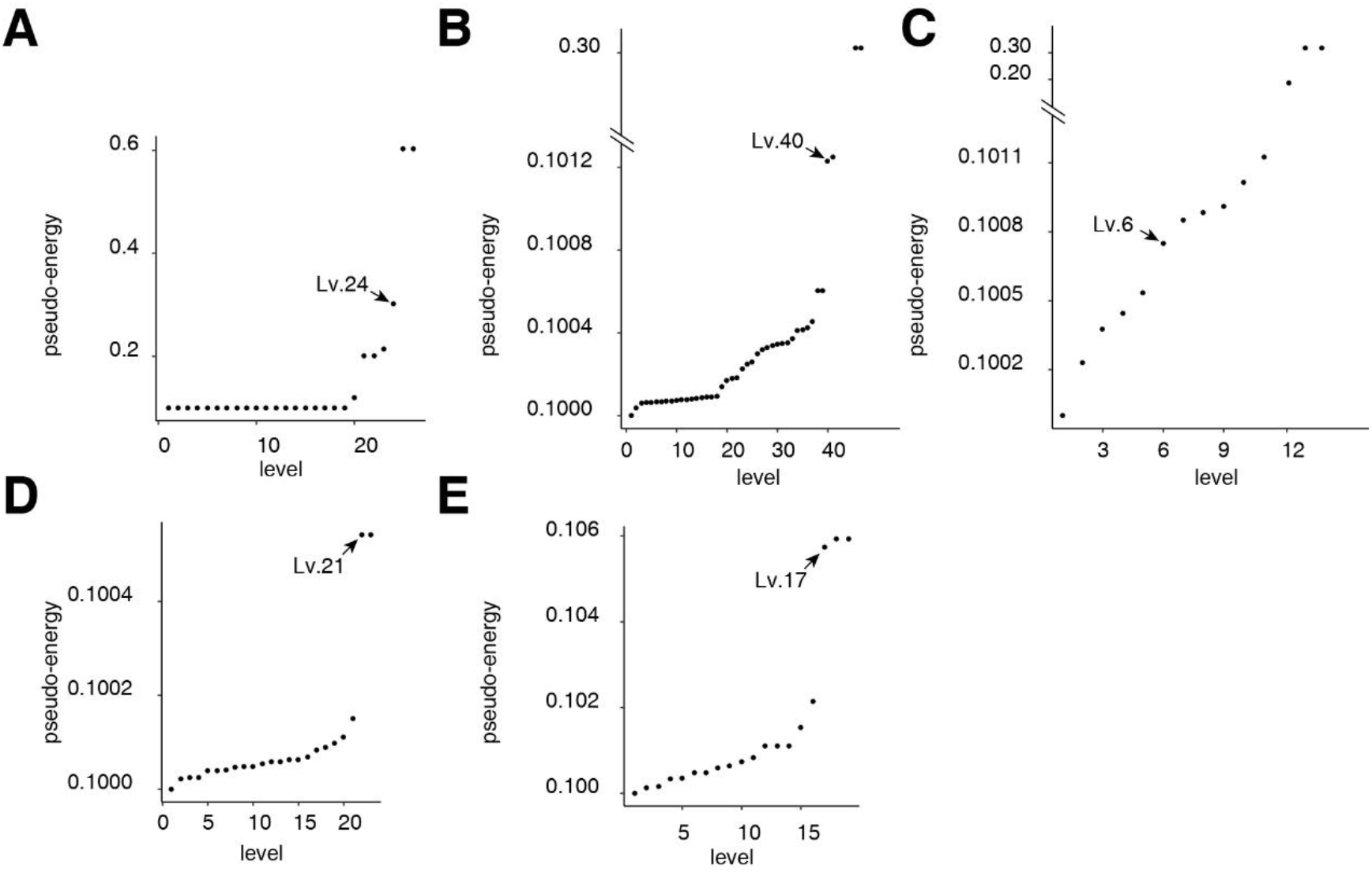
Level selection strategy.

**Supplementary figure 22.**
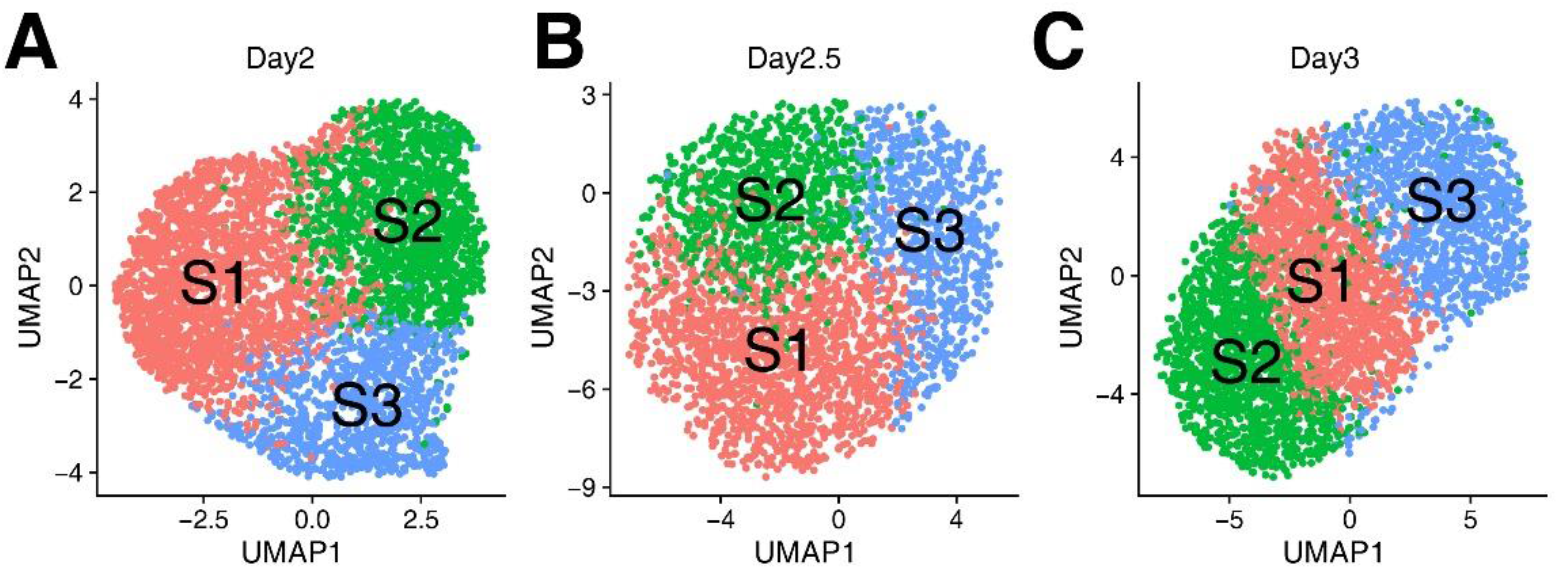
The sub-clusters in Day2, Day2.5 (the night in Day2), and Day3.

**Supplementary figure 23.**
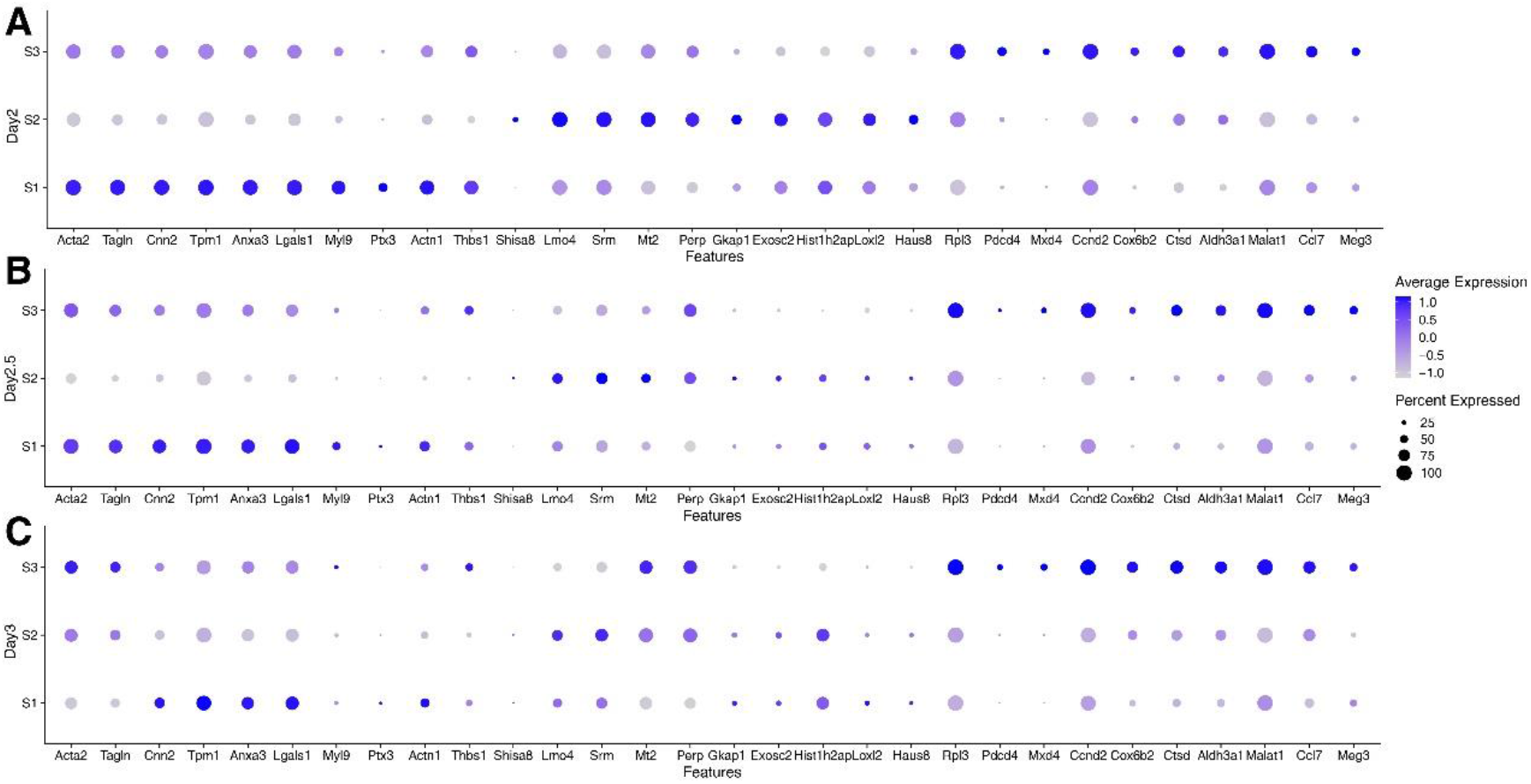
The pattern of differentially expressed genes.

**Supplementary figure 24.**
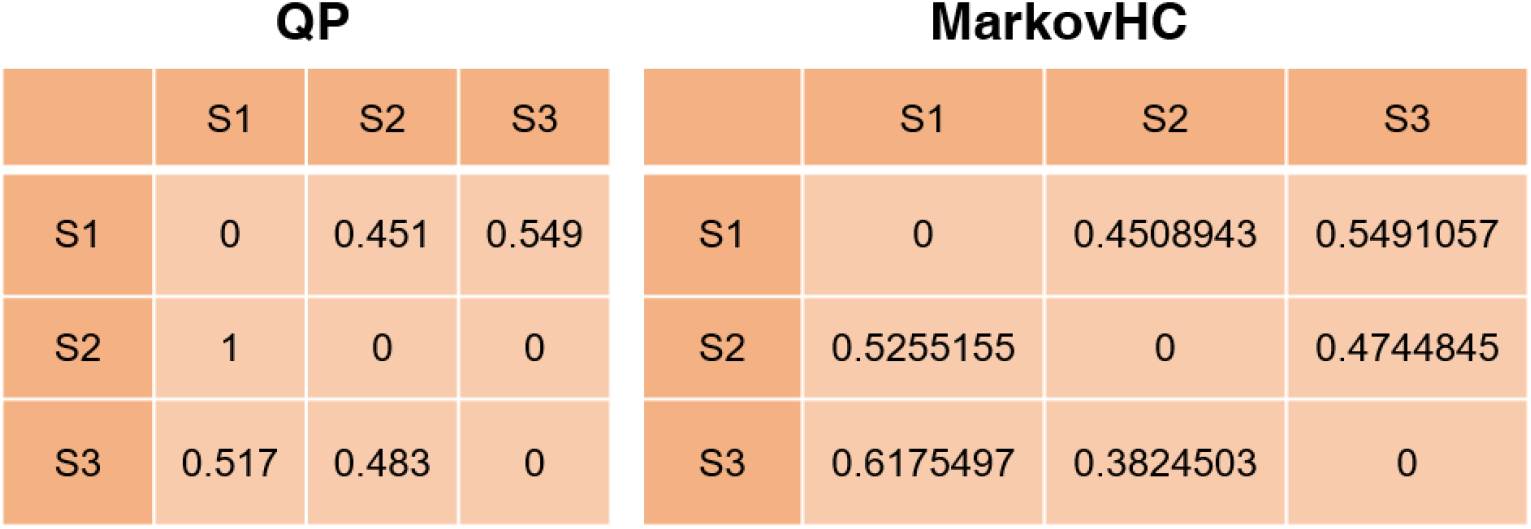
P_ij_ matrix calculated by QP (left) and MarkovHC (right)

**Supplementary figure 25.**
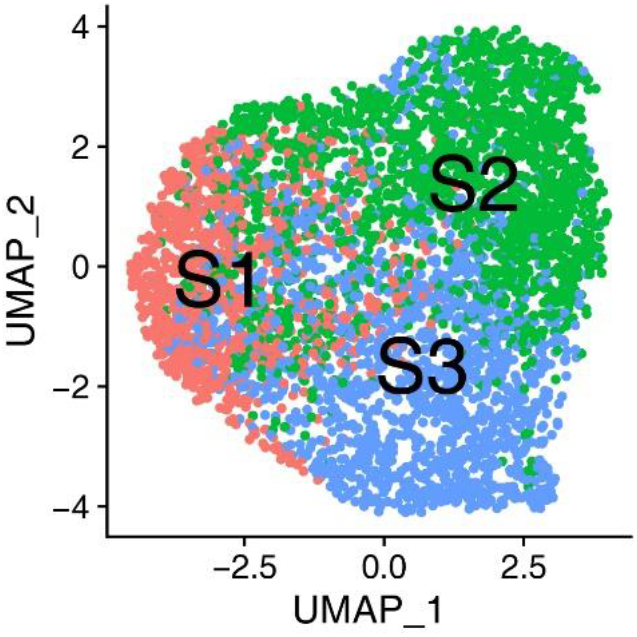
Three basins clustered by MarkovHC (right)

